# Capturing species-wide diversity of the gut microbiota and its relationship with genomic variation in the critically endangered kākāpō

**DOI:** 10.1101/2022.10.31.514450

**Authors:** Annie G. West, Andrew Digby, Anna W. Santure, Joseph G. Guhlin, Peter Dearden, Kākāpō Recovery Team, Michael W. Taylor, Lara Urban

## Abstract

The gut microbiota plays an essential role in host health that has important implications for the conservation management of threatened wildlife. While factors such as diet, medication, and habitat are known to shape the microbiota, our understanding of the entirety of factors, including the complex role of the host genomic background, remains incomplete. Our research on the gut microbiota of the critically endangered kākāpō (*Strigops habroptilus*), a flightless parrot species endemic to Aotearoa New Zealand, represents, to our knowledge, the first study to describe the gastrointestinal bacterial diversity for virtually an entire species and to assess the relationship between gut microbiota and host genomic diversity in a highly threatened population. Here we report a 16S rRNA gene-based analysis of kākāpō faecal samples representing the gut microbiota for 84% of kākāpō (n = 133). This survey was then leveraged with exceptional metadata to tease apart the impact of host genomic diversity and factors such as sex, diet, antibiotic treatment, disease status, habitat, and time of sampling on the kākāpō gut microbiota, with sex being the only covariate significantly associated with gut microbiota diversity. We find evidence of a highly polygenic genomic architecture of the gut microbiota and further identify putative associations between gut bacterial diversity and functional biological pathways related to intestinal homeostasis, inflammation, immune response and metabolism. This improved understanding of the kākāpō gut microbiota – and its relationship with host genomics – can directly benefit kākāpō management and conservation by providing new insights into the role of the gut microbiome in kākāpō health and disease mitigation. Overall, we anticipate that an integration of microbiome studies in conservation research and management will improve our understanding of how the concept of One Health with its implications for human, animal and environmental welfare can be achieved.

## Background

The microbiota is intrinsic to animal health, fitness and development [Cho and Blaser 2012], with essential roles in digestion, homeostasis, immune system regulation, reproduction and neurological processes [Spor et al. 2011; Gilbert et al. 2018]. The potential influence of the gut microbiota on the health and fitness of threatened wildlife is pertinent to conservation management programmes, especially when it comes to captive or highly managed populations [Trevelline et al. 2019; San Juan et al. 2021; West et al. 2019, 2022a, 2022b]. For example, fertility in both the critically endangered eastern black rhinoceros and threatened southern white rhinoceros was directly related to microbiota-mediated hormone production and metabolism [Antwis et al. 2019; Williams et al. 2019], enabling the identification of potential biomarkers of reproductive health to aid species conservation.

While the microbiota can be shaped by environmental factors such as diet, medication, habitat or social networks [David et al. 2014; Rodrigues Hoffmann, 2017; Rothschild et al., 2018; San Juan et al. 2021; West et al. 2022a, 2022b], our understanding of the entirety of factors that shape the microbiota remains incomplete. Microbiota composition can, for example, be significantly associated with phylogeny [Waite and Taylor 2014; Rojas et al. 2021], and can override substantial divergence in a species’ ecology (e.g., the gut microbiota of the herbivorous giant panda is similar to those of its carnivorous sister species [Xue et al. 2015]). One notable exception to this observation is among bats and avians, where the gut microbiota is strongly associated with volancy over phylogeny [Song et al. 2020]. Host genetic background can also have substantial implications for microbiota composition [Zhernakova et al. 2017], though its role remains unexplored in most species given the need for large cohorts for which both genomic and microbiota data are available. While such datasets have recently been analysed for humans [Goodrich et al. 2016; Hall et al. 2017; Awany et al. 2019; Ishida et al. 2020], to our knowledge a study on the relationship between the microbiota and host genome has not yet been undertaken for any threatened wildlife. Such information should help inform future efforts at microbiome engineering of threatened species (e.g. West et al. 2019) by teasing apart the influences of host genetics and the - potentially more manipulable - effects of environmental factors.

Here, we created a gut microbiota catalogue for virtually an entire critically endangered species, the kākāpō (*Strigops habroptilus*). We then combined our microbiota catalogue with extensive environmental data and genomic data for all individuals [Ghulin et al. 2022] to obtain a full picture of the factors that influence the microbiota diversity of an entire threatened species. The kākāpō is a flightless and nocturnal parrot endemic to Aotearoa New Zealand that was considered functionally extinct by the mid-20th century due to habitat fragmentation and invasive mammalian predators. The extant kākāpō population primarily descends from a single population discovered on Rakiura / Stewart Island, Aotearoa New Zealand, and from one male individual, Richard Henry, who was the only surviving ‘mainland’ kākāpō to successfully father offspring. Having undergone severe population bottlenecks, kākāpō are now highly inbred and suffer from low reproductive success, which is further compounded by their sporadic breeding which only occurs during heavy podocarp mast seasons (approx. every 2 - 4 years) [Merton et al. 1984; Powlesland et al. 1992; Elliott et al. 2001; Eason et al. 2006; Savage et al. 2021]. However, recent findings suggest that despite their long-term small population size, kākāpō have a reduced number of harmful mutations than expected [Dussex et al. 2021]. As a species, the kākāpō has further suffered from increased disease susceptibility, especially to exudative cloacitis, an inflammatory cloacal condition with unclear pathogenesis that significantly impacts kākāpō health [Jakob-Hoff et al. 2009, White et al. 2015]. The surviving ~200 kākāpō now live as wild populations on five predator-free offshore islands and are managed by the Kākāpō Recovery Programme of the New Zealand Department of Conservation (Te Papa Atawhai). The kākāpō is a unique bird, being the largest, the only flightless and the only lek-breeding parrot in the world [Williams, 1956; Merton et al. 1984; Powlesland et al. 2006], in addition to a strictly herbivorous diet [Best, 1984; Atkinson and Merton, 2006; Powlesland et al. 2006]. It is therefore likely that the kākāpō gut microbiota plays an important role in digestion and detoxifying plant chemicals, similar to other herbivorous species [Mackie, 2002; Amato et al. 2013, 2016]. Previous research has shown that the kākāpō gut harbours a low-diversity bacterial community that is frequently dominated by a single species of *Escherichia-Shigella*, which is highly unusual for a herbivorous animal [Waite et al. 2012, 2014, 2018; Perry et al. 2017; West et al. 2022b]. This microbiota is robust to anthropogenic influence such as supplemental feeding or hand-rearing, biogeographic variation, and individual characteristics such as sex or age [Waite et al. 2012, 2014, 2018; Perry et al. 2017].

We first produced and collated bacterial 16S rRNA gene data from faecal samples (total n = 133) for nearly the entire extant adult kākāpō population (n = 141 population size as of 31/12/2020, excluding chicks hatched in the 2019 breeding season) and several deceased individuals (n = 14), and subsequently combined it with whole-genome sequence data for the same individuals (produced under the Kākāpō125+ genome project) [Ghulin et al. 2022; Kākāpō Recovery Team]. We then investigated associations between bacterial diversity and host characteristics, including kākāpō sex, antibiotic treatment, supplemental feeding, disease status (including exudative cloacitis) and hand-rearing, as well as environmental factors such as island location and the year and season of sample collection. We leveraged this information and the genomic dataset to assess heritability, genetic architecture and genome-wide associations of gut bacterial diversity, *Escherichia-Shigella* dominance and core bacterial taxa abundance. We hope that an improved understanding of the kākāpō gut microbiota and its relationship with the host genomic background can directly benefit kākāpō management and conservation by providing new insights into the role of the microbiota in kākāpō health and fitness, following the concept of One Health which affirms the interconnectivity of all living systems and their health.

## Materials and Methods

### Sample collection

This study is based on the faecal microbiota of 133 kākāpō individuals for which (a) we collected fresh faecal material (n = 74), (b) faecal material had already been stored at −20°C from previous sampling efforts (n = 29), or (c) 16S rRNA gene amplicon sequencing data were available (n = 30) from Perry et al. [2017]. This dataset altogether contains microbiota data for 84% of the extant adult kākāpō population (119 of 141 surviving adult kākāpō as of 31/12/2020) and for 14 historical and deceased individuals.

Faecal samples were obtained from six offshore islands (Figure 1): Whenua Hou (Codfish Island, 46° 47’ S, 167° 38’ E), Pukenui (Anchor Island, 45° 45’ S, 166° 31’ E), Te Hauturu-o-Toi (Little Barrier Island, 36° 11’ S, 175° 04’ E), Te Kākahu-o-Tamatea (Chalky Island, 46° 03’ S, 166° 31’ E), Pearl Island (47° 11’ S, 167° 42’ E), and Te Pākeka (Maud Island, 41° 02’ S, 173° 53’ E). Samples from Pearl Island and Te Pākeka were collected in 1998 and 1999 (historical samples). As samples were collected opportunistically during routine monitoring, fresh faecal material was not always available. Cloacal swabs were initially collected if fresh faecal material could not be obtained, but the bacterial profiles of swabs were sufficiently dissimilar to those of faecal material that they could not be used as a proxy (Supplementary Figure 1). Where fresh faecal material was available, a portion was placed directly into a sterile polypropylene tube containing ~2.5 mL of RNA*later* solution, stored at 4°C overnight and subsequently stored at −20°C until samples could be transported off the island for processing. Where fresh faecal material could not be collected and the kākāpō individual had not already been included in the Perry et al. [2017] study, we identified faecal samples being stored at −20°C and collected as aseptically as possible during previous island excursions. These previously-collected samples were either stored immediately in RNA*later* or frozen on the island as soon as possible following collection, ranging from 1 to 4 h post-defecation. Faecal material stored in RNA*later* was shipped on ice, and frozen samples on dry ice, to Waipapa Taumata Rau The University of Auckland for DNA extraction. Accompanying metadata (Additional File 2) for all samples include location and date of sample collection, antibiotic history, exudative cloacitis history, sex of the kākāpō individuals, if supplemental feed was provided in the three months prior to sample collection and if the kākāpō were hand-reared as chicks.

**Figure 1:**
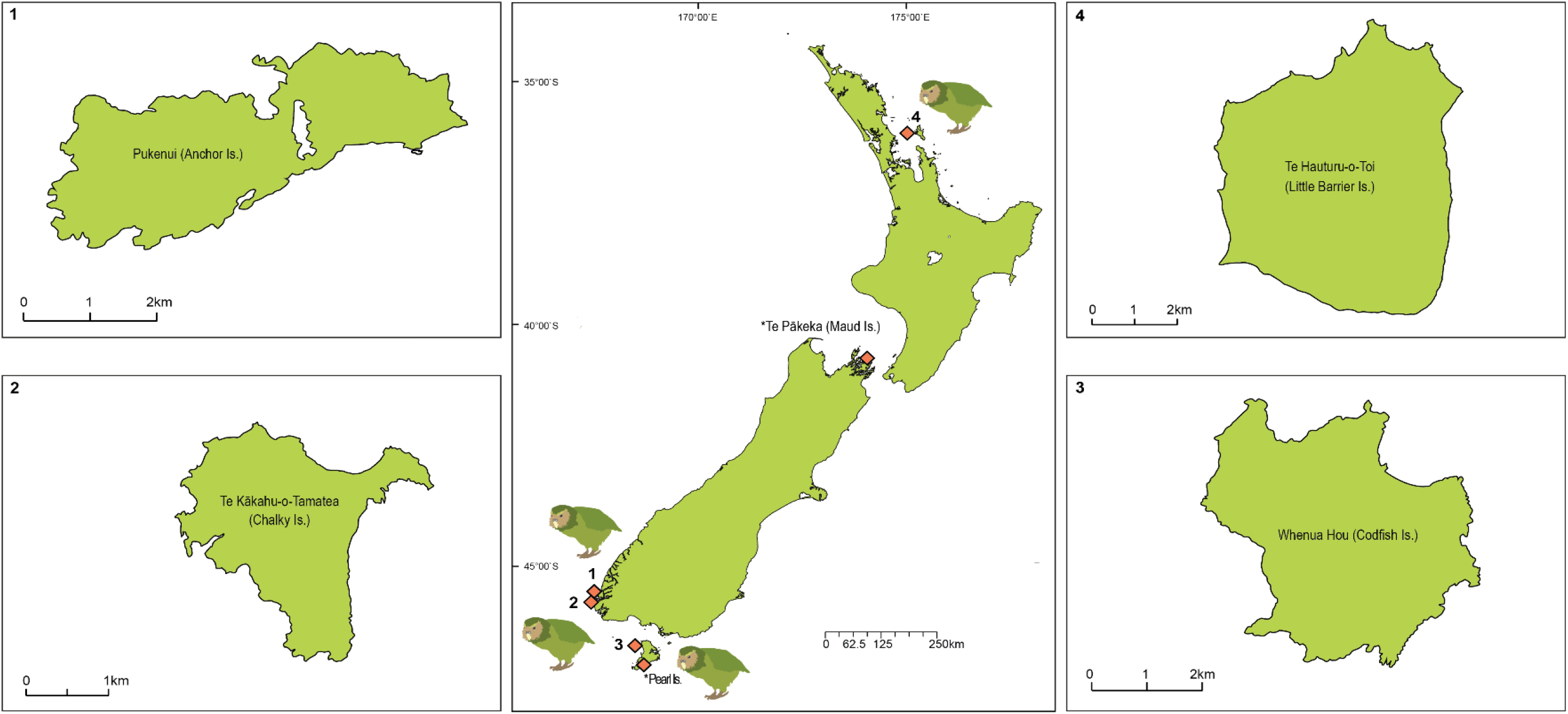
Map of Aotearoa New Zealand depicting the offshore islands where kākāpō faecal samples were collected. Historical samples were collected from *Te Pākeka (Maud Island) and *Pearl Island.

### DNA extraction

DNA was extracted from thawed faecal samples using a modified version of a CTAB-based bead-beating method previously described by Perry et al. [2017] that had been slightly altered to reduce both the presence of abundant PCR inhibitors and co-precipitation of excess salts. For each sample, 100 mg of faecal material was washed initially with PBS if stored in RNA*later*, or with 70% ethanol if directly frozen. Samples were then washed a second time with 70% ethanol before suspension in a high-salt CTAB extraction buffer with 30 mg of PVPP. Samples processed prior to mid-2019 were agitated at 5.5 ms^−1^ for 30 s using a FastPrep FP120 bead beater. As the machine malfunctioned and could not be repaired, the remaining samples were lysed using corresponding settings on a TissueLyser II machine (30 Hz for 80 s). We confirmed no systematic differences in microbiota composition between differently lysed samples (data not shown). Lysed samples were incubated at 65°C for 30 min with inversion mixing every 10 min, followed by a 24:1 chloroform/isoamyl alcohol extraction (centrifugation at 13,000 rpm for 10 min). The supernatant was transferred to new tubes containing 0.6 vol isopropanol and incubated at room temperature for 15 min followed by centrifugation at 11,500 rpm at 4°C for 30 min. The isopropanol was subsequently removed and 1 mL of 70% ethanol was added to the tubes containing pelleted DNA, and left at room temperature for 1 h to absorb excess co-precipitated salts. Samples were then centrifuged for 10 min at 13,000 rpm, the supernatant discarded and the pellet washed again with 70% ethanol followed by another 10 min centrifugation at 13,000 rpm. Pellets were briefly air-dried to remove as much ethanol as possible before the addition of 20 μL of TE buffer, 30 mg of PVPP and 800 μL of high-salt TE buffer (1.5 M NaCl). Samples were briefly vortexed (1,400 rpm) before the addition of 200 μL of pre-warmed 0.7 M NaCl - 10% CTAB solution. Samples were again incubated at 65°C for 30 min, mixed by inversion every 10 min, followed by two rounds of chloroform/isoamyl alcohol extraction. The final supernatant was centrifuged for 2 min at 13,000 rpm to remove remaining PVPP particles and incubated overnight at −20°C with 0.1 vol 3 M sodium acetate (pH 5.2) and 2 vol of 70% ethanol. After two rounds of 70% ethanol wash, the final pellet was resuspended in 20 μL of 10 mM Tris-HCl (pH 8).

### 16S rRNA gene amplification and sequencing

DNA extracts were subjected to PCR amplification of the V3-V4 16S rRNA gene amplicon region using the 341F (<u>TCG TCG GCA GCG TCA GAT GTG TAT AAG AGA CAG</u> | CCT ACG GGN GGC WGC AG) and 785R primer pair (<u>GTC TCG TGG GCT CGG AGA TGT GTA TAA GAG ACA G</u> | GG ACT ACH VGG GTA TCT AAT CC) with Illumina-compatible Nextera adaptors (underlined sequence). A KAPA 3G Plant PCR kit was used for amplification with PCR conditions described by Perry et al. [2017]. Amplicon size and the absence of a band for negative controls (extraction and PCR controls) were verified on a 1% agarose gel with SYBR Safe DNA Gel Stain (Invitrogen). PCR products were purified using AMPure XP beads or the Zymo ZR-96 DNA Clean-Up Kit, and DNA concentration was quantified on an EnSpire Multimode Plate Reader using a Qubit High Sensitivity dsDNA kit (Invitrogen). Samples were normalised to 5 ng/μL for library preparation and sequencing by Auckland Genomics Ltd on an Illumina MiSeq with 2×300 bp chemistry (samples were demultiplexed by Auckland Genomics).

### Bacterial data analysis

The 16S rRNA gene amplicon sequences were processed in R [version 4.0.1; R Core Team 2019] using the DADA2 software package [version 1.16; Callahan et al. 2016] as per the authors’ recommendations. Forward and reverse reads were trimmed to 280 bp and 240 bp, respectively, following initial primer removal. Sequence reads shorter than the truncated value were discarded, as were reads where truncQ < 2 or the number of expected errors exceeded 3 for forward and reverse reads (--maxEE parameter). The DADA2 error learning algorithm was applied to the forward and reverse reads, which were subsequently dereplicated into unique sequences. The DADA2 core sample inference algorithm was then applied to dereplicated sequences, which were merged thereafter to obtain denoised, unique amplicon sequence variants (ASVs). Sequence chimeras were excluded and taxonomy assigned using the SILVA 138 ribosomal RNA database [Quast et al. 2013]. A phylogenetic tree was constructed using the DECIPHER [version 2.16.1; Wright 2016] and phangorn [version 2.5.5; Schliep 2011] packages.

The ASV table, taxonomic assignments and phylogenetic tree were merged with corresponding metadata to create a phyloseq object using the R [version 4.1.0, R Core Team 2021] package phyloseq [version 1.34.0; McMurdie and Holmes 2013]. Non-target sequences including chloroplasts and mitochondria were removed from the dataset using phyloseq’s prune_taxa function. We also removed low-abundance ASVs (total relative abundance < 0.0001%). The data were subsequently rarefied to the minimum number of reads across samples (n = 3,480; Supplementary Table 1) to accommodate for significant differences in sequencing output across samples [Weiss et al. 2017]; rarefaction was performed 5 times and all resulting Bray-Curtis distance matrices were highly correlated (Spearman coefficients = 0.999, p-values < 0.001) using Mantel tests with 9999 permutations [vegan package version 2.5-7; Oksanen et al. 2020].

Observed and Inverse Simpson alpha-diversity indices were calculated by applying phyloseq’s estimate_richness function to the rarefied ASV data. Observed richness is a direct measure of the number of microbiota species in a given sample, while the Inverse Simpson diversity index measures microbiota species richness as well as evenness to better represent overall community structure. We tested for associations between these alpha-diversity indices and sample covariates (sex, island location, antibiotic treatment, supplemental feeding, disease and hand-rearing) using Kruskal-Wallis tests (after confirming non-normality of alpha-diversity indices with Shapiro-Wilk tests) [vegan package] (Table 1). Post-hoc pairwise comparisons between covariate factors were performed using Dunn’s test [dunn.test package version 1.3.5; Dinno, 2017] with Benjamini-Hochberg p-value correction for significant Kruskal-Wallis models.

**Table 1:**
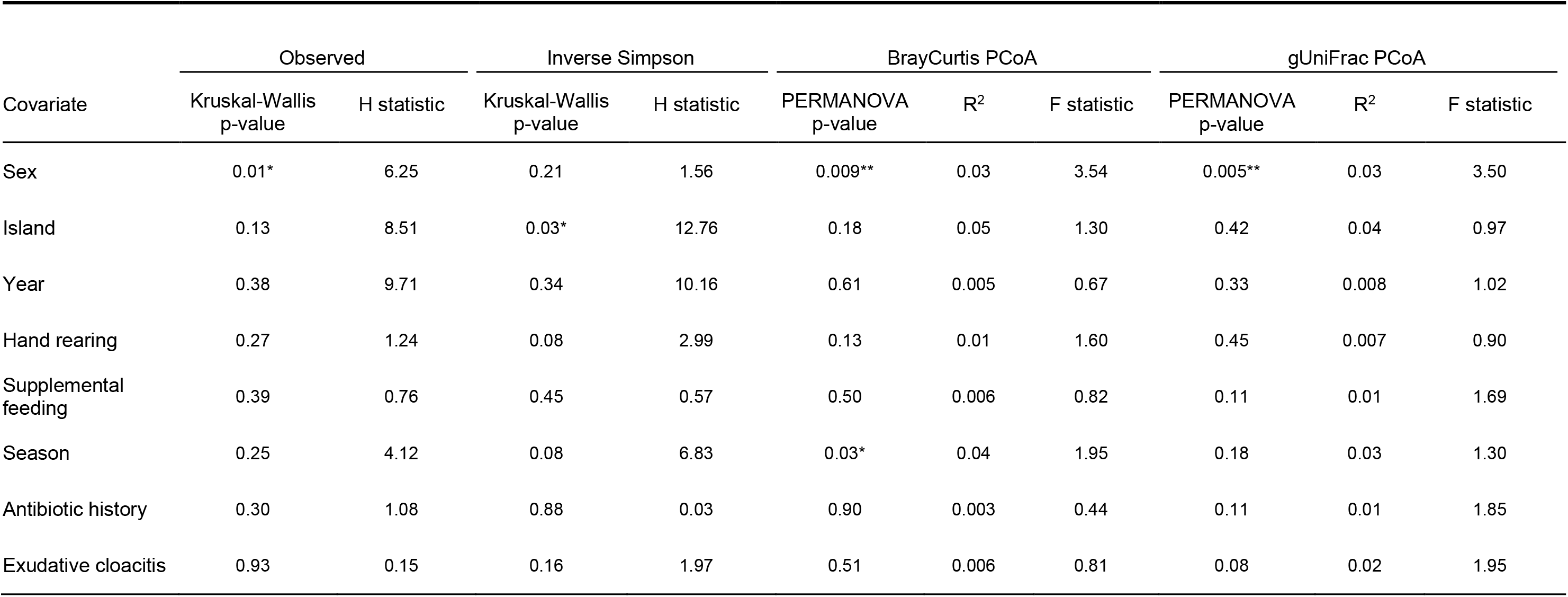
Kruskal-Wallis and PERMANOVA test results for associations between potential covariates and alpha-diversity and beta-diversity indices, respectively. * indicates marginal significance (p < 0.05), ** indicates significance (p < 0.01).

| Covariate | Observed |  | Inverse Simpson |  | BrayCurtis PCoA |  |  | gUniFrac PCoA |  |  |
| --- | --- | --- | --- | --- | --- | --- | --- | --- | --- | --- |
|  | Kruskal-Wallis p-value | H statistic | Kruskal-Wallis p-value | H statistic | PERMANOVA p-value | R<sup>2</sup> | F statistic | PERMANOVA p-value | R<sup>2</sup> | F statistic |
| Sex | 0.01* | 6.25 | 0.21 | 1.56 | 0.009** | 0.03 | 3.54 | 0.005** | 0.03 | 3.50 |
| Island | 0.13 | 8.51 | 0.03* | 12.76 | 0.18 | 0.05 | 1.30 | 0.42 | 0.04 | 0.97 |
| Year | 0.38 | 9.71 | 0.34 | 10.16 | 0.61 | 0.005 | 0.67 | 0.33 | 0.008 | 1.02 |
| Hand rearing | 0.27 | 1.24 | 0.08 | 2.99 | 0.13 | 0.01 | 1.60 | 0.45 | 0.007 | 0.90 |
| Supplemental feeding | 0.39 | 0.76 | 0.45 | 0.57 | 0.50 | 0.006 | 0.82 | 0.11 | 0.01 | 1.69 |
| Season | 0.25 | 4.12 | 0.08 | 6.83 | 0.03* | 0.04 | 1.95 | 0.18 | 0.03 | 1.30 |
| Antibiotic history | 0.30 | 1.08 | 0.88 | 0.03 | 0.90 | 0.003 | 0.44 | 0.11 | 0.01 | 1.85 |
| Exudative cloacitis | 0.93 | 0.15 | 0.16 | 1.97 | 0.51 | 0.006 | 0.81 | 0.08 | 0.02 | 1.95 |

To explore beta-diversity, we calculated Bray-Curtis and generalised UniFrac dissimilarity matrices for rarefied data using the phyloseq and GUniFrac [version 1.1; Chen 2018] packages, respectively, and subjected them to principal coordinate analysis (PCoA; vegan package). We then tested the association between beta-diversity indices and all covariates by subjecting the distance matrices to PERMANOVA (vegan) with 9999 permutations (Table 1). We further tested for homogenous group dispersion using the vegan functions betadisper and permutest. The DESeq2 package [version 1.30.0; Love et al. 2014] was used to test for differential abundance of ASVs (test = ‘Wald’. fitType = ‘local’) for significant covariates using non-rarefied data (p-values Benjamini-Hochberg adjusted).

We then agglomerated the phyloseq object at the taxonomic species level and used the plyr package [version 1.3.5; Wickham 2011] to group less abundant ASVs into the category ‘Others’ based on a per-species mean relative abundance of < 0.3%. We defined the core species as those species with a minimum prevalence of 85% and a minimum average relative abundance of 1% across all samples. The relative abundance of these core taxa were then used as taxon-specific phenotypes (see Materials and Methods: Genotype-phenotype analyses).

All data were visualised using the R packages ggplot2 [version 3.3.3; Wickham 2016], cowplot [version 1.1.1; Wilke 2020], and Manu (‘kākāpō’ colour palette) [version 0.0.1; Ram et al. 2018; Thomson 2020].

### Genotype-phenotype analyses

The genomic data of 169 kākāpō individuals were generated within the Kākāpō125+ project to represent the entire extant kākāpō population as well as previously deceased individuals. To identify a high-confidence single-nucleotide polymorphism (SNP) call set, the high-quality kākāpō reference genome (NCBI taxonomy ID: 2489341) was combined with short-read whole-genome sequencing data for the 169 individuals (2×125 bp or 2×150 bp Illumina paired-end sequencing; mean genome-wide coverage of 24.4, ranging from 13.1 to 53.4 across individuals) using DeepVariant (see Guhlin et al. 2022 for details). We used PLINK [version 1.9; Purcell et al. 2007] to filter this SNP call set for genotype missingness (> 20%), minor allele frequency (< 5%), extreme deviation from Hardy-Weinberg equilibrium (P < 10^−7^) and bi-allelicity across the 133 individuals for which we had also obtained microbiota data, resulting in 1,202,717 SNPs to be used in downstream analyses. We then pruned the SNPs for strong linkage disequilibrium (using PLINK’s indep-pairwise function with a window size of 50, a step size of 10 and a pairwise r^2^ threshold of 0.8) and used this set of SNPs to calculate the variance-standardised relationship matrix and the first 10 principal components (PCs) of the principal component analysis (PCA) of this relationship matrix.

Microbiota-related phenotypes included observed and Inverse Simpson alpha-diversity indices, dominance of *Escherichia-Shigella coli* (hereafter referred to as *ES. coli* dominance: binary variable describing if *Escherichia-Shigella coli* comprised at least 50% of rarefied sequence reads in a given sample), the first PCoA axes of the Bray-Curtis and gUniFrac ordinations, and relative sequence abundances of the three core species *Escherichia-Shigella coli*, *Escherichia-Shigella fergusonii*, and *Tyzzerella* sp. (see Results: Microbiota diversity). All quantitative phenotypes were rank-standardised in R (version 4.1.0) using the package xavamess [version 0.6; Robin 2018]. Covariates that were significantly associated with phenotypes under previous Kruskal-Wallis and PERMANOVA testing (see Materials and Methods: Bacterial data analyses) were identified as confounding factors and thus controlled for during subsequent phenotypic analyses.

We assessed the relationship between microbiota and genomic variation using the following pipeline. BayesR [version 2; Moser et al. 2015] was used to estimate heritability and partition heritability by chromosome for all phenotypes through Bayesian mixture modelling (based on MCMC permutation) with default total iteration (n = 50,000) and burn-in (n = 20,000) settings. We increased the default settings for the observed richness phenotype as we initially found a posterior mode estimate of heritability of 0%, but even after increasing the total number of iterations to 200,000 (burn-in of n = 50,000) we found no evidence for non-zero heritability. We therefore excluded the observed richness phenotype from all downstream analyses. Chromosome partitioning and MCMC chain plots (Supplementary Figure 2) were plotted using the R packages ggplot2 and cowplot.

To then investigate whether any heritable phenotypes were directly related to genome-wide inbreeding we correlated inbreeding coefficients with rank-standardised phenotypes using Pearson and Spearman correlations with the R package stats [version 4.1.0, R Core Team 2021]. Inbreeding coefficients were calculated based on the SNP callset after removing sex chromosomes using vcftools --het using default settings [version 0.1.15; Danecek et al. 2011].

The heritable traits were then subjected to genome-wide association studies (GWAS). We used RepeatABEL [version 1.1; Ronnegard et al. 2016] to perform association tests between SNPs and microbiota-related phenotypes while controlling for kākāpō sex and the first three PCs of the relationship matrix PCA (see Supplementary Figure 3 for PCA screeplot). RepeatABEL output files were then processed in R (version 4.1.0) to remove sex chromosomes. Manhattan and associated QQ plots (Supplementary Figure 4) were created using the qqman package [version 0.1.8; Turner 2018] and coloured based on the Manu package ‘kākāpō’ palette. We also confirmed that outlier individuals (n = 3) with substantially different bacterial compositions did not have any significant effect on the associations detected by GWAS by removing these from the dataset and re-running the analyses. We further calculated genomic inflation of the p-values using the GenABEL R package [version 1.8-0, Aulchenko et al. 2007]. We identified genomic regions of interest by isolating the regions spanned by peaks of nominally significant SNPs at a stringent significance threshold ***α*** of 0.001. We then identified genes underlying these nominally significant regions using a sliding-window approach (window size of 10 kbp, sliding step of 1 kbp; as optimised in Urban et al. [in prep.]) by averaging the SNP-wise to window-wise p-values using bedtools [version 2.29.2; Quinlin and Hall 2010].

We performed gene-set analyses on our SNP-wise GWAS results using the Magma software, with default settings [version 1.08; de Leeuw et al. 2015], to identify gene pathways significantly associated with microbiota-related phenotypes. We used a chicken-specific Gene Ontology (GO) pathway set, downloaded from ge-lab.org (Gallus_gallus_5.0) which, to our knowledge, was the most complete curated avian gene pathway database available. Magma output files were exported to R where the p-values were adjusted for multiple testing using Bonferroni correction [stats version 4.0.3; R Core Team 2021; Hochberg 1998]. Only the pathways that were significant at threshold ***α*** = 0.05 following Bonferroni correction are reported; gUniFrac was the only phenotype that had no significant pathways following multiple-testing correction.

## Results

### Bacterial diversity

In total, 26,649,116 raw 16S rRNA gene amplicon sequence reads (forward and reverse) were combined from available and new sequencing data, with 8,648,731 merged reads remaining following quality and chimera filtering (Materials and Methods). After removing non-target and low-abundance ASVs, 4,138 unique ASVs were identified across the dataset. Following rarefaction to 3,480 reads per sample, 3,478 unique ASVs remained. The average number of ASVs per sample was 73, ranging from 9 to 602 ASVs per sample. Overall, 98.99% of ASVs could be taxonomically assigned to a known phylum, while 77.9% were assigned to species level.

*Proteobacteria* was by far the most abundant phylum (80% of sequence reads), followed by *Firmicutes* (18.5% of reads). As previously reported for kākāpō, *Escherichia-Shigella coli* was the most abundant species, dominating the microbiota of most kākāpō faecal samples and representing 70% of all sequence reads (Figure 2A). *Tyzzerella* sp. was the second most abundant species at 14.5% of reads, followed by *Escherichia-Shigella fergusonii* at 2.3%. We defined these three taxa as the core bacterial microbiota of the kākāpō gut since they were the only species present at > 1% average relative sequence abundance in > 85% of kākāpō faecal samples (Figure 2B). Samples not dominated by *Escherichia-Shigella* exhibited greater relative abundance of *Tyzzerella* species and a generally more diverse gut microbiota (Figure 2A). The faecal sample of the kākāpō Richard Henry was anomalous, being dominated by *Pantoea* species which were not represented in appreciable numbers (> 2 reads per sample) in any other faecal sample.

**Figure 2:**
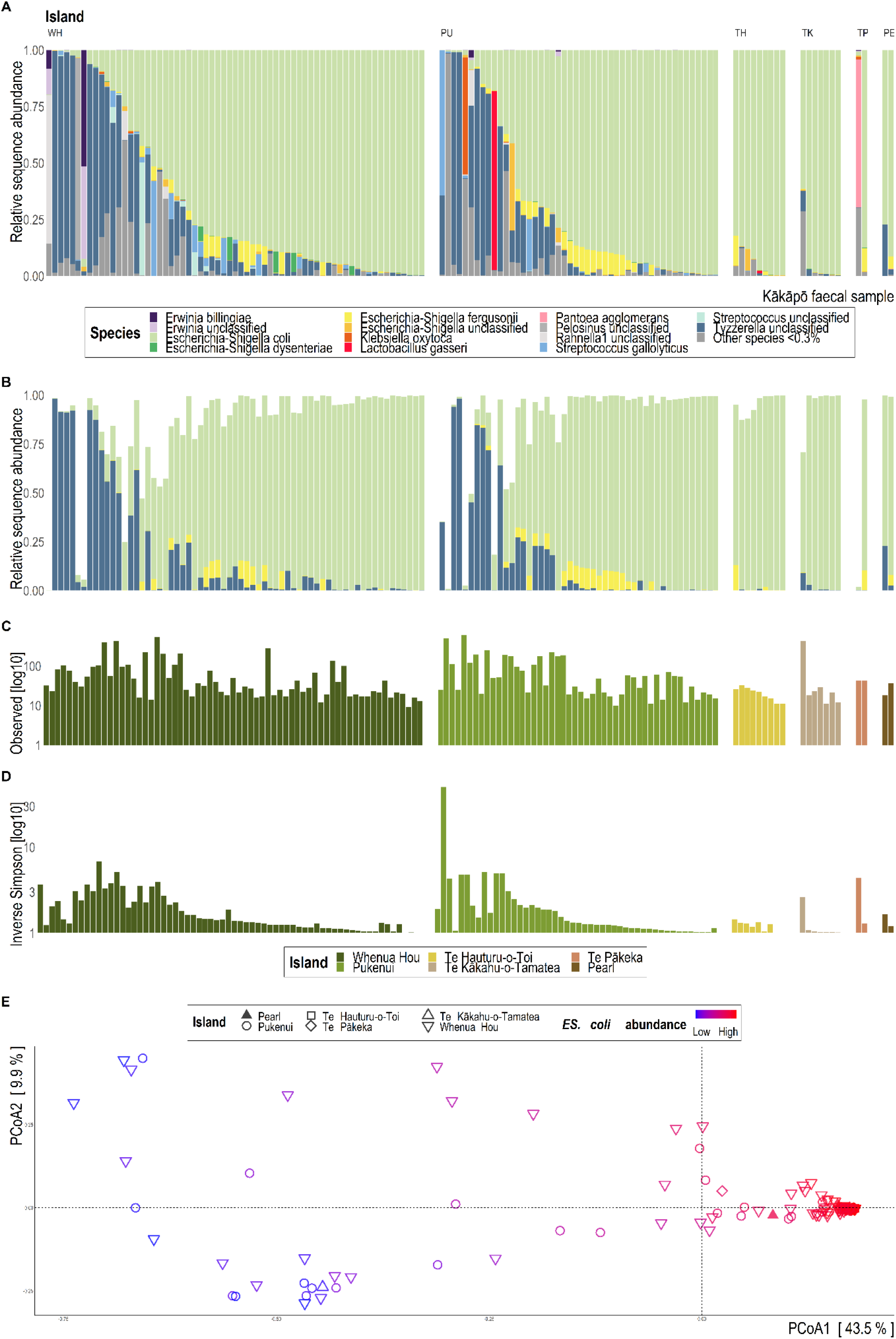
*[A] 16S rRNA gene sequence-based taxonomic distribution of bacteria within kākāpō faecal samples at species level. Taxa with mean relative 16S rRNA gene sequence abundance < 0.3% are grouped together as ‘Other species’. Each bar represents a single kākāpō faecal sample, which are grouped by island location: WH = Whenua Hou, PU = Pukenui, TH = Te Hauturu-o-Toi, TK = Te Kākāhu-o-Tamatea, TP = Te Pākeka, PE = Pearl Island. [B] Relative sequence abundance of the core bacterial species in the kākāpō gut microbiota defined as > 1% relative abundance in > 85% of individuals. [C] Log10-scaled alpha-diversity observed richness indices for each kākāpō faecal sample. [D] Log10-scaled alpha-diversity Inverse Simpson indices for each kākāpō faecal sample. [E] 16S rRNA gene sequence-based Bray-Curtis dissimilarity distances visualised via principal coordinate analysis ordination. Each dot of the PCoA represents the microbiota of a single kākāpō faecal sample. Samples are coloured by relative abundance of* Escherichia-Shigella coli *and shaped by island location*.

We obtained a mean observed richness diversity estimate of 73.1 (SD ± 107) and a mean Inverse Simpson diversity estimate of 2.2 (SD ± 4.3) across all samples. Observed bacterial richness was significantly associated with kākāpō sex (p = 0.01; Table 1), where male kākāpō exhibited lower bacterial richness than females, while the Inverse Simpson diversity index was marginally significantly associated with island location (p = 0.03; Figure 2D; Table 1). Post-hoc Dunn’s tests for pairwise comparisons showed that the significant differences in Inverse Simpson diversity index across islands were due to differences between Te Kākahu-o-Tamatea (TK) and Pukenui (PU) samples (p-value = 0.03) and TK and Whenua Hou (WH) samples (p-value = 0.02).

Bacterial beta-diversity, as measured by Bray-Curtis and gUniFrac dissimilarity matrices, was significantly associated with kākāpō sex (p < 0.01; Table 1; Materials and Methods). Season had a marginally significant effect on variation among bacterial communities (p = 0.03; Table 1), yet showed heterogeneous group dispersions (p = 0.01; Materials and Methods) which may account for much of the variation observed by the PERMANOVA. Bacterial community separation along PCoA ordination axes for both Bray-Curtis and gUniFrac indices was largely due to *Escherichia-Shigella coli* dominance (Figure 2E; Supplementary Figure 5). We confirmed that no other variable had an impact on bacterial beta-diversity by testing and visualising the potential influence of year of sample collection, supplemental feeding, season, antibiotic administration and exudative cloacitis on beta-diversity indices (Table 1; Supplementary Figure 6).

Sex was the only variable that was significantly associated with multiple indices of bacterial diversity (Table 1). We subsequently found five ASVs exhibited nominally significant differential abundance between male and female kākāpō (p < 0.05). Female kākāpō hosted 2-fold and 1.1-fold greater relative sequence abundance of a *Tyzzerella* ASV and an *ES. coli* ASV, respectively, than male kākāpō. Males exhibited 2.6-fold, 2.8-fold and 3-fold greater relative sequence abundance of two unclassified *Escherichia-Shigella* ASVs and one *Streptococcus sp*. ASV, respectively. While island location and season showed marginally significant associations (Table 1), these associations were driven by, in the case of island location, differences between the small TK group (n = 7) and our two largest sampling groups (WH = 65, PU = 48) respectively, and, in the case of season, heterogeneous group dispersions.

### Associations between bacterial and genomic diversity

Using a Bayesian hierarchical model, as implemented in BayesR, we found that most microbiota-related phenotypes had a posterior mode heritability estimate of approximately 30%, albeit with very wide 95% credible intervals (CI) (Table 2; Materials and Methods). Only observed richness showed no evidence of heritability (point estimate of 0%), even after increasing the number of MCMC chains and burn-in steps (Materials and Methods). We therefore excluded the observed richness phenotype from all downstream analyses. Furthermore, none of the phenotypes included in this study were significantly associated with genome-wide inbreeding (Table 3).

**Table 2:**
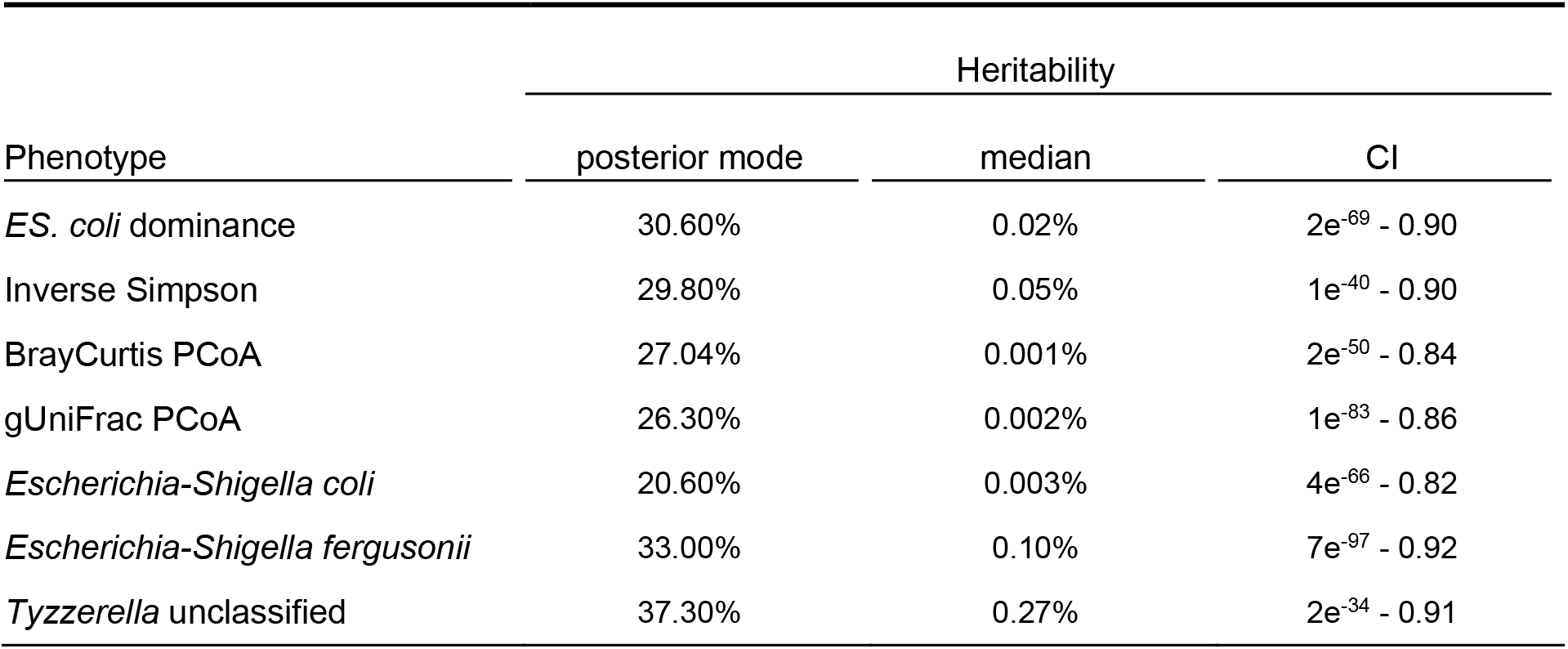
Posterior mode and median heritability for each phenotype derived from a Bayesian hierarchical model implemented in BayesR, with respective 95% credible intervals (CI). Posterior density plots are provided by Supplementary Figure 7.

| Phenotype | Heritability |  |  |
| --- | --- | --- | --- |
|  | posterior mode | median | CI |
| *ES. coli* dominance | 30.60% | 0.02% | 2e<sup>-69</sup> - 0.90 |
| Inverse Simpson | 29.80% | 0.05% | 1e<sup>-40</sup> - 0.90 |
| BrayCurtis PCoA | 27.04% | 0.001% | 2e<sup>-50</sup> - 0.84 |
| gUniFrac PCoA | 26.30% | 0.002% | 1e<sup>-83</sup> - 0.86 |
| *Escherichia-Shigella coli* | 20.60% | 0.003% | 4e<sup>-66</sup> - 0.82 |
| *Escherichia-Shigella fergusonii* | 33.00% | 0.10% | 7e<sup>-97</sup> - 0.92 |
| *Tyzzarella* unclassified | 37.30% | 0.27% | 2e<sup>-34</sup> - 0.91 |

**Table 3:**
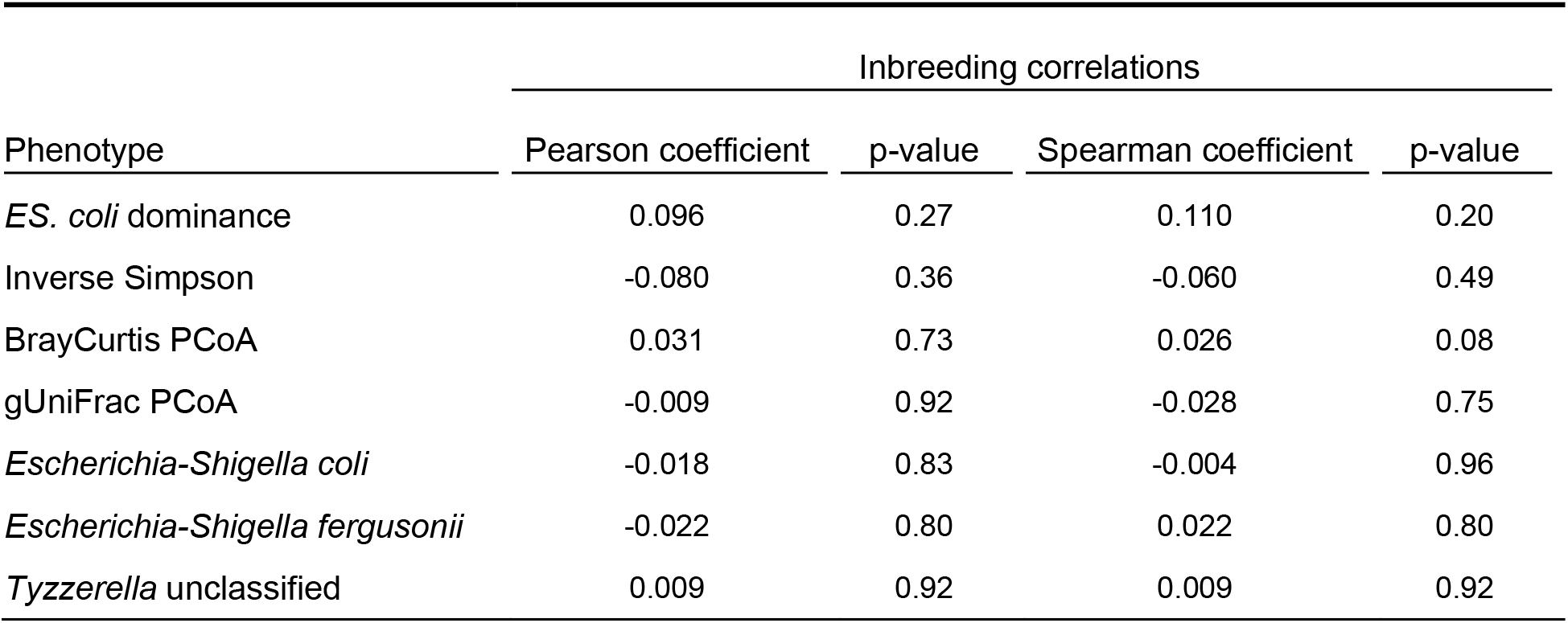
Pearson and Spearman correlations and p-values to test associations between kākāpō genome-wide inbreeding coefficients and microbiota phenotypes.

Sex was the only factor significantly associated with gut microbiota diversity, and was therefore included as a covariate of our genotype-phenotype analyses together with the first three genomic-based PCs. All heritable microbiota-related phenotypes were identified as polygenic traits, with longer chromosomes explaining larger amounts of heritability (Figures 3 & 4, top panel). According to our GWAS, no SNPs were significantly associated with microbiota-related phenotypes at a stringent genome-wide significance threshold ***α*** of 4×10^−8^ (assuming Bonferroni multiple testing correction across all SNPs). However, we found many significant associations at a stringent nominal significance threshold ***α*** of 0.001 for all phenotypes (Figures 3 and 4, bottom panel; nominally significant SNPs are coloured in red). We then aggregated the p-values in a sliding-window approach (Supplementary Figure 8; Materials and Methods) to leverage the spatially aggregated signals in order to locate candidate genes potentially underlying our nominally significant genomic regions (Table 4).

**Figure 3:**
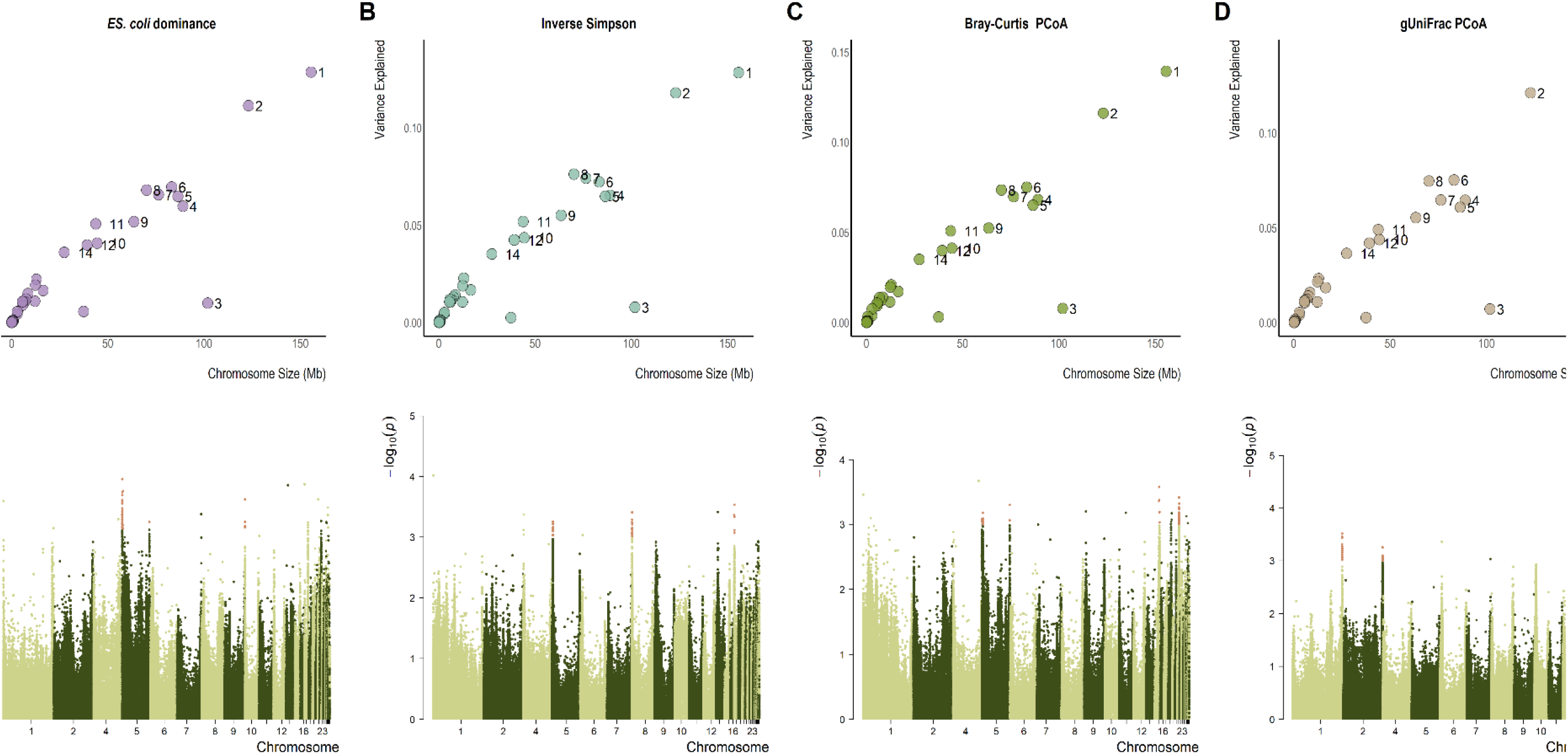
*[Top] Chromosome partitioning plots for alpha- and beta-diversity related phenotypes: [A]* ES. coli *dominance [B] Inverse Simpson diversity [C] Bray-Curtis PCoA diversity and [D] gUniFrac PCoA diversity. Each dot represents a kākāpō chromosome, where chromosome size is plotted against the amount of variance explained by that chromosome regarding the microbiota-related phenotypes. Chromosomes 3 and 13 are the kākāpō sex chromosomes. [Bottom] Manhattan plots below their corresponding chromosome partitioning plots for each alpha- and beta-diversity related phenotype. Each dot represents a SNP plotted against its log10-transformed significance value; those coloured red are nominally significant SNPs where p < 0.001*.

**Figure 4:**
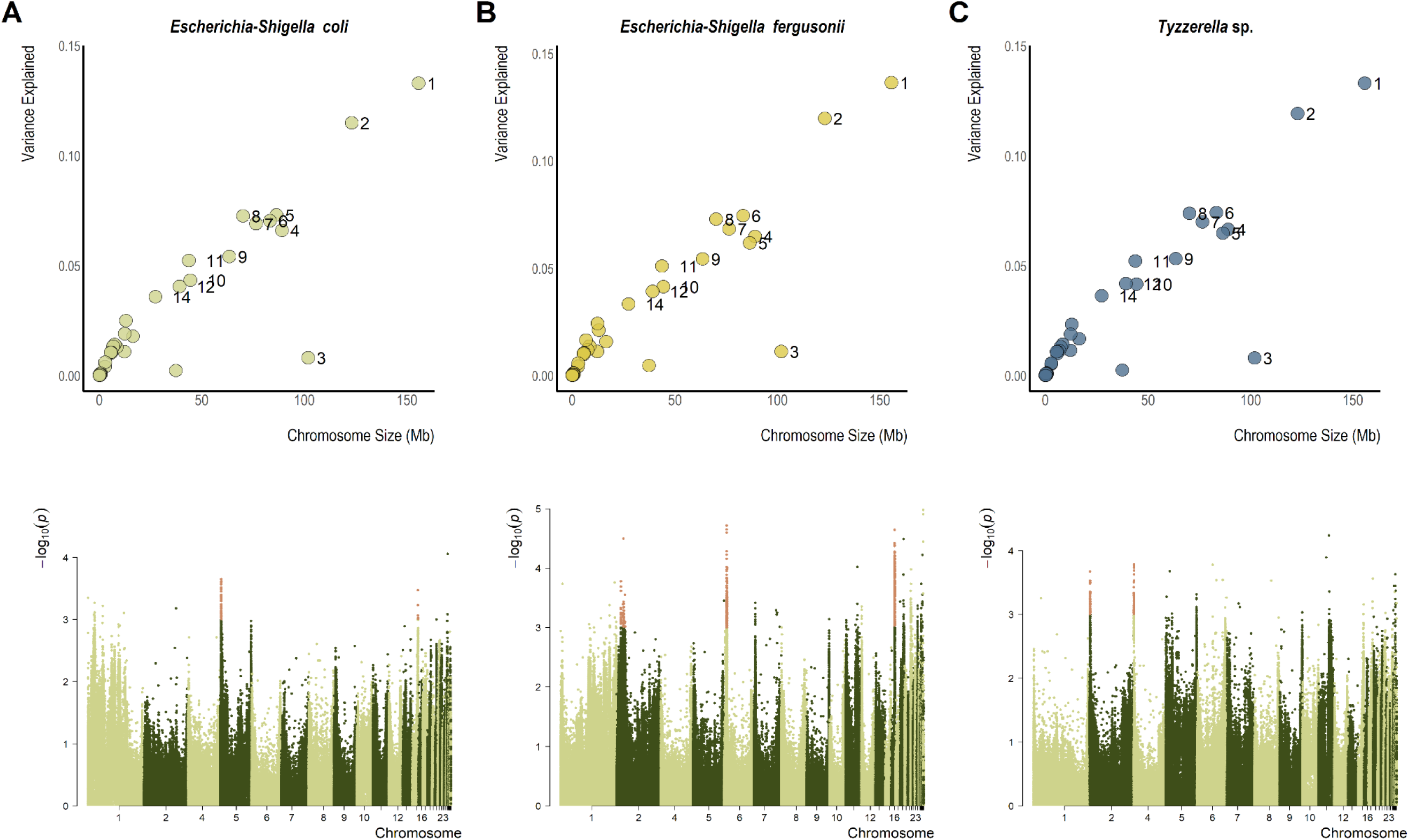
*[Top] Chromosome partitioning plots for core taxon relative abundance phenotypes: [A]* Escherichia-Shigella coli *[B]* Escherichia-Shigella fergusonii *and [C]* Tyzzerella *sp. Each dot represents a kākāpō chromosome, where chromosome size is plotted against the amount of variance explained by that chromosome regarding the microbiota-related phenotypes. [Bottom] Manhattan plots below their corresponding chromosome partitioning plots for each core taxon relative abundance phenotype. Each dot represents a SNP plotted against its log10-transformed significance value; those coloured red are significant SNPs where p < 0.001*.

**Table 4:**
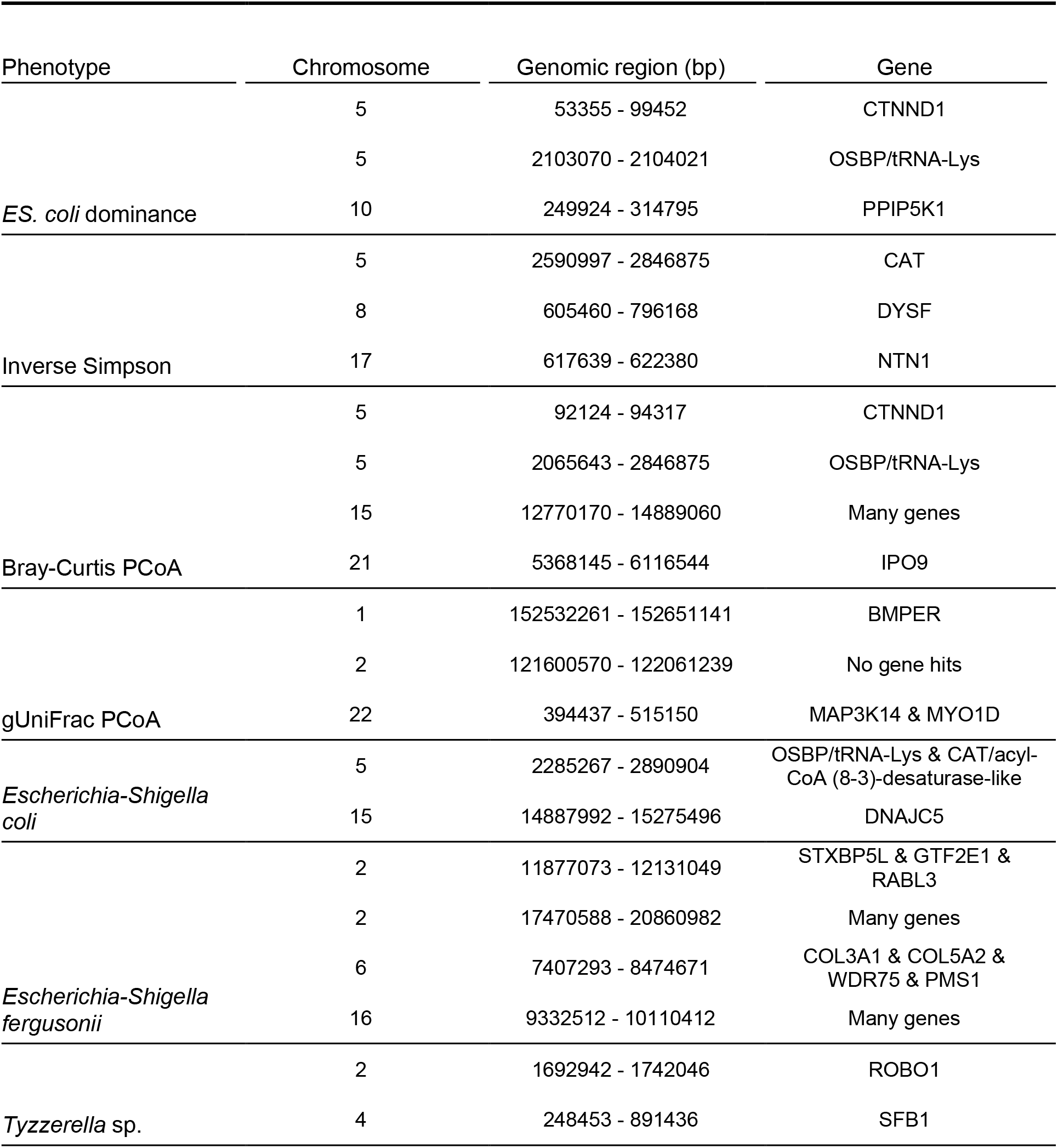
Candidate genes located in the genomic regions that are nominally significantly associated with microbiota-related phenotypes (Materials and Methods; refer to Supplementary Figure 8 for regions spanned by nominally significant SNPs and overlapping gene regions).

Our gene-set analyses identified several GO pathways that were significantly associated with gut microbiota-related phenotypes after Bonferroni multiple-testing correction across pathways. These pathways described several metabolic and cellular functions as well as processes associated with oxygen detection, vitamin B_12_ and growth factor activity (Table 5).

**Table 5:**
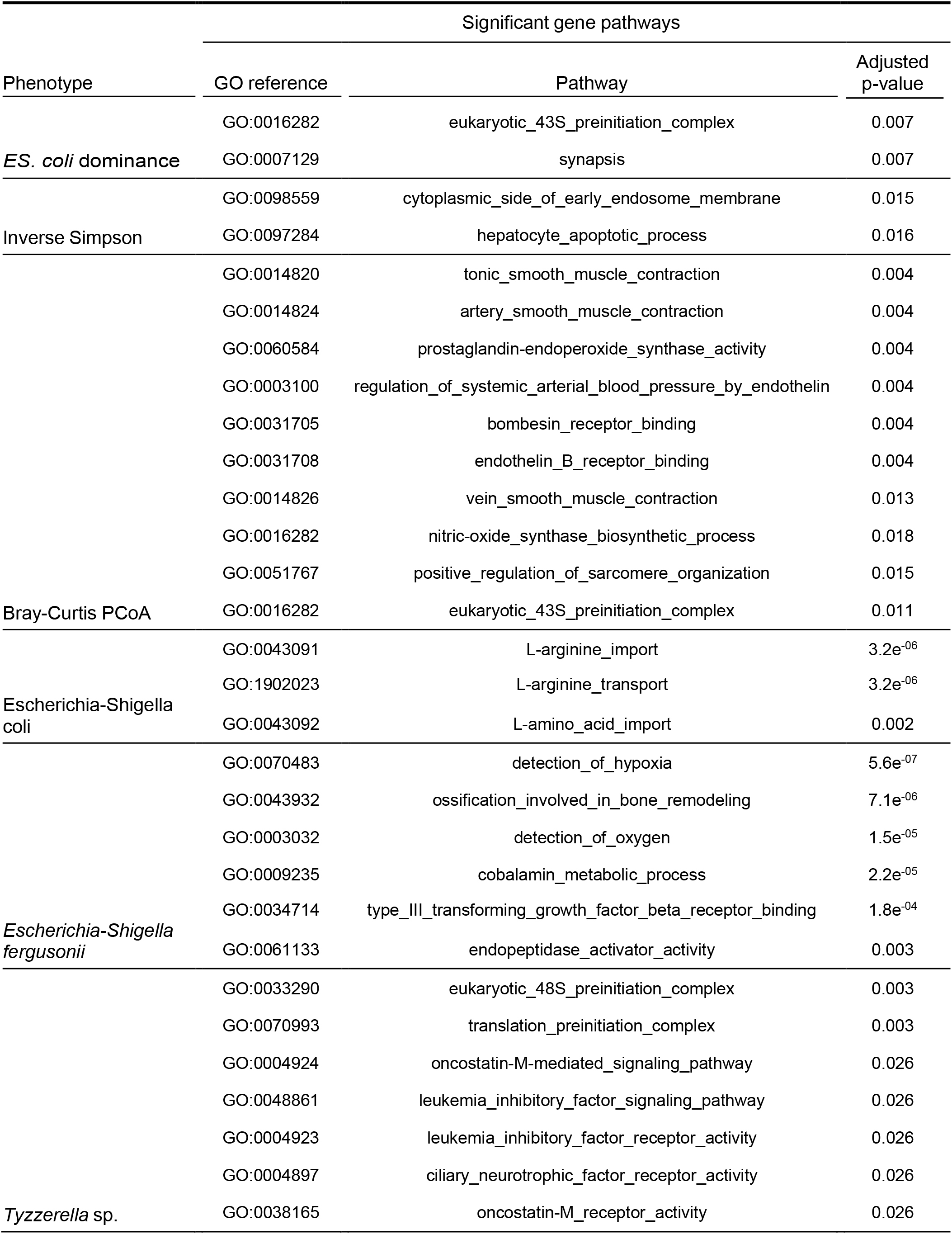
Gene Ontology (GO) pathways significantly associated with microbiota phenotypes. P-values are adjusted using Bonferroni multiple-testing correction across test pathways.

## Discussion

Our study of the kākāpō gut microbiota represents, to our knowledge, the first attempt to describe the bacterial diversity for virtually an entire species and subsequently assess the relationship between microbiota and host genomic diversity in a critically endangered population. We created a microbiota catalogue for as many extant kākāpō as possible to understand the true extent of species-wide bacterial diversity and its consistency across individuals. Indeed, in our effort to catalogue the entire population we revealed unique, significantly dissimilar microbiotas in some individuals. We used these data to study the species-wide relationship between microbiota and genome-wide diversity, taking into account extensive environmental data and individual-specific characteristics. While our genetic architecture and association analyses confirm that microbiota diversity is a highly polygenic trait in kākāpō, as expected from insights into related human studies [e.g., Ishida et al., 2020], our sampling approach allows us to be certain that we have captured the entirety of this complex relationship in a single species.

By sampling the microbiota of 84% of all surviving kākāpō and including data from additional deceased birds, we were able to draw a representative picture of overall gut bacterial diversity. Despite our best efforts to obtain faecal samples for the entire extant kākāpō species over a two year sampling period, we could not obtain faecal samples for 22 kākāpō. All kākāpō live in the wild with minimised human impact. Faecal sampling was also performed as part of routine monitoring and we were thus unable to increase our sampling efforts further. However, as a critically endangered species with a world-wide census population size of 201 (as of 15/11/2021), the sample size of our study remains small in comparison with other quantitative genomic analyses [e.g., Hong & Park, 2012]. The lack of genome-wide significant genomic regions may therefore be due to reduced power in this study. We were, however, able to identify genomic regions which, at a stringent nominal significance threshold, set themselves apart from the genomic background. Moreover, these regions contained genes that have previously been implicated in gut homeostasis and related pathways in other species. As large-scale host genomic studies of the microbiota in humans have shown similar results with respect to the polygenicity of bacterial diversity [Ishida et al., 2020], we are confident that this study of species-wide bacterial diversity and its relationship with the host genome delivers new insights into the role of microbiota in endangered species.

### The kākāpō gut microbiota

We corroborated previous findings regarding the dominance of the *Escherichia-Shigella* genus in the kākāpō gut microbiota [Waite et al., 2018], with *Escherichia-Shigella coli* identified as the most dominant species. Additional species of the *Escherichia-Shigella* genus, including *Escherichia-Shigella fergusonii*, were also prevalent and abundant throughout the dataset. We therefore confirmed this unusual dominance of one bacterial genus in the gut of a herbivorous species as a true biological peculiarity. We further identified unclassified *Tyzzerella* species and *Streptococcus gallolyticus* as prominent members of the kākāpō gut microbiota, both of which are frequent inhabitants of the avian digestive tract and of the rumen of other herbivorous species [Curtis et al. 2018; Pasquereau-Kotula et al. 2018; Yan et al. 2019; Husso et al. 2020; Liu et al. 2020a]. Generally, we observed the relative abundance of *Tyzzerella* sp. to increase as *Escherichia-Shigella coli* dominance decreased (Figure 2), though this could in part reflect the compositional nature of our rarefied data [McMurdie and Holmes 2014; Gloor et al. 2017; Weiss et al. 2017]. Some species of *Clostridia* (the class to which *Tyzzerella* bacteria belong) are considered positive commensal bacteria in the gut of humans and other mammals [Guo et al. 2020], though whether this holds true for avian species remains to be determined. We further identified Richard Henry to be the only kākāpō individual with a gut microbiota dominated by several *Pantoea* species. As Richard Henry was the last surviving kākāpō from the New Zealand mainland, with all other kākāpō originating solely from one offshore island (Rakiura / Stewart Island), this pattern may reflect lost historical geographic variation in gut microbiota composition with potentially important implications for future reintroductions. As we also assessed the microbiota of several of his offspring, we can rule out that the distinct genomic background of Richard Henry is responsible for his unique microbiota.

We found that kākāpō sex was significantly associated with bacterial alpha- and beta-diversity, an association which has been previously reported in other avian species [Borda-Molina et al. 2020; Liu et al. 2020; Zhu et al. 2020]. However, relationships between sex and the gut microbiota are often difficult to interpret in the presence of other influential factors such as diet and habitat [Elderman et al. 2018; Kim et al. 2020]. All other environmental factors and characteristics of individual kākāpō showed no significant association with the gut microbiota, including variables such as supplemental feeding and antibiotic treatment. Island residence and season of sample collection showed marginally significant associations with bacterial diversity, but we attributed these signals to substantial discrepancies in sample size and heterogeneous group dispersions, respectively. We also found no evidence that exudative cloacitis, the most prevalent disease amongst adult kākāpō which causes an inflammation of the cloaca and lower reproductive tract, was associated with gut microbiota diversity. Since the causative agent of this disease is still unknown this result might point to a viral or fungal entity as opposed to a bacterial pathogen, requiring the application of metagenomic or metatranscriptomic approaches. However, as the kākāpō individuals did not necessarily suffer from cloacitis inflammation at the time of sampling, another explanation for this null result could conceivably reflect the temporal variability of the gut microbiota with respect to disease outbreaks.

### Associations between bacterial and genomic diversity

We report evidence of a relationship between kākāpō genomic variation and their gut microbiota. Using Bayesian mixture modelling, we found some evidence of non-zero heritability for nearly all traits describing gut bacterial diversity, including alpha-diversity, beta-diversity and taxon-specific abundances, and described the complex polygenic architecture underlying these traits (Figures 3 and 4). Given the previously-described complex relationship between host genomic diversity and their microbiota, the polygenicity of kākāpō gut microbiota-associated phenotypes was an expected outcome [Goodrich et al. 2017; Awany et al. 2019]. While this polygenic architecture does not allow us to fully understand the genetic underpinnings of the kākāpō gut microbiota, we used GWAS and gene-set enrichment analyses based on chicken-specific gene pathways to identify associated genes and pathways and assess their relationship with bacterial diversity.

We found suggestive evidence of a relationship between the kākāpō gut microbiota and homeostasis of the host’s intestinal mucosal epithelium. GO pathways such as nitric oxide synthesis, L-arginine transport, transforming growth factor beta (TGF-***β***) receptor binding and prostaglandin-endoperoxide synthase were significantly associated with various traits describing gut bacterial diversity (Table 5). These gene pathways, while acting in many physiological systems, have important roles in the gastrointestinal tract. L-arginine is required to synthesise nitric oxide [Moncada and Higgs 1995; Coleman 2001], an essential component in mucosal epithelial defense [Kubes and Wallace 1995; Rosselli et al. 1998; Lundberg et al. 2008, Cinelli 2020] and commensal gut microbiota homeostasis [Inserra et al. 2019]. As a powerful vasodilator and signalling molecule, nitric oxide also protects the mucosal lining and stimulates wound healing by increasing blood flow to sites of injury or irritation, increasing mucus production and modulating immune response against irritants and microorganisms in the mucosal lining [Kubes and Wallace 1995; Bjornë et al. 2004; Wink et al. 2011; Jӓdert et al. 2012]. Correspondingly, we identified gene pathways significantly associated with microbiota-related phenotypes that were related to smooth muscle contraction, blood pressure regulation and pulmonary vasoconstriction via endothelin B receptor activity [Ladenheim 2013; Kowalczyk et al. 2015; Nguyen and Gerstein 2019] (Table 5). TGF-***β*** is a cytokine/growth factor that plays a vital role in intestinal epithelial homeostasis [Lichtman et al. 2016], while the prostaglandin-endoperoxide synthase pathway maintains the integrity of the gastric and intestinal mucosal lining [Peskar 2001]. This relationship between the gut microbiota and intestinal mucosal epithelium homeostasis can further be corroborated by individual genes that are amongst those most significantly associated with bacterial diversity, including *CTNND1*, *COL3A1*, *COL5A2*, and *PPIP5K1*, which are known to, amongst other functions, play important roles in gut homeostasis (Table 4; Supplementary Figure 8). *CTNND1* contributes to epithelial homeostasis as a core component of adherens junctions [Smalley-Freed et al. 2010; Daulagala et al. 2019], while *COL3A1* and *COL5A2* have functions in wound healing of the intestinal epithelium [Kuivaniemi and Tromp 2019]. The *PPIP5K1* kinase has been implicated in the formation of gastric leiomyoma submucosal tumors [Gu et al. 2017]. Chronic inflammation of the mucosal epithelium has also previously been associated with an over-representation of *Enterobacteriaceae* and *Verrucomicrobiaceae* bacteria in humans [Ahmed et al. 2018], both of which are observed in the kākāpō gut microbiota.

We further found suggestive evidence of an interaction between gut bacterial diversity and host inflammation and immune response. TGF-***β***, Oncostatin M, and leukemia inhibitory factor (Table 5) are cytokines that play important roles in the inflammatory response of multiple host organs, including the gut. TGF-***β*** modulates intestinal inflammation and is an important regulator of T-cell activity and differentiation [Wahl et al. 2004; Li and Flavell 2008; Hou et al. 2018]. Oncostatin M is upregulated in the inflamed epithelium of humans with chronic ulcerative colitis and inflammatory bowel disease [Li et al. 2020], while leukemia inhibitory factor promotes intestinal epithelial cell proliferation and has elevated expression in inflamed colon tissues [Guimbaud et al. 1998; Nicola and Babon 2015]. Moreover, nitric oxide (Table 5) modulates the immune response against microorganisms in the mucosal lining [Kubes and Wallace 1995; Bjorn et al. 2004; Wink et al. 2011; Jadert et al. 2012] and constitutes a vital part of the immune response through, e.g., mediation of T-cell activity [García-Ortiz and Serrador 2018] and inhibition of reactive oxygen species [Coleman 2002]. Reactive oxygen species are produced in excess upon inflammation of the intestinal lining [Kowalczyk et al. 2015] and are also known to be regulated by essential antioxidant enzymes directly encoded by the *CAT* gene, one of our top candidate genes (Table 4; Supplementary Figure 8).

Finally, we found some evidence of an association between the kākāpō gut microbiota and host metabolism. For example, the *OSBP* gene (Table 4, Supplementary Figure 8) is known to be responsible for cholesterol homeostasis [Mesmin et al. 2013; Kentala et al. 2016], with mounting evidence for an indirect regulation of cholesterol homeostasis by gut microbiota [Le Roy et al., 2019]. The cobalamin (vitamin B_12_) metabolic pathway was further significantly associated with *Escherichia-Shigella fergusonii* abundance (Table 5); Vitamin B_12_ deficiency can significantly alter the composition of intestinal microbiota, as shown in mice where deficiency enabled the proliferation of taxa such as *Shigella* [Lurz et al. 2020].

## Conclusions

This study assesses the bacterial diversity of virtually an entire extant species and presents first evidence of associations between the genome of a critically endangered species and its gut microbiota. An improved understanding of the kākāpō gut microbiota – and its relationship with host genomics – can directly benefit kākāpō management and conservation by providing new insights into the role of the gut microbiome in kākāpō health and disease mitigation. We were able to identify putative associations between the gut microbiota and functional pathways related to intestinal homeostasis, inflammation, immune response and metabolism. We propose future studies to elucidate the specific effects of factors such as disease and supplemental feeding on our observed relationships between microbial and genomic diversity. To achieve a complete understanding of the complex link between the gut microbiota and host functions related to immunity, inflammation or disease susceptibility, we aim to apply in-depth functional meta-omic approaches covering viral, archaeal and eukaryotic diversity and compare them with our current analyses which are solely focused on bacteria. Overall, we suggest an integration of microbiome studies in conservation research and management to obtain a better understanding of how the concept of One Health with its implications for human, animal and environmental welfare can be achieved.

## List of abbreviations

ASV: amplicon sequence variant
SNP: single nucleotide polymorphism
GWAS: genome-wide association study
PCA: principal components analysis
PCoA: principal coordinate analysis
TGF-*β*: transforming growth factor beta

## Contributions

AW, AD, MT and LU conceived the study. AW and AD organised the samples. The entire Kākāpō Recovery Programme, most importantly Daryl Eason, Lydia Uddstrom, Deidre Vercoe and Jodie Cran, helped with the collection of the samples. AW performed all laboratory work. AW and LU performed all computational analyses with important input from AS and MT. ML, JG, PD and LU produced the genomic dataset. AW, MT and LU wrote the manuscript, with input from all co-authors.

## Acknowledgements

The authors would like to thank the Kākāpō Recovery Team for their continued support, aid and interest in our ongoing microbiome research, and Genomics Aotearoa for providing a framework for collaborative research in the genomics space. The authors thank the Kākāpō125+ consortium and Ngāi Tahu for providing access to the kākāpō genomic dataset. The authors also wish to acknowledge the use of New Zealand eScience Infrastructure (NeSI) high performance computing facilities, consulting support and training services as part of this research. New Zealand’s national facilities are provided by NeSI and funded jointly by NeSI’s collaborator institutions and through the Ministry of Business, Innovation and Employment’s Research Infrastructure programme (https://www.nesi.org.nz). We would also like to acknowledge Prof Neil Gemmell and the Department of Anatomy, University of Otago, for their ongoing support.

## Declarations

Samples collected for this study were approved by the New Zealand Department of Conservation (Te Papa Atawhai) and did not require ethics approval from its Animal Ethics Committee under the New Zealand Animal Welfare Act.

## Availability of data and materials

The raw 16S rRNA gene amplicon sequence data are available in the NCBI SRA repository, under Bioproject accession number PRJNA859416. Metadata, bacterial taxonomic assignments, the non-rarefied ASV table, ASV data analyses and GWAS pipeline and analyses have been included as Additional files 2, 3, 4, 5 and 6 respectively. Genomic data are available via an application form at the Genomics Aotearoa Data Repository (https://repo.data.nesi.org.nz) as per the Global Indigenous Data Alliance guidelines.

Additional file 1: Supplementary material

Additional file 2: Sample metadata table

Additional file 3: Bacterial ASV taxonomic assignments

Additional file 4: Non-rarefied ASV table

Additional file 5: R studio ASV statistical analyses

Additional file 6: GWAS pipeline and analyses

