## Supplementary Figures (Additional file 1) for "Capturing species-wide diversity of the gut microbiota and its relationship with genomic variation in the critically endangered kākāpō"

Additional file 1: Supplementary Material

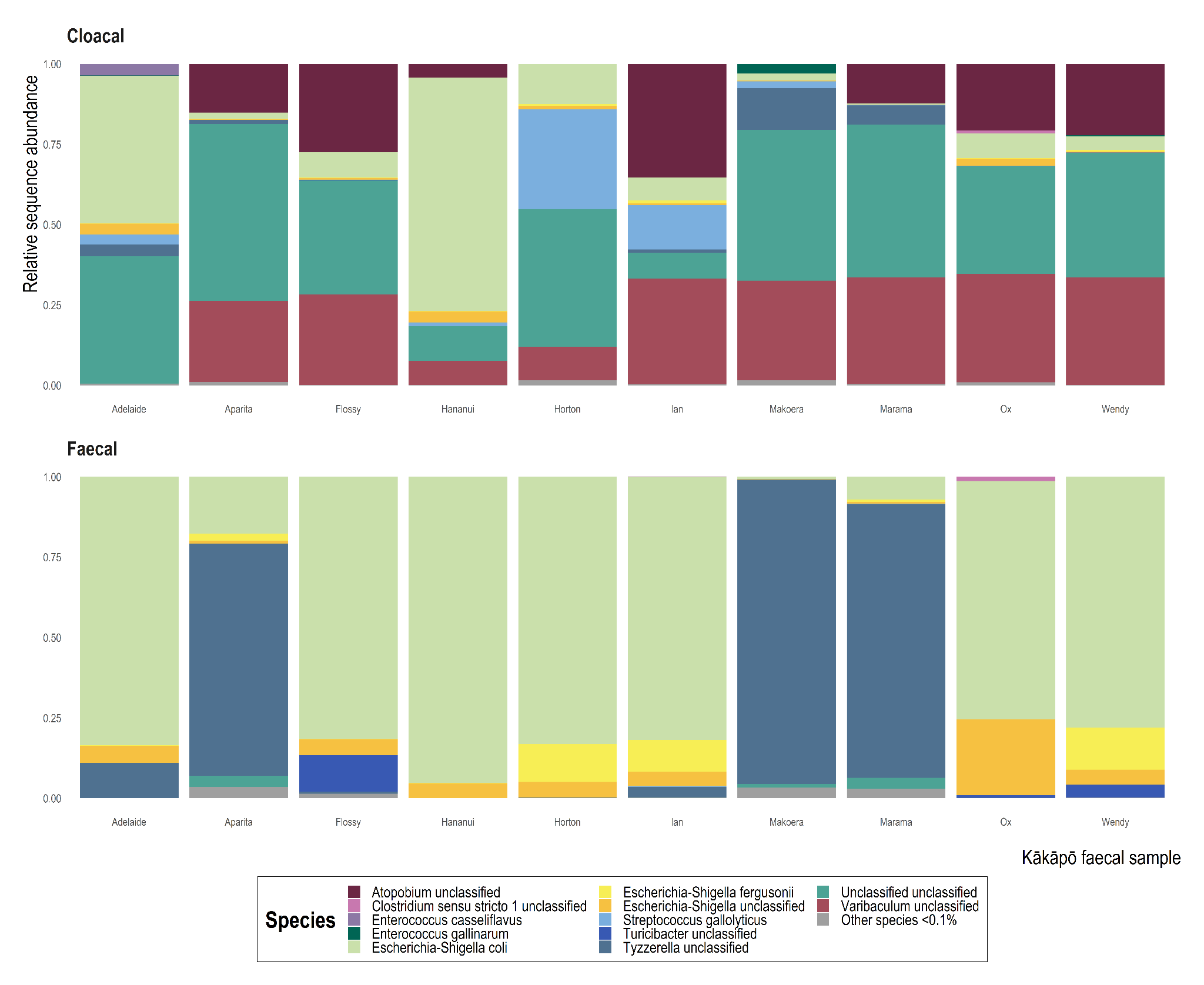

*Supplementary Figure 1: 16S rRNA gene sequence-based taxonomic distribution of bacteria within kākāpō cloacal swabs and faecal samples, demonstrating discordance between the two sample types. There are corresponding cloacal swabs and faecal samples obtained from the same kākāpō for 10 individuals in total. Each bar represents a single sample.*

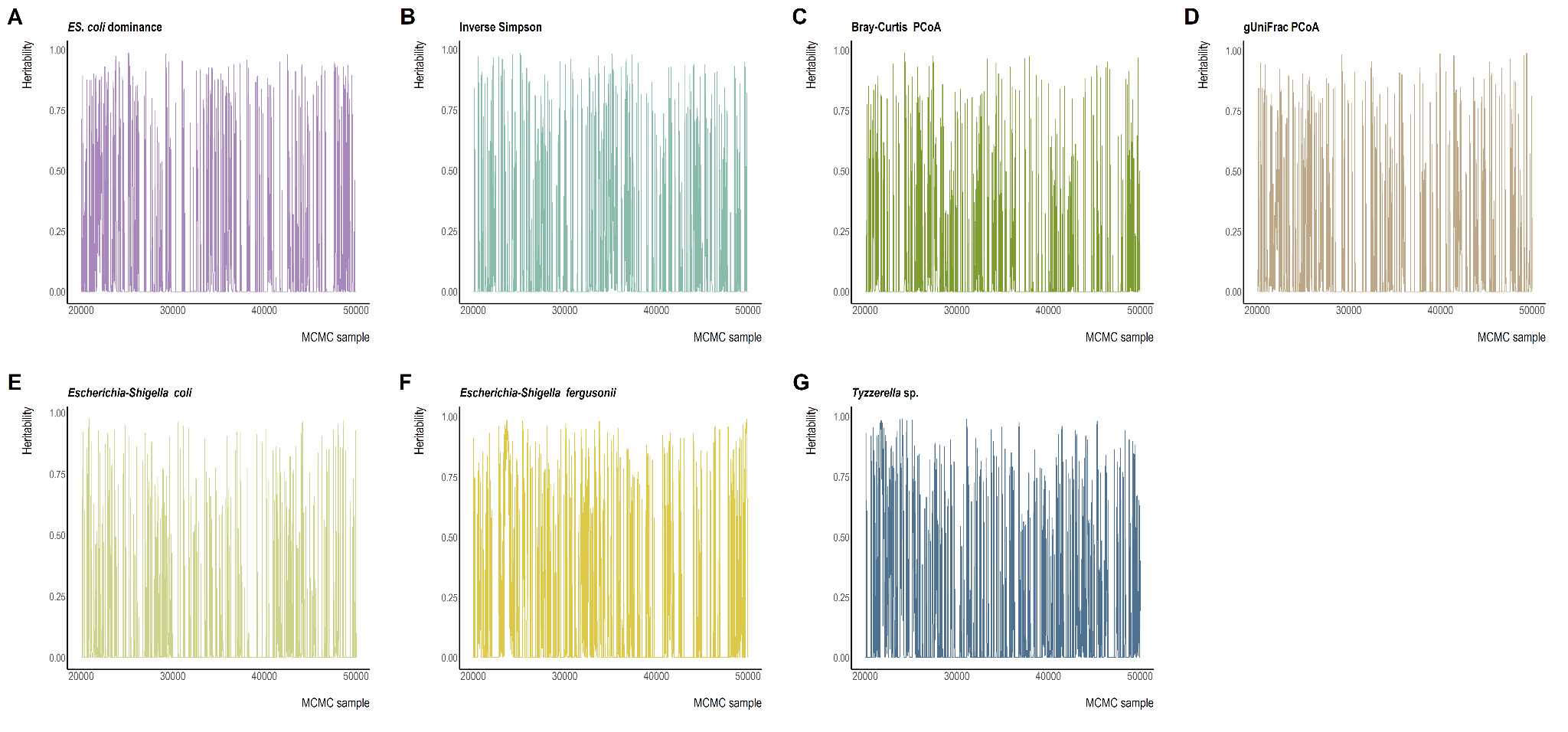

*Supplementary Figure 2: MCMC chain plots produced using Bayesian mixture modelling in BayesR for [A]* ES. coli *dominance [B] Inverse Simpson Diversity [C] Bray-Curtis PcoA diversity [D] gUniFrac PcoA diversity [E]* Escherichia-Shigella coli *relative abundance [F]* Escherichia-Shigella fergusonii *relative abundance and [G]* Tyzzerella *sp. relative abundance phenotypes.*

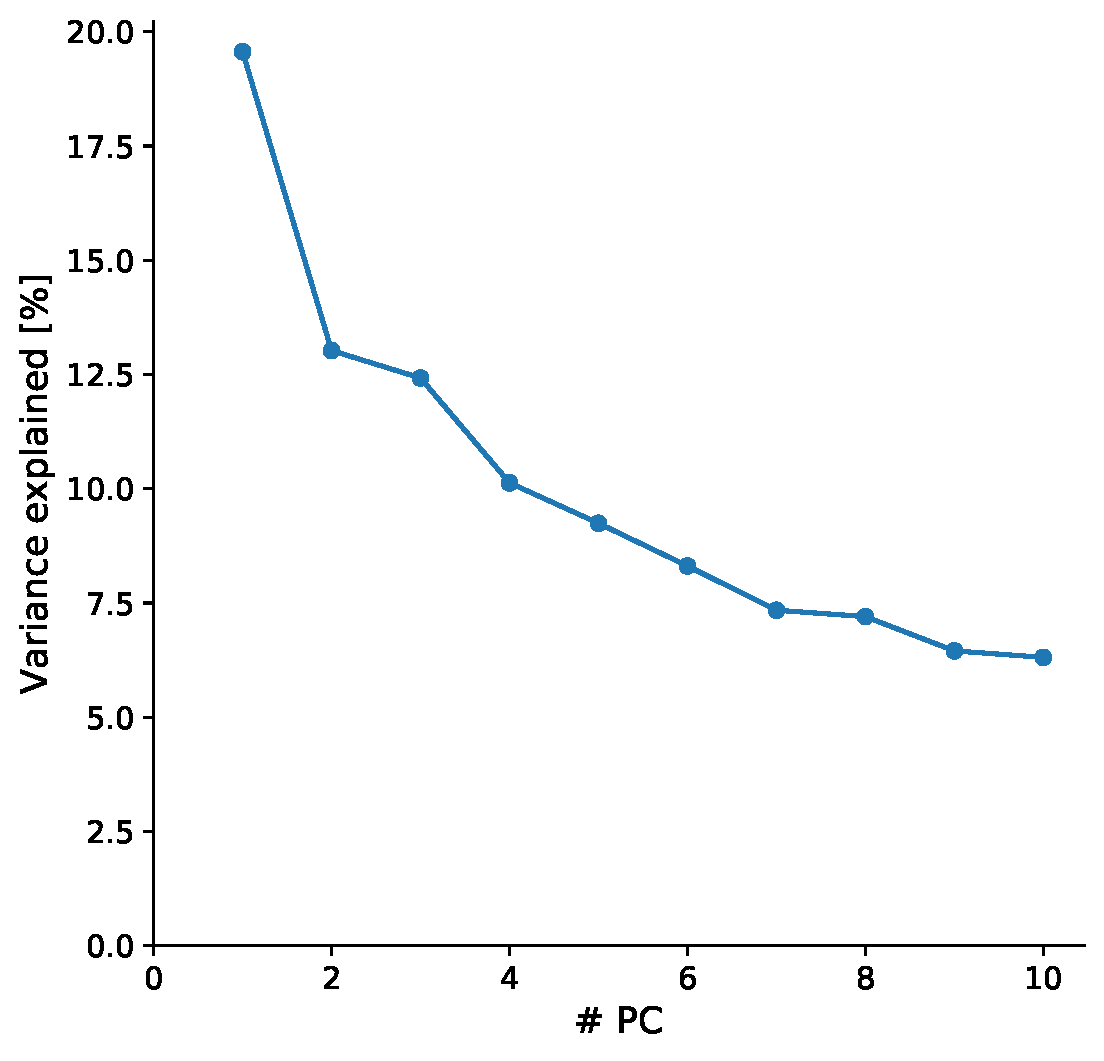

*Supplementary Figure 3: Scree plot of the principal components obtained from PCA of the variance-standardised relationship matrix (Genotype-phenotype analyses; Material and Methods).*

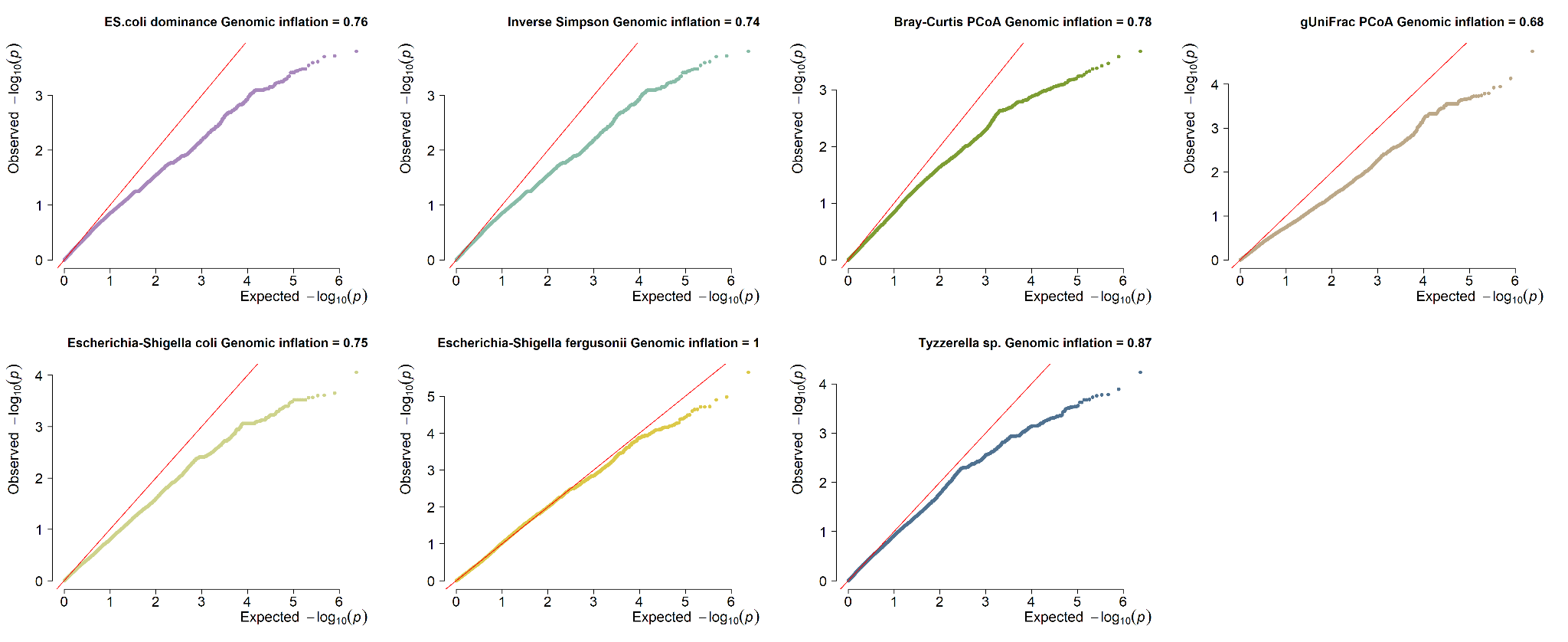

*Supplementary Figure 4: QQ plots based on RepeatABEL GWAS p-values for [A]* ES. coli *dominance [B] Inverse Simpson Diversity [C] Bray-Curtis PcoA diversity [D] gUniFrac PcoA diversity [E]* Escherichia-Shigella coli *relative abundance [F]* Escherichia-Shigella fergusonii *relative abundance and [G]* Tyzzerella *sp. relative abundance phenotypes. Genomic inflation of p-values listed for each phenotype.*

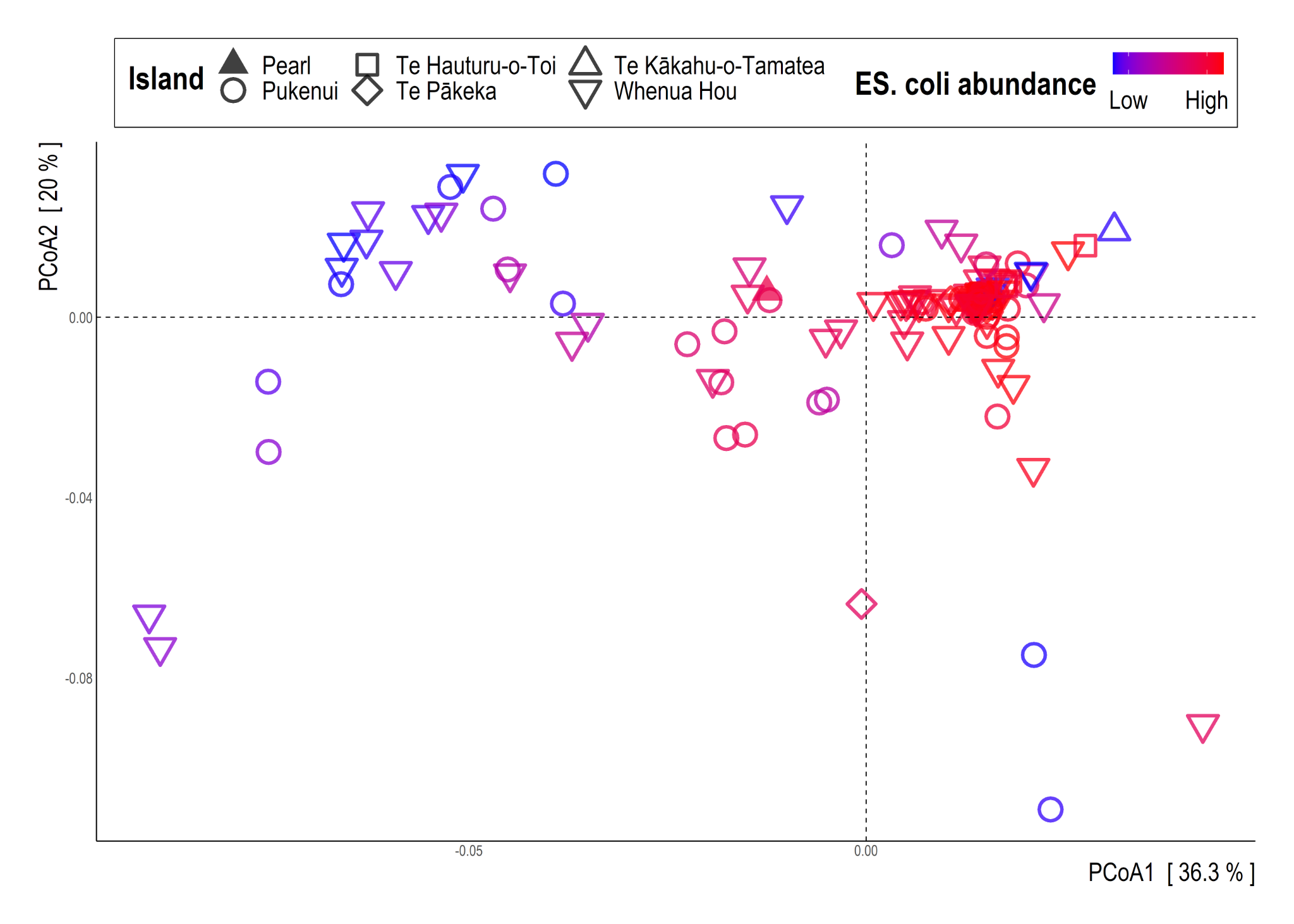

*Supplementary Figure 5: 16S rRNA gene sequence-based generalised UniFrac dissimilarity distances visualised via principal coordinate analysis ordination. Each dot of the PCoA represents the microbiota of a single kākāpō faecal sample. Samples are coloured by relative abundance of* Escherichia-Shigella coli *and shaped by island location.*

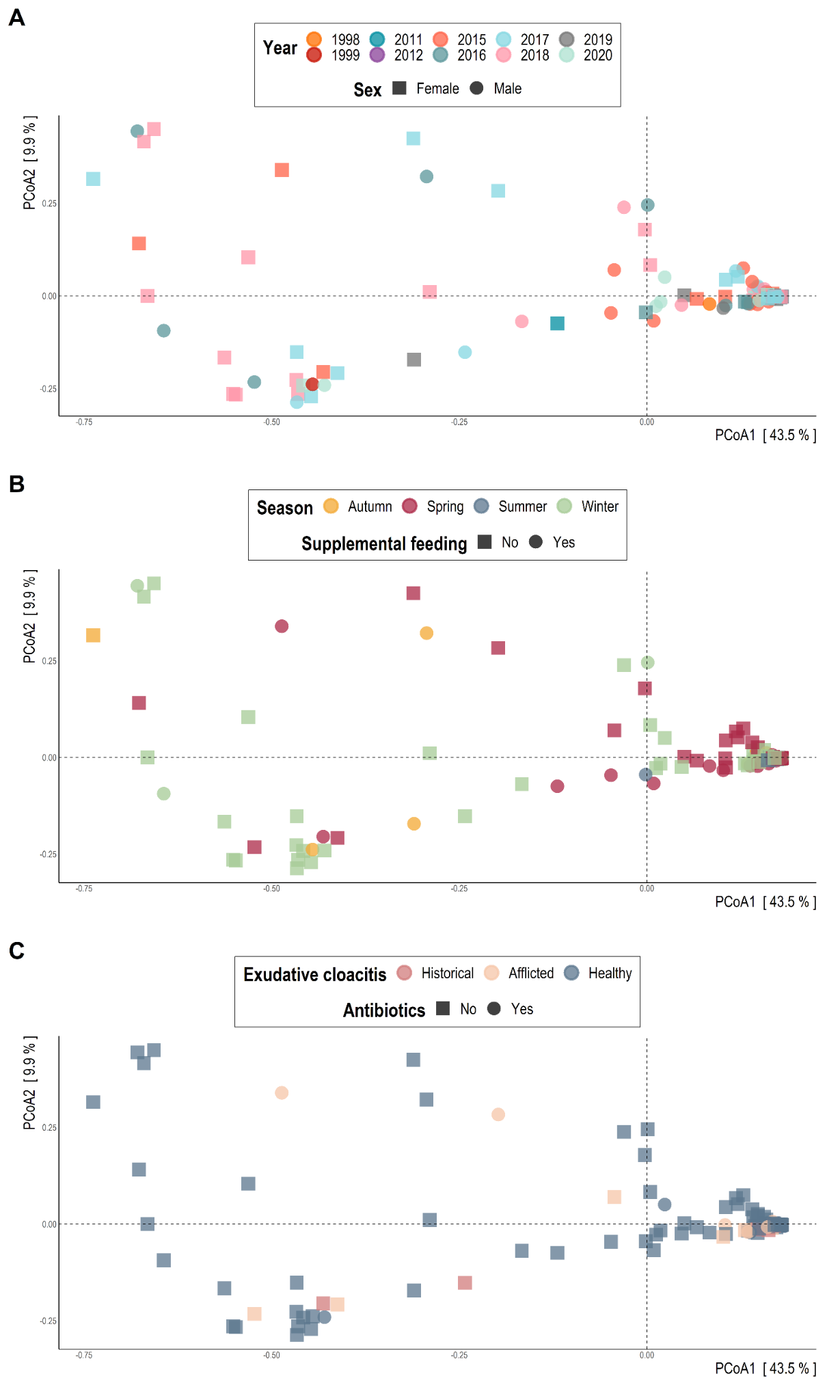

*Supplementary Figure 6: 16S rRNA gene sequence-based Bray-Curtis dissimilarity distances visualised via principal coordinate analysis ordination. Each dot of the PCoA represents the microbiota of a single kākāpō faecal sample. [A] samples are coloured by year of collection and shaped by kākāpō sex. [B] samples are coloured by season of collection and shaped by supplemental feeding. [C] samples are coloured by exudative cloacitis history and shaped by antibiotic treatment history.*

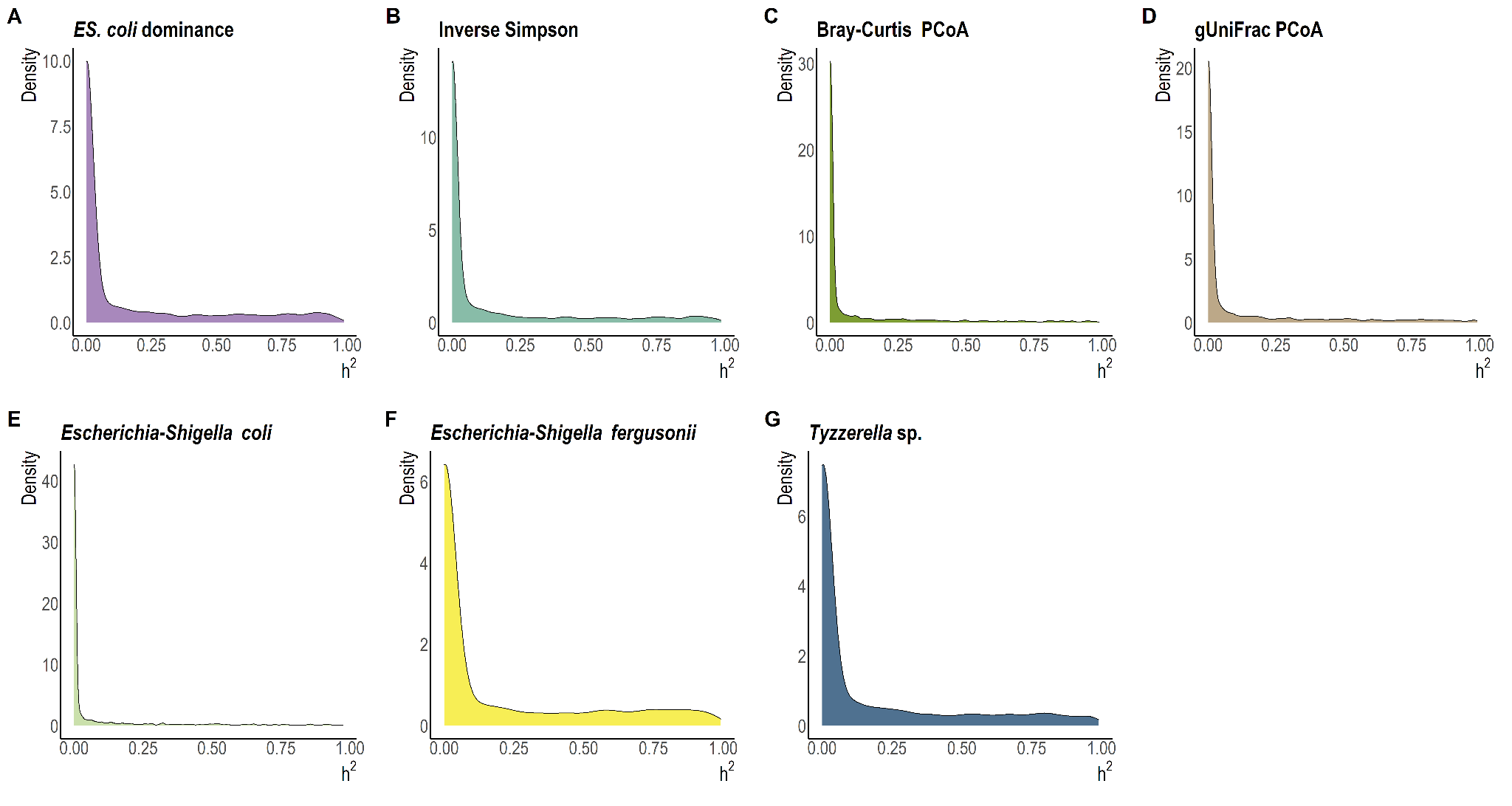

*Supplementary Figure 7: Density plots of the heritability estimate posterior mode per microbiota-related phenotype, as obtained from the MCMC analysis.*

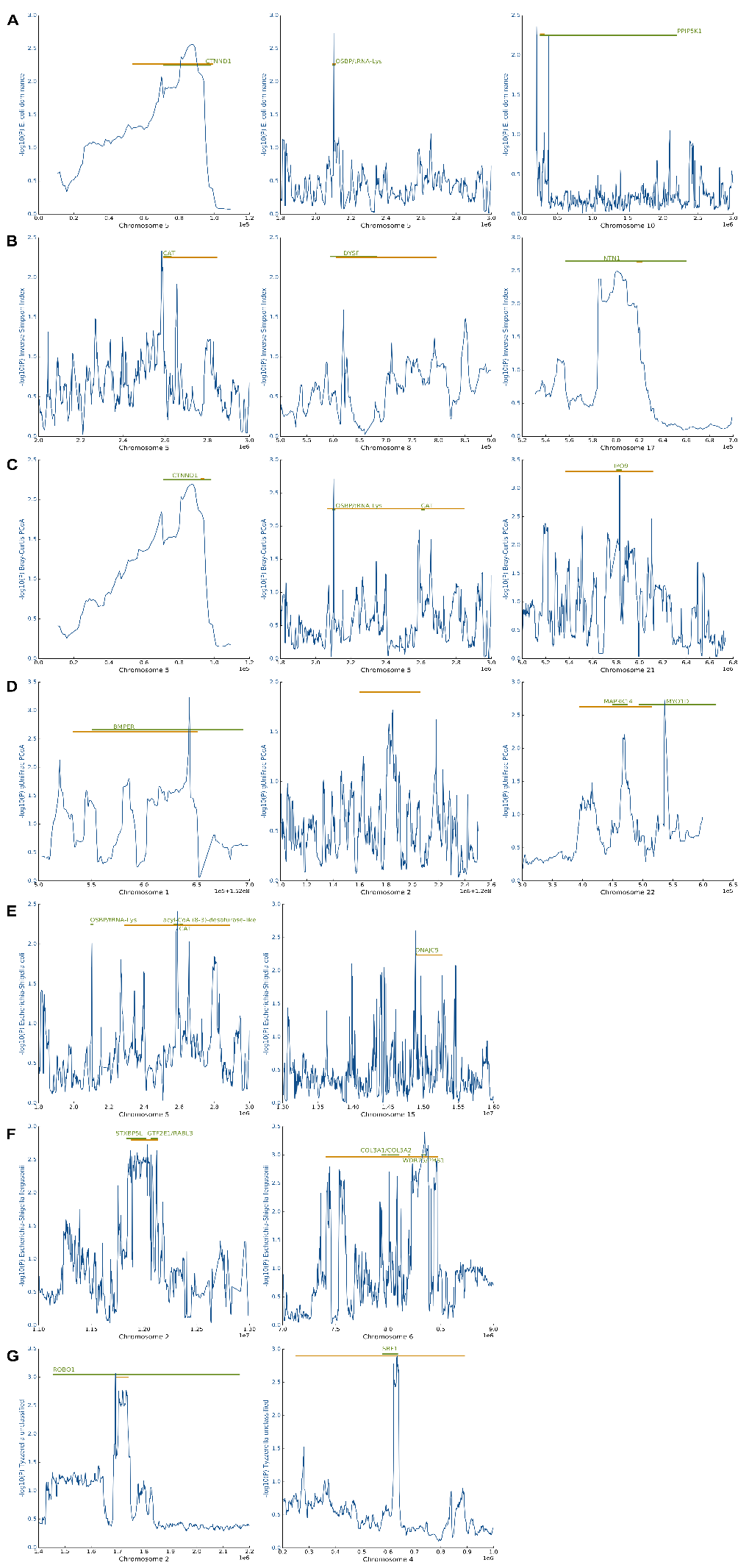

*Supplementary Figure 8: Sliding window plots where SNP-wise p-values are averaged to window-wise p-values in order to pinpoint the exact genomic region of nominally significant associations and to identify genes underlying these regions. Sliding window plots with significant genes identified for phenotypes [A]* ES. coli *dominance [B] Inverse Simpson diversity [C] Bray-Curtis PcoA [D] gUniFrac PcoA [E]* Escherichia-Shigella coli *relative abundance [F]* Escherichia-Shigella fergusonii *relative abundance [G]* Tyzzerella *sp. relative abundance. The orange bar represents the chromosome position range over which we identified significant SNPs, while the green bar indicates the length and position of an annotated gene within which our significant SNPs lie.*
