## Additional file 5 for "Capturing species-wide diversity of the gut microbiota and its relationship with genomic variation in the critically endangered kākāpō": R_statistical_analyses.html

Kakapo adult bacterial analyses


### Kakapo adult bacterial analyses

###### Annie G. West

Load packages into session, and print package version.

```
library(ggplot2); packageVersion("ggplot2")
library(Manu); packageVersion("Manu")
library(phyloseq); packageVersion("phyloseq")
library(gridExtra); packageVersion("gridExtra")
library(ggsignif); packageVersion("ggsignif")
library(ggrepel); packageVersion("ggrepel")
library(ggsci); packageVersion("ggsci")
library(hrbrthemes); packageVersion("hrbrthemes")
library(tidyverse); packageVersion("tidyverse")
library(viridis); packageVersion("viridis")
library(extrafont); packageVersion("extrafont")
set.seed(100)
```

### Import data

```
seqtab.nochim <- data.frame(readRDS("NeSI_adultK_output/16S_ASV_table_nochim.rds")) 
sum(rowSums(seqtab.nochim)) 

rownames(seqtab.nochim)[rownames(seqtab.nochim) == "B-MURPHY"] <- "B_MURPHY"
rownames(seqtab.nochim)[rownames(seqtab.nochim) == "DUSKY-J"] <- "DUSKY_J"
rownames(seqtab.nochim)[rownames(seqtab.nochim) == "EGILSAY-S"] <- "EGILSAY_S"
rownames(seqtab.nochim)[rownames(seqtab.nochim) == "KOTIU-J"] <- "KOTIU_J"
rownames(seqtab.nochim)[rownames(seqtab.nochim) == "TE-KINGI"] <- "TE_KINGI"

taxa <- readRDS("16S_taxa_table.rds")
Ptree <- readRDS("phylo_tree_adultK.rds")

map <- read.table("kakapo_adult_16S_map.txt",header = T, sep = '\t')
row.names(map) <- map$SampleID

# Make sure row names are the same between map and ASV table
identical(rownames(seqtab.nochim), rownames(map))
setdiff(rownames(seqtab.nochim), rownames(map))


seq.reads = data.frame(rowSums(seqtab.nochim))
map.reads = cbind(map, seq.reads)
rownames(map.reads) = map.reads$Name
```

```
taxa <- data.frame(taxa)
taxa$Family[is.na(taxa$Family)]="Unclassified"
taxa$Genus[is.na(taxa$Genus)]="Unclassified"
taxa$Species[is.na(taxa$Species)]="unclassified"
taxa$Taxonomy <- paste(taxa$Genus, taxa$Species, sep=" ")

taxmatrix <- as.matrix(taxa)
```

Create a phyloseq object for downstream analyses:

```
ps <- phyloseq(tax_table(taxmatrix),sample_data(map),
               otu_table(seqtab.nochim, taxa_are_rows = F), 
               phy_tree(Ptree$tree))
ps
```

### Filtering with phyloseq

##### Taxonomic filtering

Use to filter out non-target taxa e.g. mitochondria etc.

```
# Show available ranks in the dataset
rank_names(ps)

ps_tax_table <- as.data.frame(tax_table(ps))
sum(is.na(ps_tax_table$Phylum))
str(ps_tax_table$Phylum)
sapply(ps_tax_table, n_distinct)
levels(factor(ps_tax_table$Kingdom))

library(dplyr); packageVersion("dplyr")
count(ps_tax_table, Kingdom)

archaea = subset_taxa(ps, Kingdom == "Archaea")
archaea
sum(rowSums(data.frame(otu_table(archaea))))
```

```
ps_tax_table[is.na(ps_tax_table)]="Unclassified"
asv_table_ps<-as.data.frame(t(otu_table(ps)))
asv_table_wTax_ps <- cbind(asv_table_ps, ps_tax_table)

Phyla.sum.ps<-aggregate(asv_table_wTax_ps[,1:133], list(asv_table_wTax_ps$Phylum),sum)
row.names(Phyla.sum.ps)<- Phyla.sum.ps$Group.1
Phyla.sum.ps$Group.1 <- NULL
sum(rowSums(Phyla.sum.ps)) 
rowSums(Phyla.sum.ps != 0) #number of samples that phylum occurs in

P_rowsum.ps <- rowSums(Phyla.sum.ps)
P_rowsum.ps #714 reads unclassified
dir.create("tables")
write.csv(P_rowsum.ps, "tables/P_rowsum_ps.csv")

Class.sum.ps <- aggregate(asv_table_wTax_ps[,1:133], list(asv_table_wTax_ps$Class),sum)
row.names(Class.sum.ps)<- Class.sum.ps$Group.1
Class.sum.ps$Group.1 <- NULL
C_rowsum.ps <- rowSums(Class.sum.ps) 
write.csv(C_rowsum.ps, "tables/Class_sum_table_adult_kakapo_ps.csv")
write.csv(Class.sum.ps, "tables/Class_sum_per_sample_adult_kakapo_ps.csv")
```

```
# Create table, number of features for each phyla
table(tax_table(ps)[, "Phylum"], exclude = NULL)

ps0 <- subset_taxa(ps, !is.na(Phylum) & !Phylum %in% c("", "uncharacterized"))

phyla2Filter = c("<NA>")
ps1 = subset_taxa(ps0, !Phylum %in% phyla2Filter)
ps1

phyla2Filter = c("Chloroplast","Rickettsiales")
ps1 = subset_taxa(ps1, !Order %in% phyla2Filter)
ps1
rank_names(ps1)
table(tax_table(ps1)[,"Phylum"], exclude = NULL)
```

```
minTotRelAbun = 1e-6
x = taxa_sums(ps1)
keepTaxa = (x / sum(x)) > minTotRelAbun  
ps2 = prune_taxa(keepTaxa, ps1)
ps2

table(tax_table(ps2)[,"Phylum"], exclude = NULL)
```

### Rarefaction

```
#set.seed(100)
source('ggrarefaction_curves.R')
library(vegan)

rcurves <- ggrare(ps2, step = 500, se = FALSE)
rcurves_intcpt <- rcurves + geom_vline(xintercept=3480, color = "#CED38C") + theme_bw() +
  theme(panel.border = element_rect(color="black",size=0.7,linetype=1, fill = "NA"), 
        panel.background = element_blank(),
        axis.title.y = element_text(size = 12),
        axis.title.x = element_text(size=12)) +
  xlab("Sequence reads per faecal sample") + ylab("Richness") + ggtitle("Adult k\u101k\u101p\u14d faecal samples") +
  coord_cartesian(xlim = c(0, 50000))
rcurves_intcpt

ggsave("Rarefaction_curve_16S_adult_kakapo.png", rcurves_intcpt, height = 8, width = 12, bg = "white")


rASV <- rarefy_even_depth(otu_table(ps2, taxa_are_rows = F), sample.size = 3480,replace = FALSE)
# Save the rarefied ASV table
write.csv(rASV, "tables/rASV_table_adult_kakapo.csv")

Ntree <- phyloseq::phy_tree(ps2)
Ntaxa <- tax_table(ps2)
 
rASV.df = as.data.frame(rASV)
rASV.df = rASV.df[,order(colSums(rASV.df),decreasing = T)]
Ntaxa.df = as.data.frame(Ntaxa)

to.remove <- setdiff(rownames(Ntaxa.df), colnames(rASV.df))
Ntaxa.df = Ntaxa.df[!row.names(Ntaxa.df) %in% to.remove,]
setdiff(rownames(Ntaxa.df), colnames(rASV.df))

identical(rownames(Ntaxa.df), colnames(rASV.df))
Ntaxa.df = Ntaxa.df[colnames(rASV.df),]
identical(rownames(Ntaxa.df), colnames(rASV.df))

Ntaxa.df$ASV_ID <- paste("ASV_", 1:nrow(Ntaxa.df), sep="")
Ntaxa.df$concat = paste(Ntaxa.df$ASV_ID, Ntaxa.df$Taxonomy, sep = "_")
rownames(Ntaxa.df) = Ntaxa.df$concat
names(rASV.df) = rownames(Ntaxa.df)


Ntaxa.df$ASV_ID = NULL
Ntaxa.df = as.matrix(Ntaxa.df)

psRT <- phyloseq(rASV,
                sample_data(map),
                Ntaxa,
                Ntree)

psR <- phyloseq(otu_table(rASV.df, taxa_are_rows = F),
                sample_data(map),
                tax_table(Ntaxa.df))
ps2
psR

write.csv(cbind(t(rASV),as.data.frame(tax_table(psR))), "tables/rASV_wTax_adult_kakapo.csv")
RA_rASV <- transform_sample_counts(rASV, function(x) x/sum(x))
write.csv(cbind(t(RA_rASV),as.data.frame(tax_table(psR))), "tables/RA_rASV_wTax_adult_kakapo.csv") 

tax1 <- tax_table(ps2)
taxR <- tax_table(psR)

dplyr::setdiff(tax1,taxR)
```

##### Mantel test

```
library(vegan)

rASV.dist = vegan::vegdist(rASV, method = "bray")
rASV1.dist = vegan::vegdist(rASV1, method = "bray")
rASV2.dist = vegan::vegdist(rASV2, method = "bray")
rASV3.dist = vegan::vegdist(rASV3, method = "bray")
rASV4.dist = vegan::vegdist(rASV4, method = "bray")

mantel(rASV.dist, rASV1.dist, method = "spearman", permutations = 9999)

mantel(rASV.dist, rASV2.dist, method = "spearman", permutations = 9999)

mantel(rASV.dist, rASV3.dist, method = "spearman", permutations = 9999)

mantel(rASV.dist, rASV4.dist, method = "spearman", permutations = 9999)
```

### Summaries

```
ntaxa(psR)
nsamples(psR)
sample_names(psR)[1:5]
rank_names(psR)
sample_variables(psR)
otu_table(psR)[1:5, 1:5]
tax_table(psR)[1:5, 1:4]

asv_df <- t(otu_table(psR))
avg_ASV <- colSums(asv_df != 0)
write.csv(avg_ASV, "tables/avg_ASV.csv")

prev_asv_df <- otu_table(psR)
prev_ASV <- colSums(prev_asv_df != 0)
write.csv(prev_ASV, "tables/prev_ASV.csv") 

summary_data <- map %>%
    group_by(Island) %>%
    summarise(Count = n()) 
summary_data
```

```
#How many reads per Phylum to get % abundance

asv_table<-as.data.frame(t(otu_table(psR)))
taxo_table<-as.data.frame(tax_table(psR))
asv_table_wTax <- cbind(asv_table, taxo_table)

Phyla.sum<-aggregate(asv_table_wTax[,1:133], list(asv_table_wTax$Phylum),sum)
row.names(Phyla.sum)<- Phyla.sum$Group.1
Phyla.sum$Group.1 <- NULL
P_rowsum <- rowSums(Phyla.sum)
P_rowsum

write.csv(Phyla.sum, "tables/Phyla_sum_per_sample_adult_kakapo.csv")
write.csv(P_rowsum, "tables/Phyla_rowsum_adult_kakapo.csv")
```

```
Class.sum <- aggregate(asv_table_wTax[,1:133], list(asv_table_wTax$Class),sum)
row.names(Class.sum)<- Class.sum$Group.1
Class.sum$Group.1 <- NULL
C_rowsum <- rowSums(Class.sum) 
write.csv(C_rowsum, "tables/Class_sum_table_adult_kakapo.csv")
write.csv(Class.sum, "tables/Class_sum_per_sample_adult_kakapo.csv")
```

```
Order.sum <- aggregate(asv_table_wTax[,1:133], list(asv_table_wTax$Order),sum)
row.names(Order.sum)<- Order.sum$Group.1
Order.sum$Group.1 <- NULL
O_rowsum <- rowSums(Order.sum) 
write.csv(O_rowsum, "tables/Order_sum_table_adult_kakapo.csv")
write.csv(Order.sum, "tables/Order_sum_per_sample_adult_kakapo.csv")
```

```
Genus.sum <- aggregate(asv_table_wTax[,1:133], list(asv_table_wTax$Genus),sum)
row.names(Genus.sum)<- Genus.sum$Group.1
Genus.sum$Group.1 <- NULL
G_rowsum <- rowSums(Genus.sum) 
write.csv(G_rowsum, "tables/genus_sum_table_adult_kakapo.csv")
write.csv(Genus.sum, "tables/Genus_sum_per_sample_adult_kakapo.csv")
```

```
taxa.sum <- aggregate(asv_table_wTax[,1:133], list(asv_table_wTax$Taxonomy),sum)
row.names(taxa.sum)<- taxa.sum$Group.1
taxa.sum$Group.1 <- NULL
t_rowsum <- rowSums(taxa.sum) 
write.csv(t_rowsum, "tables/taxa_sum_table_adult_kakapo.csv")
write.csv(taxa.sum, "tables/taxa_sum_per_sample_adult_kakapo.csv")

taxa.df = rowSums(taxa.sum != 0)
write.csv(taxa.df, "tables/prev_taxa.csv")
```

### Taxa plots

```
taxaplot_theme <-  theme_ipsum() + theme(plot.title = element_text(size = 35),
                                         
          axis.text.x = element_blank(),

          axis.ticks.x = element_blank(),

          axis.text.y=element_text(size=28),

          axis.title.y=element_text(size=35),
          
          axis.title.x = element_text(size = 35),
          
          strip.text.x = element_text(size = 26),

          panel.grid.major = element_blank(),

          panel.grid.minor = element_blank(),

          panel.background = element_blank(),

          legend.box.background = element_rect(),

          legend.box.margin = margin(5, 5, 5, 5),

          legend.position="bottom",

          legend.title = element_text(face = "bold", size = 35),
          
          legend.text = element_text(size = 30))

dir.create("taxa_plots")
```

##### Phylum

```
library(plyr); packageVersion("plyr")
# get abundance in %
others <- transform_sample_counts(psR, function(x) x/sum(x))

# agglomerate taxa
glom <- tax_glom(others, taxrank = 'Phylum')

# create dataframe from phyloseq object
df <- psmelt(glom)

# convert Phylum to a character vector from a factor
df$Phylum <- as.character(df$Phylum)

# group dataframe by Phylum, calculate median rel. abundance
means <- ddply(df, ~Phylum, function(x) c(mean=mean(x$Abundance)))

# find Phyla whose rel. abund. is less than 1%
remainder <- means[means$mean <= 0.001,]$Phylum 

# change their name to "Others"
df[df$Phylum %in% remainder,]$Phylum <- 'Others <0.1%' 

print(levels(factor(df$Phylum)))
```

```
island.labs <- list("Pearl"="Pearl", "Pukenui"="Pukenui", "Te Hauturu-o-Toi"="Te Hauturu-o-Toi", "Te Pakeka"="Te P\u101keka", "Te Kakahu-o-Tamatea"="Te K\u101kahu-o-Tamatea", "Whenua Hou"="Whenua Hou")

df$Island <- factor(df$Island, levels = c("Whenua Hou","Pukenui","Te Hauturu-o-Toi","Te Kakahu-o-Tamatea","Te Pakeka","Pearl"), 
                  labels = c("WH", "PU", "TH", "TK", "TP", "PE"))
```

```
POplot <- ggplot(data=df, aes(x=Name, y=Abundance, fill = factor(Phylum, 
                                                                 levels =  c("Acidobacteriota","Actinobacteriota","Bacteroidota","Firmicutes","Proteobacteria","Verrucomicrobiota","Others <0.1%"))))


phylum_plot <- POplot + geom_bar(aes(), stat="identity", position="stack")  + 
  
  guides(fill=guide_legend(title="Phylum")) +
  
  taxaplot_theme +
  
  labs(x="K\u101k\u101p\u14d faecal sample", y="Relative sequence abundance") + 
  
  facet_wrap(~Island, scales = "free_x", nrow = 3) +
  
  scale_fill_futurama(alpha=0.75) 
  
phylum_plot
ggsave("taxa_plots/phylum_others_plot_adult_kakapo.png", plot = phylum_plot2, height = 15, width = 22,dpi = 300, bg = "white")
```

##### Species

```
glomS <- tax_glom(others, taxrank = 'Taxonomy')

dfS <- psmelt(glomS)

dfS$Taxonomy <- as.character(dfS$Taxonomy)

meansS <- ddply(dfS, ~Taxonomy, function(x) c(mean=mean(x$Abundance)))

remainderS1 <- meansS[meansS$mean <= 0.003,]$Taxonomy

# change their name to "Others"
dfS[dfS$Taxonomy %in% remainderS1,]$Taxonomy <- 'Other species <0.3%'

print(levels(factor(dfS$Taxonomy)))
```

```
taxa.colours.list.aug2021 <- c(
  "#402060","#D6C1DE","#CAE0AB","#4EB265","#F7EE55","#F6C141","#E8601C","#FF0F39","#FF99AB","#B1B1B1","#e2dfdf","#7BAFDE", "#ADE2D0BF","#4f7190","#9f9f9f")

dfS$Island <- factor(dfS$Island, levels = c("Whenua Hou","Pukenui","Te Hauturu-o-Toi","Te Kakahu-o-Tamatea","Te Pakeka","Pearl"), 
                  labels = c("WH", "PU", "TH", "TK", "TP", "PE"))

SOplot2 <- ggplot(data=dfS, aes(x=Name, y=Abundance, 
                                fill = factor(Taxonomy, levels =  c("Erwinia billingiae","Erwinia unclassified","Escherichia-Shigella coli",
                                                                    "Escherichia-Shigella dysenteriae","Escherichia-Shigella fergusonii",
                                                                    "Escherichia-Shigella unclassified","Klebsiella oxytoca","Lactobacillus gasseri",
                                                                    "Pantoea agglomerans","Pelosinus unclassified","Rahnella1 unclassified",
                                                                    "Streptococcus gallolyticus","Streptococcus unclassified",
                                                                    "Tyzzerella unclassified","Other species <0.3%"))))

Species_plot2 <- SOplot2 + ggtitle("Island") + geom_bar(aes(), stat="identity", position="fill")  +
  
  guides(fill=guide_legend(title="Species", nrow = 4)) + 
  
  taxaplot_theme + 
  
  labs(x="K\u101k\u101p\u14d faecal sample") + 
  
  facet_grid(~Island,  scales = "free_x", space = "free") +
  
  scale_fill_manual(values = taxa.colours.list.aug2021)

Species_plot2
ggsave("taxa_plots/Taxonomy_adult_kakapo_0.3.png", width = 22, height = 15, dpi=400, bg = "white")
```

###### Reordered by E.coli

```
library(dplyr)
library(tidyverse); packageVersion("tidyverse")
dfS %>% 
  group_by(Name) %>% 
  summarise(Sum = sum(Abundance)) %>% 
  arrange(Sum) %>% 
  print(n = Inf)

library(forcats); packageVersion("forcats")

dfS_reorder <- dfS 

dfS_reorder$Name_new <- 
  as.character(dfS_reorder$Name)

Name_new_levels <-
  dfS_reorder %>%
  filter(Taxonomy == "Escherichia-Shigella coli")  %>%
  group_by(Name_new)%>% 
  summarise(Sum = sum(Abundance)) %>%
  arrange(Sum) %>%
  pull(Name_new) %>%
  unique

dfS_reorder$Name_new <-
  factor(dfS_reorder$Name_new,
         levels = Name_new_levels)
```

```
Ecoli_SO <- ggplot(data=dfS_reorder, aes(x=Name_new, y=Abundance, 
                                        fill = factor(Taxonomy, levels =  c("Erwinia billingiae","Erwinia unclassified","Escherichia-Shigella coli",
                                                                    "Escherichia-Shigella dysenteriae","Escherichia-Shigella fergusonii",
                                                                    "Escherichia-Shigella unclassified","Klebsiella oxytoca","Lactobacillus gasseri",
                                                                    "Pantoea agglomerans","Pelosinus unclassified","Rahnella1 unclassified",
                                                                    "Streptococcus gallolyticus","Streptococcus unclassified",
                                                                    "Tyzzerella unclassified","Other species <0.3%"))))

Ecoli_SOplot <- Ecoli_SO + ggtitle("Island") + geom_col(aes(), position="fill")  +
  
  guides(fill=guide_legend(title="Species", nrow = 4)) + 
  
  taxaplot_theme + theme(strip.text.x = element_text(size = 23)) +
  
  labs(x="K\u101k\u101p\u14d faecal sample", y="Relative sequence abundance      ") + 
  
  facet_grid(~Island,  scales = "free_x", space = "free") + 
  
  scale_fill_manual(values = taxa.colours.list.aug2021)

Ecoli_SOplot
ggsave("taxa_plots/Taxonomy_Ecoli_ordered_island_adult_kakapo.png", width = 22, height = 15, dpi=400, bg = "white")
```

##### Core plot

```
core.colours <- c("#CAE0AB","#F7EE55","#4f7190")

core.taxa.df <- subset(dfS_reorder, Taxonomy == "Escherichia-Shigella coli" | Taxonomy == "Escherichia-Shigella fergusonii" | Taxonomy == "Tyzzerella unclassified")

core.taxa <- ggplot(data=core.taxa.df, aes(x=Name_new, y=Abundance, fill=Taxonomy))
core.taxa.plot <- core.taxa + geom_col(aes()) + 
  
  taxaplot_theme + 
  
  theme(legend.position = "none", strip.text.x = element_blank(), axis.title.x = element_blank()) +
  
  labs(y="Relative sequence abundance      ") + 
  
  facet_grid(~Island,  scales = "free_x", space = "free") + 
  
  scale_fill_manual(values = core.colours)

core.taxa.plot
ggsave("taxa_plots/core_taxonomy_Ecoli_ordered_island_adult_kakapo.png", width = 22, height = 15, dpi=400, bg = "white")
```

### GWAS rank normalise transformation

```
library(xavamess)

gwas_otu = data.frame(t(taxa.sum))
gwas_otu = gwas_otu[,order(colSums(gwas_otu),decreasing = T)]

e.coli.df = data.frame(gwas_otu$Escherichia.Shigella.coli)
rownames(e.coli.df) = rownames(gwas_otu)
rownames(e.coli.df) == rownames(map)
rownames(e.coli.df) = map$Name

tyz.df = data.frame(gwas_otu$Tyzzerella.unclassified)
rownames(tyz.df) = map$Name

eferg.df = data.frame(gwas_otu$Escherichia.Shigella.fergusonii)
rownames(eferg.df) = map$Name

gwas_taxa = list(e.coli.df, tyz.df$gwas_otu.Tyzzerella.unclassified, eferg.df$gwas_otu.Escherichia.Shigella.fergusonii)

gwas_taxa = data.frame(gwas_taxa)

names(gwas_taxa) = c("Escherichia_Shigella.coli","Tyzzerella.unclassified","Escherichia_Shigella.fergusonii")

hist(gwas_taxa$Escherichia_Shigella.fergusonii)

gwas_taxa_rank = gwas_taxa
gwas_taxa_rank$Escherichia_Shigella.coli = rank.normalize(gwas_taxa_rank[,1], FUN = qnorm)
gwas_taxa_rank$Tyzzerella.unclassified = rank.normalize(gwas_taxa_rank[,2], FUN = qnorm)
gwas_taxa_rank$Escherichia_Shigella.fergusonii = rank.normalize(gwas_taxa_rank[,3], FUN = qnorm)

hist(gwas_taxa_rank$Escherichia_Shigella.fergusonii)

write.table(gwas_taxa_rank,"gwas_taxa_df_ES_T_EF.csv", sep = '\t', row.names = T)
```

##### Test covariates

```
map_taxa <- map
map_taxa$Name == rownames(gwas_taxa)
map_taxa <- cbind(map_taxa, gwas_taxa)

library(dplyr)

#Island
map_taxa_is <- map_taxa[,c(3,18:20)]
map_taxa_is %>% 
      group_by(Island) %>% 
      summarise_each(funs(sum=sum(., na.rm=TRUE)))
map_taxa_is %>% group_by(Island) %>% summarize(count=n())

shapiro.test(map_taxa_is$Escherichia_Shigella.coli)
shapiro.test(map_taxa_is$Escherichia_Shigella.fergusonii)
shapiro.test(map_taxa_is$Tyzzerella.unclassified)

kruskal.test(map_taxa_is$Escherichia_Shigella.coli, map_taxa_is$Island)
kruskal.test(map_taxa_is$Escherichia_Shigella.fergusonii, map_taxa_is$Island)
kruskal.test(map_taxa_is$Tyzzerella.unclassified, map_taxa_is$Island)

library(dunn.test); packageVersion("dunn.test")
dunn.test(map_taxa_is$Escherichia_Shigella.coli, map_taxa_is$Island, method = "bh")
dunn.test(map_taxa_is$Tyzzerella.unclassified, map_taxa_is$Island, method = "bh")


#Sex
map_taxa_sex <- map_taxa[,c(15,18:20)]
map_taxa_sex %>% 
      group_by(sex) %>% 
      summarise_each(funs(sum=sum(., na.rm=TRUE)))
map_taxa_sex %>% group_by(sex) %>% summarize(count=n())

kruskal.test(map_taxa_sex$Escherichia_Shigella.coli, map_taxa_sex$sex)
kruskal.test(map_taxa_sex$Escherichia_Shigella.fergusonii, map_taxa_sex$sex) 
kruskal.test(map_taxa_sex$Tyzzerella.unclassified, map_taxa_sex$sex)


#Supplemental feeding
map_taxa_sf <- map_taxa[,c(7,18:20)]
map_taxa_sf %>% 
      group_by(Supplemental_Feeding) %>% 
      summarise_each(funs(sum=sum(., na.rm=TRUE)))
map_taxa_sf %>% group_by(Supplemental_Feeding) %>% summarize(count=n())

kruskal.test(map_taxa_sf$Escherichia_Shigella.coli, map_taxa_sf$Supplemental_Feeding)
kruskal.test(map_taxa_sf$Escherichia_Shigella.fergusonii, map_taxa_sf$Supplemental_Feeding)
kruskal.test(map_taxa_sf$Tyzzerella.unclassified, map_taxa_sf$Supplemental_Feeding) 


#Season
map_taxa_season <- map_taxa[,c(9,18:20)]
map_taxa_season %>% 
      group_by(Season) %>% 
      summarise_each(funs(sum=sum(., na.rm=TRUE)))
map_taxa_season %>% group_by(Season) %>% summarize(count=n())

kruskal.test(map_taxa_season$Escherichia_Shigella.coli, map_taxa_season$Season) 
kruskal.test(map_taxa_season$Escherichia_Shigella.fergusonii, map_taxa_season$Season)
kruskal.test(map_taxa_season$Tyzzerella.unclassified, map_taxa_season$Season) 

dunn.test(map_taxa_season$Escherichia_Shigella.coli, map_taxa_season$Season, method = "bh")
dunn.test(map_taxa_season$Tyzzerella.unclassified, map_taxa_season$Season, method = "bh")


#Year
map_taxa_year <- map_taxa[,c(4,18:20)]
map_taxa_year %>% 
      group_by(Year) %>% 
      summarise_each(funs(sum=sum(., na.rm=TRUE)))
map_taxa_year %>% group_by(Year) %>% summarize(count=n())

kruskal.test(map_taxa_year$Escherichia_Shigella.coli, map_taxa_year$Year)
kruskal.test(map_taxa_year$Escherichia_Shigella.fergusonii, map_taxa_year$Year)
kruskal.test(map_taxa_year$Tyzzerella.unclassified, map_taxa_year$Year)


#Hand rearing
map_taxa_hr <- map_taxa[,c(10,18:20)]
map_taxa_hr %>% 
      group_by(Hand_Rearing) %>% 
      summarise_each(funs(sum=sum(., na.rm=TRUE)))
map_taxa_hr %>% group_by(Hand_Rearing) %>% summarize(count=n())

kruskal.test(map_taxa_hr$Escherichia_Shigella.coli, map_taxa_hr$Hand_Rearing)
kruskal.test(map_taxa_hr$Escherichia_Shigella.fergusonii, map_taxa_hr$Hand_Rearing)
kruskal.test(map_taxa_hr$Tyzzerella.unclassified, map_taxa_hr$Hand_Rearing)


#Antibiotics
map_taxa_ab <- map_taxa[,c(16,18:20)]
map_taxa_ab %>% 
      group_by(Antibiotics) %>% 
      summarise_each(funs(sum=sum(., na.rm=TRUE)))
map_taxa_ab %>% group_by(Antibiotics) %>% summarize(count=n())

kruskal.test(map_taxa_ab$Escherichia_Shigella.coli, map_taxa_ab$Antibiotics)
kruskal.test(map_taxa_ab$Escherichia_Shigella.fergusonii, map_taxa_ab$Antibiotics)
kruskal.test(map_taxa_ab$Tyzzerella.unclassified, map_taxa_ab$Antibiotics)


#Exudative cloacitis
map_taxa_cb <- map_taxa[,17:20]
map_taxa_cb %>% 
      group_by(Crusty_Bum) %>% 
      summarise_each(funs(sum=sum(., na.rm=TRUE)))
map_taxa_cb %>% group_by(Crusty_Bum) %>% summarize(count=n())

kruskal.test(map_taxa_cb$Escherichia_Shigella.coli, map_taxa_cb$Crusty_Bum)
kruskal.test(map_taxa_cb$Escherichia_Shigella.fergusonii, map_taxa_cb$Crusty_Bum)
kruskal.test(map_taxa_cb$Tyzzerella.unclassified, map_taxa_cb$Crusty_Bum)
```

### Alpha Diversity

```
rich_psR <- estimate_richness(psR, measures = c("Observed", "InvSimpson"))
rich <- list(rich_psR, sample_data(psR)$Name, sample_data(psR)$Island,sample_data(psR)$Supplemental_Feeding,sample_data(psR)$Season,sample_data(psR)$Year,sample_data(psR)$Hand_Rearing,sample_data(psR)$sex,sample_data(psR)$Antibiotics,sample_data(psR)$Crusty_Bum)
names(rich) <- c("alpha_diversity", "Name", "Island","Supplemental_Feeding","Season","Year","Hand_Rearing","sex","Antibiotics","Crusty_Bum")

write.csv(rich, "alphadivR_kakapo_adults.csv")
rich_GWAS <- data.frame(rich)
rich_GWAS <- rich_GWAS[,c(4,1,3)]
names(rich_GWAS) <- c("Name","Observed","InvSimpson")
write.csv(rich_GWAS,"richR_GWAS_table.csv")
```

```
# Island
if (shapiro.test(rich$alpha_diversity[["Observed"]])$p.value > 0.05)
{print(summary(aov(rich$alpha_diversity[["Observed"]]~rich$Island, data = rich)))} else {print(kruskal.test(rich$alpha_diversity$Observed, rich$Island))}

if (shapiro.test(rich$alpha_diversity[["InvSimpson"]])$p.value > 0.05)
{print(summary(aov(rich$alpha_diversity[["InvSimpson"]]~rich$Island, data = rich)))} else {print(kruskal.test(rich$alpha_diversity$InvSimpson, rich$Island))} #0.026

# Sex
if (shapiro.test(rich$alpha_diversity[["Observed"]])$p.value > 0.05)
{print(summary(aov(rich$alpha_diversity[["Observed"]]~rich$sex, data = rich)))} else {print(kruskal.test(rich$alpha_diversity$Observed, rich$sex))} #0.016

if (shapiro.test(rich$alpha_diversity[["InvSimpson"]])$p.value > 0.05)
{print(summary(aov(rich$alpha_diversity[["InvSimpson"]]~rich$sex, data = rich)))} else {print(kruskal.test(rich$alpha_diversity$InvSimpson, rich$sex))}

# Supplemental Feeding
if (shapiro.test(rich$alpha_diversity[["Observed"]])$p.value > 0.05)
{print(summary(aov(rich$alpha_diversity[["Observed"]]~rich$Supplemental_Feeding, data = rich)))} else {print(kruskal.test(rich$alpha_diversity$Observed, rich$Supplemental_Feeding))}

if (shapiro.test(rich$alpha_diversity[["InvSimpson"]])$p.value > 0.05)
{print(summary(aov(rich$alpha_diversity[["InvSimpson"]]~rich$Supplemental_Feeding, data = rich)))} else {print(kruskal.test(rich$alpha_diversity$InvSimpson, rich$Supplemental_Feeding))}

# Season
if (shapiro.test(rich$alpha_diversity[["Observed"]])$p.value > 0.05)
{print(summary(aov(rich$alpha_diversity[["Observed"]]~rich$Season, data = rich)))} else {print(kruskal.test(rich$alpha_diversity$Observed, rich$Season))}

if (shapiro.test(rich$alpha_diversity[["InvSimpson"]])$p.value > 0.05)
{print(summary(aov(rich$alpha_diversity[["InvSimpson"]]~rich$Season, data = rich)))} else {print(kruskal.test(rich$alpha_diversity$InvSimpson, rich$Season))}

# Year
if (shapiro.test(rich$alpha_diversity[["Observed"]])$p.value > 0.05)
{print(summary(aov(rich$alpha_diversity[["Observed"]]~rich$Year, data = rich)))} else {print(kruskal.test(rich$alpha_diversity$Observed, rich$Year))}

if (shapiro.test(rich$alpha_diversity[["InvSimpson"]])$p.value > 0.05)
{print(summary(aov(rich$alpha_diversity[["InvSimpson"]]~rich$Year, data = rich)))} else {print(kruskal.test(rich$alpha_diversity$InvSimpson, rich$Year))}

# Hand Rearing
if (shapiro.test(rich$alpha_diversity[["Observed"]])$p.value > 0.05)
{print(summary(aov(rich$alpha_diversity[["Observed"]]~rich$Hand_Rearing, data = rich)))} else {print(kruskal.test(rich$alpha_diversity$Observed, rich$Hand_Rearing))}

if (shapiro.test(rich$alpha_diversity[["InvSimpson"]])$p.value > 0.05)
{print(summary(aov(rich$alpha_diversity[["InvSimpson"]]~rich$Hand_Rearing, data = rich)))} else {print(kruskal.test(rich$alpha_diversity$InvSimpson, rich$Hand_Rearing))}

# Antibiotics
if (shapiro.test(rich$alpha_diversity[["Observed"]])$p.value > 0.05)
{print(summary(aov(rich$alpha_diversity[["Observed"]]~rich$Antibiotics, data = rich)))} else {print(kruskal.test(rich$alpha_diversity$Observed, rich$Antibiotics))}

if (shapiro.test(rich$alpha_diversity[["InvSimpson"]])$p.value > 0.05)
{print(summary(aov(rich$alpha_diversity[["InvSimpson"]]~rich$Antibiotics, data = rich)))} else {print(kruskal.test(rich$alpha_diversity$InvSimpson, rich$Antibiotics))}

# Crusty Bum
if (shapiro.test(rich$alpha_diversity[["Observed"]])$p.value > 0.05)
{print(summary(aov(rich$alpha_diversity[["Observed"]]~rich$Crusty_Bum, data = rich)))} else {print(kruskal.test(rich$alpha_diversity$Observed, rich$Crusty_Bum))}

if (shapiro.test(rich$alpha_diversity[["InvSimpson"]])$p.value > 0.05)
{print(summary(aov(rich$alpha_diversity[["InvSimpson"]]~rich$Crusty_Bum, data = rich)))} else {print(kruskal.test(rich$alpha_diversity$InvSimpson, rich$Crusty_Bum))}
```

```
simpson_dunn <- data.frame(dunn.test(rich$alpha_diversity$InvSimpson, rich$Island, method = "bh"))
```

##### Bar plots

```
rich_df = data.frame(rich)
rich_df = data.frame(rich_df[,c(1:4,9)])
names(rich_df) = c("Observed","InvSimpson","Name","Island","Sex")
rich_df$Island <- factor(rich_df$Island, levels = c("Whenua Hou","Pukenui","Te Hauturu-o-Toi","Te Kakahu-o-Tamatea","Te Pakeka","Pearl"), 
                  labels = c("Whenua Hou", "Pukenui", "Te Hauturu-o-Toi", "Te K\u101kahu-o-Tamatea", "Te P\u101keka", "Pearl"))

rich_df$Name <-
  factor(rich_df$Name,
         levels = Name_new_levels)

kakapo_colours <- c("#4b5e1e", "#7D9D33", "#DCC949", "#BCA888", "#CD8862", "#775B24")

Observed_bar <- ggplot(data=rich_df, aes(x=Name,y=Observed, fill = Island)) + 
  
  geom_bar(aes(), stat="identity", position="stack")  + scale_y_log10() +
  
 theme_ipsum() + theme(axis.text.x = element_blank(),

          axis.ticks.x = element_blank(),

          axis.text.y=element_text(size=28),

          axis.title.y=element_text(size=35),
          
          axis.title.x = element_blank(),
          
          strip.text.x = element_blank(),

          panel.grid.major = element_blank(),

          panel.grid.minor = element_blank(),

          panel.background = element_blank(),

          legend.position = "none") +
  
  labs(y="Observed [log10]      ") + 
  
  facet_grid(~Island, scales = "free_x", space = "free")  + 
  
  scale_fill_manual(values = kakapo_colours)

Observed_bar

dir.create("alpha_div")
ggsave("alpha_div/Observed_barplot.png", width = 22, height = 15, dpi=400, bg = "white")
```

```
invsimp_bar <- ggplot(data=rich_df, aes(x=Name,y=InvSimpson, fill=Island)) + 
  
  geom_bar(aes(), stat="identity", position="stack")  + scale_y_log10() +
  
 theme_ipsum() + theme(axis.text.x = element_blank(),

          axis.ticks.x = element_blank(),

          axis.text.y=element_text(size=28),

          axis.title.y=element_text(size=35),
          
          axis.title.x = element_blank(),
          
          strip.text.x = element_blank(),

          panel.grid.major = element_blank(),

          panel.grid.minor = element_blank(),

          panel.background = element_blank(),

          legend.box.background = element_rect(),

          legend.box.margin = margin(5, 5, 5, 5),

          legend.position = "bottom",

          legend.title = element_text(face = "bold", size = 35),
          
          legend.text = element_text(size = 30)) +
  
  labs(y="Inverse Simpson [log10]      ") + 
  
  facet_grid(~Island, scales = "free_x", space = "free") + 
  
  scale_fill_manual(values = kakapo_colours) 

invsimp_bar

ggsave("alpha_div/invsimp_barplot.png", width = 22, height = 15, dpi=400, bg = "white")
```

```
myadptheme <- theme_ipsum() + theme(plot.title = element_text(size = 26),
  
          plot.subtitle = element_text(size = 24),
          
          axis.text.x = element_blank(),

          axis.title.x = element_text(size=35),

          axis.ticks.x = element_blank(),

          axis.text.y=element_text(size=25),

          axis.title.y=element_text(size=35),

          axis.line.x=element_line(color="black",size=1.0,linetype=1),

          axis.line.y=element_line(color="black",size=1.0,linetype=1),

          panel.grid.major = element_blank(),

          panel.grid.minor = element_blank(),

          panel.background = element_blank(),

          legend.box.background = element_rect(),

          legend.box.margin = margin(5, 5, 5, 5),

          legend.position="bottom",

          legend.title = element_text(face = "bold", size = 35),
          
          legend.text = element_text(size = 30))

sex.boxplot <- ggplot(rich_df, aes(x=Sex, y=Observed)) +
    geom_boxplot(size=1.0, aes(fill=Sex)) + 
    myadptheme + theme(plot.title = element_text(size = 35)) +
    scale_fill_manual(values = kakapo_colours) + 
    labs(y = "Observed richness", x = "K\u101k\u101p\u14d sex") +
    ggtitle("Wilcoxon p = 0.01")

sex.boxplot

ggsave("kakapo_sex_Observed_boxplot.png", sex.boxplot, height = 10, width = 10, dpi = 400, bg = "white")
```

### Beta Diversity

##### BrayCurtis PCoA

```
psR_RA <- transform_sample_counts(psR, function(x) {x/sum(x)})
psR_RA_table <- otu_table(psR_RA)
psR.dist <- vegdist(psR_RA_table, method="bray")
pcoa_bray<-cmdscale(psR.dist, k=2, eig=T)

library(bios2mds)
scree.plot(pcoa_bray$eig)

bray.pcoa.var.per<-round(pcoa_bray$eig/sum(pcoa_bray$eig)*100, 1)
bray.pcoa.values <- pcoa_bray$points

#make a table for GWAS using BrayCurtis pcoa points
write.csv(pcoa.values,"braycurtis_pcoa_values.csv")

bray.pcoa.data <- data.frame(Sample=rownames(bray.pcoa.values),
                        X=bray.pcoa.values[,1],
                        Y=bray.pcoa.values[,2])


bray.pcoa.data$Name <- map$Name
bray.pcoa.data$Year <- map$Year
bray.pcoa.data$Island <- map$Island
bray.pcoa.data$Supplemental_Feeding <- map$Supplemental_Feeding
bray.pcoa.data$E.coli_dominance <- map$R_E_abundance
bray.pcoa.data$Season <- map$Season
bray.pcoa.data$Cloacitis = map$Crusty_Bum
bray.pcoa.data$Antibiotics <- map$Antibiotics
bray.pcoa.data$Sex <- map$sex

bray.pcoa.data$Cloacitis = factor(bray.pcoa.data$Cloacitis, 
                                        levels = c("after", "yes","no"),
                                        labels = c("Historical","Afflicted","Healthy"))
bray.pcoa.data$Year = factor(bray.pcoa.data$Year)
bray.pcoa.data$Sex = factor(bray.pcoa.data$Sex)

dir.create("ordinations")
```

```
library(ggtext)
ord_theme = theme_ipsum() + 
  theme(plot.title = element_markdown(size = 35),
    
    axis.title.x = element_text(size=35),
    
    axis.title.y =element_text(size=35),

    axis.line.x=element_line(color="black",size=0.5,linetype=1),

    axis.line.y=element_line(color="black",size=0.5,linetype=1),

    panel.grid.major = element_blank(),

    panel.grid.minor = element_blank(),

    panel.background = element_blank(),

    legend.box.background = element_rect(),

    legend.box.margin = margin(5, 5, 5, 5),

    legend.text = element_markdown(size=25),
    
    legend.title = element_markdown(face="bold", size = 30),
    
    legend.position = "top")
```

```
bray.pcoa.coli <- ggplot(data=bray.pcoa.data, aes(x = X, y = Y, colour = E.coli_dominance)) +
  geom_point(size=7, stroke=2, alpha=0.75, aes(shape=Island)) +
  xlab(paste("PCoA1 ","[", bray.pcoa.var.per[1], "%", "]")) +
  ylab(paste("PCoA2 ", "[",bray.pcoa.var.per[2], "%", "]")) + 
  geom_vline(xintercept = 0, linetype = "dashed") + 
  geom_hline(yintercept = 0, linetype = "dashed") +
  
  scale_color_gradient(low = "blue", high = "red", name="*ES. coli* abundance",
                        labels=c("Low","High"),breaks=c(500,2950)) +
  scale_shape_manual(values=c(17,21:25),labels = c("Pearl", "Pukenui", "Te Hauturu-o-Toi", 
                                                            "Te P\u101keka", "Te K\u101kahu-o-Tamatea", "Whenua Hou")) +
  ord_theme +  guides(shape = guide_legend(order = 1)) 

bray.pcoa.coli

ggsave("bray_pcoa_coli.png", bray.pcoa.coli, height = 10, width = 17, bg = "white", dpi = 400)
```

###### Fig 2

```
library(cowplot); packageVersion("cowplot")

fig2_together = plot_grid(Ecoli_SOplot, core.taxa.plot, Observed_bar, invsimp_bar,  bray.pcoa.coli,labels = c("A","B","C","D","E"), label_size = 30, ncol=1, axis = c("lb"), rel_heights = c(2,1.3,0.8,1.2,1.9))
fig2_together
ggsave("Figure2.png", fig2_together, height = 40, width = 27, dpi=400, bg = "white")
```

```
f_colours2 <- c("#FF6F00BF", "#C71000BF", "#008EA0BF", "#8A4198BF", "#FF6348BF", "#5A9599BF","#84D7E1BF", "#FF95A8BF", "#767676", "#ADE2D0BF", "#1A5354BF", "#3F4041BF", "#FDE725FF","#D6C1DE")

year_sex = ggplot(data=bray.pcoa.data, aes(x = X, y = Y, colour =Year)) + 
  
  geom_point(size = 7, alpha=0.75, stroke=2, aes(shape=Sex)) +  
  
  xlab(paste("PCoA1 ","[", bray.pcoa.var.per[1], "%", "]")) +
  ylab(paste("PCoA2 ", "[",bray.pcoa.var.per[2], "%", "]")) + 
  
  geom_vline(xintercept = 0, linetype = "dashed") +
  geom_hline(yintercept = 0, linetype = "dashed") +
  
  ord_theme + theme(legend.box = "vertical") + 
  
  guides(color = guide_legend(order = 1)) +
    
  scale_shape_manual(values=c(15,16,21:25)) +
  
  scale_color_manual(values = f_colours2)  

year_sex
```

```
season.col <- c("#f3a833","#ac2847","#5b768d","#a6cb96")

season_sf <- ggplot(data=bray.pcoa.data, aes(x = X, y = Y, colour =Season)) + 
  
  geom_point(size = 7, alpha=0.75, stroke=2, aes(shape=Supplemental_Feeding)) +  
  
  xlab(paste("PCoA1 ","[", bray.pcoa.var.per[1], "%", "]")) +
  ylab(paste("PCoA2 ", "[",bray.pcoa.var.per[2], "%", "]")) + 
  
  geom_vline(xintercept = 0, linetype = "dashed") +
  geom_hline(yintercept = 0, linetype = "dashed") +
  
  ord_theme + theme(legend.box = "vertical") + 
  
  guides(color = guide_legend(order = 1)) +
    
  scale_shape_manual(values=c(15,16,21:25), labels = c("No","Yes"), name = "Supplemental feeding") +
  
  scale_color_manual(values = season.col) 

season_sf
```

```
crusty.col <- c("#d17c7c","#f6c6a8","#5b768d")

crusty_antib = ggplot(data = bray.pcoa.data, aes(x = X, y = Y, colour =Cloacitis)) + 
  geom_point(size = 7, alpha=0.75, stroke=2, aes(shape = Antibiotics)) +  
  
  xlab(paste("PCoA1 ","[", bray.pcoa.var.per[1], "%", "]")) +
  ylab(paste("PCoA2 ", "[",bray.pcoa.var.per[2], "%", "]")) + 
  
  geom_vline(xintercept = 0, linetype = "dashed") +
  geom_hline(yintercept = 0, linetype = "dashed") +
  
  ord_theme + theme(legend.box = "vertical") + 
  
  guides(color = guide_legend(order = 1)) +
    
  scale_shape_manual(values=c(15,16,21:25), labels = c("No", "Yes")) +
  
  scale_color_manual(values = crusty.col, name="Exudative cloacitis")

crusty_antib
```

```
season.col <- c("#f3a833","#ac2847","#5b768d","#a6cb96")

season_island <- ggplot(data=bray.pcoa.data, aes(x = X, y = Y, colour =Season)) + 
  
  geom_point(size = 7, alpha=0.75, stroke=2, aes(shape=Island)) +  
  
  xlab(paste("PCoA1 ","[", bray.pcoa.var.per[1], "%", "]")) +
  ylab(paste("PCoA2 ", "[",bray.pcoa.var.per[2], "%", "]")) + 
  
  geom_vline(xintercept = 0, linetype = "dashed") +
  geom_hline(yintercept = 0, linetype = "dashed") +
  
  ord_theme + theme(legend.box = "vertical") + 
  
  guides(color = guide_legend(order = 1)) +
    
  scale_shape_manual(values=c(15,16,21:25)) +
  
  scale_color_manual(values = season.col) 

season_island
```

```
year_island = ggplot(data=bray.pcoa.data, aes(x = X, y = Y, colour =Year)) + 
  
  geom_point(size = 7, alpha=0.75, stroke=2, aes(shape=Island)) +  
  
  xlab(paste("PCoA1 ","[", bray.pcoa.var.per[1], "%", "]")) +
  ylab(paste("PCoA2 ", "[",bray.pcoa.var.per[2], "%", "]")) + 
  
  geom_vline(xintercept = 0, linetype = "dashed") +
  geom_hline(yintercept = 0, linetype = "dashed") +
  
  ord_theme + theme(legend.box = "vertical") + 
  
  guides(color = guide_legend(order = 1)) +
    
  scale_shape_manual(values=c(15,16,21:25)) +
  
  scale_color_manual(values = f_colours2)  

year_island
```

```
year_season = ggplot(data=bray.pcoa.data, aes(x = X, y = Y, colour =Year)) + 
  
  geom_point(size = 7, alpha=0.75, stroke=2, aes(shape=Season)) +  
  
  xlab(paste("PCoA1 ","[", bray.pcoa.var.per[1], "%", "]")) +
  ylab(paste("PCoA2 ", "[",bray.pcoa.var.per[2], "%", "]")) + 
  
  geom_vline(xintercept = 0, linetype = "dashed") +
  geom_hline(yintercept = 0, linetype = "dashed") +
  
  ord_theme + theme(legend.box = "vertical") + 
  
  guides(color = guide_legend(order = 1)) +
    
  scale_shape_manual(values=c(15,16,21:25)) +
  
  scale_color_manual(values = f_colours2)  

year_season
```

```
suppF5 <- plot_grid(year_sex, season_sf, crusty_antib, labels = c("A","B","C"), label_size = 30, ncol=1, axis = c("lb"))
suppF5  
ggsave("SuppFig5.png", suppF5, height = 25, width = 15, dpi=400, bg = "white")

sampling_effects <- plot_grid(season_island, year_island, year_season,label_colour = c("A","B","C"), label_size = 30, ncol = 1, axis = c("lb"))
sampling_effects
ggsave("Sampling_effects_pcoa.png", sampling_effects, height = 25, width = 15, dpi = 400, bg = "white")
```

##### gUniFrac PCoA

```
library(GUniFrac); packageVersion("GUniFrac")
library(phangorn); packageVersion("phangorn")

rASV.unif <- as.data.frame(rASV)
gunifrac.tree <- midpoint(phy_tree(psRT))
gunifracs <- GUniFrac(rASV.unif, gunifrac.tree,  alpha=c(0,0.5,1))$unifracs

dw <- gunifracs[, , "d_1"]
du <- gunifracs[, , "d_UW"]
dv <- gunifracs[, , "d_VAW"]
d0 <- gunifracs[, , "d_0"]
d5 <- gunifracs[, , "d_0.5"]


##Use d0.5 > Chen et al. 2012
PermanovaG(gunifracs[, , c("d_0", "d_0.5", "d_1")] ~ map$Island)

d5.dist <- as.dist(d5)
pcoa_guniF<-cmdscale(d5, k=3, eig=T)

library(bios2mds)
scree.plot(pcoa_guniF$eig)

gpcoa.var.per<-round(pcoa_guniF$eig/sum(pcoa_guniF$eig)*100, 1)
gpcoa.values <- pcoa_guniF$points

#make a table for GWAS using gUniFrac pcoa points
write.csv(gpcoa.values,"GUniFrac_pcoa_values.csv")

gpcoa.data <- data.frame(Sample=rownames(gpcoa.values),
                        X=gpcoa.values[,1],
                        Y=gpcoa.values[,2],
                        Z=gpcoa.values[,3])

gpcoa.data$Name <- map$Name
gpcoa.data$Island <- map$Island
gpcoa.data$Supplemental_Feeding <- map$Supplemental_Feeding
gpcoa.data$E.coli_dominance <- map$R_E_abundance
gpcoa.data$Season <- map$Season
gpcoa.data$Cloacitis = map$Crusty_Bum
```

```
Ntaxa.df.unif <- as.data.frame(Ntaxa)
setdiff(rownames(Ntaxa.df.unif), colnames(rASV.unif))
to.remove <- setdiff(rownames(Ntaxa.df.unif), colnames(rASV.unif))
Ntaxa.df.unif = Ntaxa.df.unif[!row.names(Ntaxa.df.unif) %in% to.remove,]
setdiff(rownames(Ntaxa.df.unif), colnames(rASV.unif))

identical(rownames(Ntaxa.df.unif), colnames(rASV.unif))
rASV.unif = rASV.unif[,rownames(Ntaxa.df.unif)]

Ntaxa.df.unif$ASV_ID <- paste("ASV_", 1:nrow(Ntaxa.df.unif), sep="")
Ntaxa.df.unif$concat = paste(Ntaxa.df.unif$ASV_ID, Ntaxa.df.unif$Taxonomy, sep = "_")
rownames(Ntaxa.df.unif) = Ntaxa.df.unif$concat
names(rASV.unif) = rownames(Ntaxa.df.unif)


g.vec.sp<-envfit(pcoa_guniF$points , rASV.unif, perm=1000)

g.vec.sp.df<-
  as.data.frame(scores(g.vec.sp, "vectors"))

g.vec.sp.df$species<-rownames(g.vec.sp.df)


g.vec.sp.df.sig <- g.vec.sp.df[c(1:6), ]
```

```
evals2 = g.pcoa_guniF$eig
  
ggplot(data=gpcoa.data, aes(x = X, y = Y, colour = Island)) +
  geom_point(size=6, alpha=0.75, shape=17) +
  xlab(paste("PCoA1 ","[", gpcoa.var.per[1], "%", "]")) +
  ylab(paste("PCoA2 ", "[",gpcoa.var.per[2], "%", "]")) +
  geom_vline(xintercept = 0, linetype = "dashed") + 
  geom_hline(yintercept = 0, linetype = "dashed") +
  geom_segment(data=g.vec.sp.df.sig, 
               aes(x=0,xend=Dim1,y=0,yend=Dim2),
               arrow = arrow(length = unit(0.3, "cm")),
               colour="red", size = 1,
               inherit.aes=FALSE) + 
  geom_text(data=g.vec.sp.df.sig,
            aes(x=Dim1,y=Dim2,label=species),size=4.3,
            inherit.aes=FALSE) + 
  scale_color_manual(values = get_pal("Kakapo"),labels = c("Pearl", "Pukenui", "Te Hauturu-o-Toi", 
                                                            "Te P\u101keka", "Te K\u101kahu-o-Tamatea", "Whenua Hou")) + 
  ord_theme + xlim(-0.65, 0.77)
  
ggsave("ordinations/gpcoa_taxonomic_vectors_adult_kakapo.png",width=17, height = 10, dpi=400, bg = "white")
```

```
ggplot(data=gpcoa.data, aes(x = X, y = Y, colour = Island)) +
  geom_point(size=6, alpha=0.75, shape=17) +
  xlab(paste("PCoA1 ","[", gpcoa.var.per[1], "%", "]")) +
  ylab(paste("PCoA2 ", "[",gpcoa.var.per[2], "%", "]")) +
  geom_vline(xintercept = 0, linetype = "dashed") + 
  geom_hline(yintercept = 0, linetype = "dashed") +
  ord_theme +
  scale_color_manual(values = get_pal("Kakapo"),labels = c("Pearl", "Pukenui", "Te Hauturu-o-Toi", 
                                                            "Te P\u101keka", "Te K\u101kahu-o-Tamatea", "Whenua Hou"))
```

```
ecoli.gunifrac <- ggplot(data=gpcoa.data, aes(x = X, y = Y, colour = E.coli_dominance)) +
  geom_point(size=7, stroke=2, alpha=0.75, aes(shape=Island)) +
  xlab(paste("PCoA1 ","[", gpcoa.var.per[1], "%", "]")) +
  ylab(paste("PCoA2 ", "[",gpcoa.var.per[2], "%", "]")) + 
  geom_vline(xintercept = 0, linetype = "dashed") + 
  geom_hline(yintercept = 0, linetype = "dashed") +
  
  scale_color_gradient(low = "blue", high = "red", name="ES. coli abundance",
                        labels=c("Low","High"),breaks=c(500,2950)) +
  scale_shape_manual(values=c(17,21:25),labels = c("Pearl", "Pukenui", "Te Hauturu-o-Toi", 
                                                            "Te P\u101keka", "Te K\u101kahu-o-Tamatea", "Whenua Hou")) +
  ord_theme +  guides(shape = guide_legend(order = 1)) 

ecoli.gunifrac 

ggsave("SuppF4.png",ecoli.gunifrac,width=14, height = 10, dpi=400, bg = "white")
```

### PERMANOVA

##### Bray-Curtis

```
permanova.dis <- phyloseq::distance(psR, "bray")

vegan::adonis2(permanova.dis ~ Island, as(sample_data(psR), "data.frame"), permutations = 9999)

vegan::adonis2(permanova.dis ~ sex, as(sample_data(psR), "data.frame"), permutations = 9999)
 
vegan::adonis2(permanova.dis ~ Year, as(sample_data(psR), "data.frame"), permutations = 9999)

vegan::adonis2(permanova.dis ~ Hand_Rearing, as(sample_data(psR), "data.frame"), permutations = 9999)

vegan::adonis2(permanova.dis ~ Supplemental_Feeding, as(sample_data(psR), "data.frame"), permutations = 9999)

vegan::adonis2(permanova.dis ~ Season, as(sample_data(psR), "data.frame"), permutations = 9999)

vegan::adonis2(permanova.dis ~ Antibiotics, as(sample_data(psR), "data.frame"), permutations = 9999)

vegan::adonis2(permanova.dis ~ Crusty_Bum, as(sample_data(psR), "data.frame"), permutations = 9999)
```

```
vegan::adonis2(permanova.dis ~ sex + Island + Year + Hand_Rearing + Supplemental_Feeding + Season + Antibiotics + Crusty_Bum , as(sample_data(psR), "data.frame"), by="margin",permutations = 9999)
#                      Df SumOfSqs      R2      F Pr(>F)   
#sex                    1   0.6405 0.02845 3.9203 0.0058 **
#Island                 5   0.8613 0.03826 1.0544 0.3686   
#Year                   1   0.0937 0.00416 0.5732 0.7267   
#Hand_Rearing           1   0.2827 0.01256 1.7300 0.1078   
#Supplemental_Feeding   1   0.1300 0.00578 0.7958 0.5201   
#Season                 3   0.6933 0.03079 1.4143 0.1380   
#Antibiotics            1   0.0969 0.00430 0.5929 0.7143   
#Crusty_Bum             1   0.1782 0.00792 1.0907 0.3132   
#Residual             118  19.2797 0.85640                 
#Total                132  22.5126 1.00000  

#season dispersion not homogenous
```

Check that any significance detected in PERMANOVA is not due to
uneven dispersion between groups.

```
metadata = sample_data(psR)
beta <- vegan::betadisper(permanova.dis, metadata$Season)
beta
vegan::permutest(beta)

beta_boxplot <- boxplot(beta, xlab="Season") 
beta_boxplot
```

##### gUniFrac

```
vegan::adonis2(d5.dist ~ sex, data=map, permutations = 9999)
vegan::adonis2(d5.dist ~ Island, data=map, permutations = 9999)
vegan::adonis2(d5.dist ~ Year, data=map, permutations = 9999)
vegan::adonis2(d5.dist ~ Hand_Rearing, data=map, permutations = 9999)
vegan::adonis2(d5.dist ~ Supplemental_Feeding, data=map, permutations = 9999)
vegan::adonis2(d5.dist ~ Season, data=map, permutations = 9999)
vegan::adonis2(d5.dist ~ Antibiotics, data=map, permutations = 9999)
vegan::adonis2(d5.dist ~ Crusty_Bum, data=map, permutations = 9999)
```

```
vegan::adonis2(d5.dist ~ sex + Island + Year + Hand_Rearing + Supplemental_Feeding + Season + Antibiotics + Crusty_Bum , data=map, by="margin",permutations = 9999)
#                      Df SumOfSqs      R2      F Pr(>F)  
#sex                    1 0.006387 0.02283 3.1010 0.0120 *
#Island                 5 0.009883 0.03532 0.9597 0.4749  
#Year                   1 0.003496 0.01249 1.6973 0.1314  
#Hand_Rearing           1 0.002050 0.00733 0.9954 0.3901  
#Supplemental_Feeding   1 0.003714 0.01327 1.8032 0.0955 .
#Season                 3 0.004088 0.01461 0.6616 0.8290  
#Antibiotics            1 0.001965 0.00702 0.9540 0.4000  
#Crusty_Bum             1 0.002116 0.00756 1.0275 0.3690  
#Residual             118 0.243041 0.86867                
#Total                132 0.279784 1.00000
```

### Differential abundance

```
library(plyr); packageVersion("plyr")
library(dplyr); packageVersion("dplyr")
library(microbiome); packageVersion("microbiome")
library(DESeq2); packageVersion("DESeq2")

my_theme <- theme_ipsum() + theme(panel.grid.major = element_blank(), panel.grid.minor = element_blank(), axis.text.y = element_text(face="italic"))

ps2.ASV.df = as.data.frame(otu_table(ps2))
ps2.ASV.df = ps2.ASV.df[,order(colSums(ps2.ASV.df),decreasing = T)]
ps2.taxa.df = as.data.frame(tax_table(ps2))

identical(rownames(ps2.taxa.df), colnames(ps2.ASV.df))
setdiff(rownames(ps2.taxa.df), colnames(ps2.ASV.df))
ps2.taxa.df = ps2.taxa.df[colnames(ps2.ASV.df),]

ps2.taxa.df$ASV_ID <- paste("ASV_", 1:nrow(ps2.taxa.df), sep="")
ps2.taxa.df$concat = paste(ps2.taxa.df$ASV_ID, ps2.taxa.df$Taxonomy, sep = "_")
rownames(ps2.taxa.df) = ps2.taxa.df$concat
names(ps2.ASV.df) = rownames(ps2.taxa.df)

ps2.taxa.df$ASV_ID = NULL
ps2.taxa.df$concat = NULL
ps2.taxa.df = as.matrix(ps2.taxa.df)

ps2_edit <- phyloseq(otu_table(ps2.ASV.df, taxa_are_rows = F),
                     tax_table(ps2.taxa.df),
                     sample_data(map))
ps2_edit
```

```
ps.sta <- ps2_edit
ps.st0 = filter_taxa(ps.sta, function(x) sum(x > 3) > (0.05*length(x)), TRUE)
ps.st1 <- ps.st0
```

```
meta.st <- meta(ps.st0)
meta.st$sex <- as.factor(meta.st$sex)
diagdds_st = phyloseq_to_deseq2(ps.st1, ~ sex)
 gm_mean = function(x, na.rm=TRUE){
     exp(sum(log(x[x > 0]), na.rm=na.rm) / length(x))
 }
 geoMeans = apply(counts(diagdds_st), 1, gm_mean)
 diagdds_st = estimateSizeFactors(diagdds_st, geoMeans = geoMeans)
 dds_st = DESeq(diagdds_st, test="Wald", fitType="local")
```

```
otu.ab1 <- abundances(ps.st1)

res1 = results(dds_st, cooksCutoff = FALSE)
res_tax1 = cbind(as.data.frame(res1), as.matrix(rownames(otu.ab1)), OTU = rownames(res1))
res_tax1 = cbind(as(res_tax1, "data.frame"), as(tax_table(ps.st1)[rownames(res_tax1), ], "matrix"))
res_tax_sig1 = subset(res_tax1, padj < 0.05 & 0 < abs(log2FoldChange))
res_tax1$Significant <- ifelse(rownames(res_tax1) %in% rownames(res_tax_sig1) , "Yes", "No")
res_tax1$Significant[is.na(res_tax1$Significant)] <- "No"
sig_res1 <- res_tax1[rownames(res_tax_sig1),"OTU"]
res_table1 <- data.frame(res_tax_sig1$baseMean , res_tax_sig1$log2FoldChange,res_tax_sig1$padj)
row.names(res_table1) <- rownames(res_tax_sig1)

data_to_write1 <-res_tax_sig1[,c("baseMean","log2FoldChange","pvalue","padj","Phylum", "Class", "Order", "Family", "Genus","Species","Taxonomy","OTU")]
data_to_write1$DifferentiallyAbundant <-levels(meta.st[,"sex"])[as.numeric(data_to_write1$log2FoldChange>0)+1]

# Total numer of OTUs DA
nrow(data_to_write1) #differentially abundant ASVs
length(unique(data_to_write1$Genus)) #how many genera
length(unique(data_to_write1$Taxonomy)) #how many species
write.csv(data_to_write1,"tables/deseq_comparison_sex.csv")
df1 <- mutate(data_to_write1, Taxonomy, Taxonomy= paste(data_to_write1$Taxonomy ))
```

```
sex.diff.asvs <- ggplot(df1, aes(log2FoldChange, Taxonomy)) + 
  geom_point(aes(color = DifferentiallyAbundant), shape = 21, size = 3, stroke = 2) + 
  scale_color_manual(values= c("#7D9D33","#CED38C")) + my_theme + 
  theme(axis.text.y = element_text(face="italic"),
        panel.border = element_rect(color="black",size=0.7,linetype=1, fill = "NA"), 
        panel.background = element_blank(),
        axis.title.y = element_text(size = 16),
        axis.title.x = element_text(size=16),
        legend.box.background = element_rect(),
        legend.box.margin = margin(5, 5, 5, 5),
        legend.title = element_text(face = "bold", size = 15),
        legend.text = element_text(size = 13)) +  geom_vline(xintercept = 0) 
sex.diff.asvs 
ggsave("Deseq_sex_plot.png", width = 10, height = 5, dpi=400)
```
