## Additional file 6 for "Capturing species-wide diversity of the gut microbiota and its relationship with genomic variation in the critically endangered kākāpō": GWAS_microbiome_pipeline.html

GWAS code for R and NeSI


### GWAS code for R and NeSI

###### Annie G. West

### 1. Import mapping file and create subset\_mic file

```
setwd("./GWAS_final/")
maps = read.csv('GWAS_final/kakapo_adult_16S_map.txt', sep='\t')
rownames(maps) = maps$Name


alpha.div <- read.csv("GWAS_final/richR_GWAS_table.csv", header = T, row.names = 2)
rownames(alpha.div)==row.names(maps)
alpha.div = alpha.div[rownames(maps),]
rownames(alpha.div) == rownames(maps)
maps$Observed <- alpha.div$Observed
maps$Shannon <- alpha.div$Shannon
maps$InvSimpson <- alpha.div$InvSimpson
```

```
subset = data.frame(FID = rownames(maps), IID = rownames(maps),
                    stringsAsFactors = F)
library('dplyr')


for (i in 1:dim(subset)[1]){
  if (length(strsplit(subset$FID[i],' ')[[1]])==2){
    subset$FID[i] = strsplit(subset$FID[i],' ')[[1]][1]
    subset$IID[i] = strsplit(subset$IID[i],' ')[[1]][2]
  }
}

write.table(subset,"GWAS_final/subset_mic.tsv", sep = ' ', row.names = F, col.names = F)
```

### 2. Create genomic dataset and obtain genomic PCs in NeSI

```
module load PLINK/1.09b6.16
  
#Filtering down to microbiome data set:

plink --bcf /WORKINGDIRECTORY/05_GWAS_herit_partitioning/plink_final/Trained_nr.bcf --make-bed --keep ./GWAS/subset_mic.tsv  --chr-set 89 --maf 0.05 --geno 0.2 --hwe 0.0000001 --snps-only just-acgt --keep-allele-order --biallelic-only strict --vcf-filter --out ./genomic_data/plink_f_nr_mic

#Identifying variants we want to filter for LD:
  
plink --bfile ./genomic_data/plink_f_nr_mic --indep-pairwise 50 10 0.8 --chr-set 89 --out ./genomic_data/LD_nr --noweb

#Filter for LD and exclude:
  
plink --bfile ./genomic_data/plink_f_nr_mic  --exclude ./genomic_data/LD_nr.prune.out --chr-set 89 --noweb --make-bed --out         ./genomic_data/plink_f_nr_mic_LD

#PCA on LD filtered >>pcs from relatonship matrix:

plink --bfile ./genomic_data/plink_f_nr_mic_LD --pca var-wts 10 --chr-set 89 --out ./genomic_data/pca
```

### 3. Create covariate file to control for when running GWAS

Download the eigenvector file

```
pcs <- read.csv('pca.eigenvec', sep=' ', header = F)
colnames(pcs) <- c('FID', 'IID', 'PC1', 'PC2', 'PC3', 'PC4', 'PC5', 'PC6', 'PC7', 'PC8', 'PC9', 'PC10')

pcs$concat = pcs$FID
for (i in 1:dim(pcs)[1]){
  if (pcs$FID[i] != pcs$IID[i]){
    pcs$concat[i] = paste(pcs$FID[i],pcs$IID[i])
  }
}

rownames(maps) == pcs$concat
setdiff(rownames(maps), pcs$concat)
maps = maps[pcs$concat,]

pcs$concat = NULL

pcs$Sex <- maps$sex

pcs = pcs[,c('FID', 'IID', 'PC1', 'PC2', 'PC3', 'Sex')]

screeplot(pcs[,3:12])

write.table(pcs,"GWAS_final/cov.tsv", sep = ' ', row.names = F)
```

### 4. Create phenotype files

```
library(xavamess)

#E.coli dominance
phe.dom = pcs[,c('FID','IID')]
phe.dom$E.coli_dominant = maps$E.coli_dominant
phe.dom$pheno <- 1
phe.dom$pheno[which(maps$E.coli_dominant=='yes')] = 2

phe.dom.t <- data.frame(phe.dom[,c('FID','IID','pheno')])
dir.create("GWAS_final/phenotype_files")
write.table(phe.dom.t, "GWAS_final/phenotype_files/dominance.tsv",sep=' ', quote=FALSE, row.names = F,na = "NA")
```

```
#Import BrayCurtis pcoa axis PCs  
pcoa <- read.csv("braycurtis_pcoa_values.csv", row.names = 1)

rownames(pcoa) == rownames(maps)
pcoa = pcoa[rownames(maps),]
identical(rownames(pcoa),rownames(maps))

phe.pcoa = pcs[,c('FID','IID')]
phe.pcoa$pheno = pcoa$V1
hist(phe.pcoa$pheno)
phe.pcoa$pheno = rank.normalize(phe.pcoa[,3], FUN = qnorm)
hist(phe.pcoa$pheno)

phe.pcoa.t <- data.frame(phe.pcoa[,c('FID','IID','pheno')])
write.table(phe.pcoa.t, "GWAS_final/phenotype_files/pcoa.tsv",sep=' ', quote=FALSE, row.names = F,na = "NA")
```

```
#Import GUniFrac pcoa axis PCs 
gunifrac_pcoa <- read.csv("GUniFrac_pcoa_values.csv", row.names = 1)
rownames(gunifrac_pcoa) == maps$SampleID
gunifrac_pcoa = gunifrac_pcoa[maps$SampleID,]
identical(rownames(gunifrac_pcoa), maps$SampleID)

phe.gunifrac <- pcs[,c('FID','IID')]
phe.gunifrac$pheno = gunifrac_pcoa$V1
hist(phe.gunifrac$pheno)
phe.gunifrac$pheno = rank.normalize(phe.gunifrac[,3], FUN = qnorm)
hist(phe.gunifrac$pheno)

phe.gunifrac.t <- data.frame(phe.gunifrac[,c('FID','IID','pheno')])
write.table(phe.gunifrac.t, "GWAS_final/phenotype_files/gunifrac_pcoa.tsv",sep=' ', quote=FALSE, row.names = F,na = "NA")
```

```
#Observed
phe.obs = pcs[,c('FID','IID')]
phe.obs$pheno = maps$Observed
hist(phe.obs$pheno)
phe.obs$pheno <- rank.normalize(phe.obs[,3], FUN = qnorm)
hist(phe.obs$pheno)
phe.obs.t <- data.frame(phe.obs[,c('FID','IID','pheno')])
write.table(phe.obs.t, "GWAS_final/phenotype_files/Observed_R.tsv",sep=' ', quote=FALSE, row.names = F,na = "NA")
```

```
#InvSimpson
phe.simpson = pcs[,c('FID','IID')]
phe.simpson$pheno = maps$InvSimpson
hist(phe.simpson$pheno)
phe.simpson$FID[which(phe.simpson$pheno == max(phe.simpson$pheno))] #Aumaria, Alice without Aumaria
phe.simpson$pheno = rank.normalize(phe.simpson[,3], FUN = qnorm)
hist(phe.simpson$pheno)
phe.simpson.t <- data.frame(phe.simpson[,c('FID','IID','pheno')])
write.table(phe.simpson.t, "GWAS_final/phenotype_files/invsimpson_R.tsv", quote=FALSE, sep=' ', row.names = F,na = "NA")
```

```
#Import taxa rank.normalised abundances
gwas_taxa.df = read.csv("/GWAS_final/GWAS_plink_analysis/gwas_taxa_df_ES_T_EF.csv", sep = '\t')
rownames(gwas_taxa.df) == rownames(maps)
setdiff(rownames(gwas_taxa.df),rownames(maps))
gwas_taxa.df = gwas_taxa.df[rownames(maps),]
```

```
phe.ES = pcs[,c('FID','IID')]
phe.ES$pheno = gwas_taxa.df$Escherichia_Shigella.coli
hist(phe.ES$pheno)
phe.ES.t <- data.frame(phe.ES[,c('FID','IID','pheno')])
write.table(phe.ES.t, "GWAS_final/phenotype_files/Escherichia_Shigella_coli_rankNorm.tsv", quote=FALSE, sep=' ', row.names = F,na = "NA")

phe.TZ = pcs[,c('FID','IID')]
phe.TZ$pheno = gwas_taxa.df$Tyzzerella.unclassified
hist(phe.TZ$pheno)
phe.TZ.t <- data.frame(phe.TZ[,c('FID','IID','pheno')])
write.table(phe.TZ.t, "GWAS_final/phenotype_files/Tyzzerella_unclassified_rankNorm.tsv", quote=FALSE, sep=' ', row.names = F,na = "NA")

phe.EF = pcs[,c('FID','IID')]
phe.EF$pheno = gwas_taxa.df$Escherichia_Shigella.fergusonii
hist(phe.EF$pheno)
phe.EF.t <- data.frame(phe.EF[,c('FID','IID','pheno')])
write.table(phe.EF.t, "GWAS_final/phenotype_files/Escherichia_Shigella_fergusonii_rankNorm.tsv", quote=FALSE, sep=' ', row.names = F,na = "NA")
```

```
phe.map = maps
phe.map$pcoa = pcoa$V1
phe.map$gunifrac = gunifrac_pcoa$V1
phe.map$PC1 = pcs$PC1
phe.map$PC2 = pcs$PC2
phe.map$PC3 = pcs$PC3
phe.map$EScoli = phe.ES$pheno
phe.map$ESferg = phe.EF$pheno
phe.map$Tyz = phe.TZ$pheno

write.csv(phe.map, "kakapo_adult_map_phenotypes.csv")
```

### 5. Inbreeding correlation

```
inbred.coef = read.table("out.het", header = T)

setdiff(rownames(maps), inbred.coef$INDV)
inbred.coef$INDV[inbred.coef$INDV == "Te_Kingi"] <- "Te Kingi"
inbred.coef$INDV[inbred.coef$INDV == "Bluster_Murphy"] <- "Bluster Murphy"
inbred.coef$INDV[inbred.coef$INDV == "Tau_Kuhurangi"] <- "Tau Kuhurangi"
inbred.coef$INDV[inbred.coef$INDV == "Richard_Henry"] <- "Richard Henry"

kakapo = rownames(maps)

inbred.coef.2 = inbred.coef[inbred.coef$INDV %in% kakapo,]

cor(phe.TZ$pheno, inbred.coef.2$F, method = "pearson")
cor.test(phe.TZ$pheno, inbred.coef.2$F, method = "pearson")

cor(phe.TZ$pheno, inbred.coef.2$F, method = "spearman")
cor.test(phe.TZ$pheno, inbred.coef.2$F, method = "spearman")

plot(phe.simpson$pheno, inbred.coef.2$F)
```

### 6. GWAS with RepeatAbel in NeSI - logistic for binary traits

Example for observed phenotype:

```
#use RepeatABEL via NeSI command line: 
module load MPFR/4.0.2-GCCcore-9.2.0  --with-mpfr-include=/opt/nesi/CS400_centos7_bdw/MPFR/4.0.2-GCCcore-9.2.0/include/
module load GMP/6.2.0-GCCcore-9.2.0
module load R/4.0.1-gimkl-2020a
R
```

```
library('GenABEL')
library("RepeatABEL")
library("pracma")
library("Rmpfr")
rm(list=ls())


# produce specific genotype files (via plink)
# then transform them to a *raw datafile with the following command
convert.snp.tped(tped = "genomic_data/plink_f_nr_mic_abel.tped", 
                 tfam="genomic_data/plink_f_nr_mic_abel.tfam", 
                 out = "genomic_data/plink_f_nr_mic_abel.raw", strand = "+")


# produce phenotype files (once per phenotype)
phe <- read.table(file = "RepeatABEL/Observed_R.tsv", sep = ' ', header = TRUE)
cov <- read.table(file = "GWAS/cov.tsv", sep = ' ', header = TRUE)

phe <- subset(phe, select = -c(FID))
colnames(phe) = c('id','pheno')
phe$sex = 0
phe$sex[which(cov$Sex == 'Male')] = 1
phe$id <- as.character(phe$id)
phe$id[duplicated(phe$id)] = 'Rhenry'
phe$id <- as.factor(phe$id)

write.table(phe, file = "RepeatABEL/Observed_abel.tsv", row.names = FALSE)

# load the phenotype and genotype data in the required formats
df <- load.gwaa.data(phe = "RepeatABEL/Observed_abel.tsv", gen = "genomic_data/RepeatABEL/plink_f_nr_mic_abel.raw")

ren = rownames(df@gtdata)
ren[duplicated(rownames(df@gtdata))] <- 'Rhenry'
df@gtdata@idnames = ren

# only if binary data (since we have coded the data according to plink standards so far)
#df@phdata$pheno[df@phdata$pheno==1] = 0
#df@phdata$pheno[df@phdata$pheno==2] = 1

# simple GLS (no repeated effects)
GWAS1 <- rGLS(pheno ~ sex, genabel.data = df, phenotype.data = df@phdata)
dat = results(GWAS1)[!is.na(results(GWAS1)$P1df), ]
min(dat$P1df) # check that minimum P-value is not zero
write.csv(dat, file = "RepeatABEL/GWAS_Observed_abel.csv")
#Download data
```

### 7. Manhattan and QQ plots

Investigate where SNPs lie in kakapo genome - https://www.ncbi.nlm.nih.gov/genome/?term=txid2489341[orgn
Google significantly associated genes

```
library(qqman)
library(stringr)
library(ggplot2)
library(fdrtool)
library(jsonlite)
library(data.table)
```

```
simpson <- read.table('GWAS_final/RepeatABEL/GWAS_invsimpson_R_abel.csv', header = T, sep = ',') 
pcoa <- read.table("GWAS_final/RepeatABEL/GWAS_pcoa_abel.csv", header = T, sep = ',')
dom <- read.table("GWAS_final/RepeatABEL/GWAS_dominance_abel.csv", header = T, sep = ',')
gunifrac <- read.table("GWAS_final/RepeatABEL/GWAS_gunifrac_pcoa_abel.csv", header = T, sep = ',')
```

```
dat = dom
dat = dat[!is.na(dat$P1df), ]
dat = dat[dat$Chromosome!=3,]
dat = dat[dat$Chromosome!=13,]

dat2 = simpson
dat2 = dat2[!is.na(dat2$P1df), ]
dat2 = dat2[dat2$Chromosome!=3,]
dat2 = dat2[dat2$Chromosome!=13,]

dat3 = pcoa
dat3 = dat3[!is.na(dat3$P1df), ]
dat3 = dat3[dat3$Chromosome!=3,]
dat3 = dat3[dat3$Chromosome!=13,]

dat4 = gunifrac
dat4 = dat4[!is.na(dat4$P1df), ]
dat4 = dat4[dat4$Chromosome!=3,]
dat4 = dat4[dat4$Chromosome!=13,]

# access top SNP
min(dat$P1df) 
dat$X[which(dat$P1df == min(dat$P1df))]

top_snps <- data.frame(dat[(dat$P1df<0.001),]) 
dat2[(dat2$P1df<0.001)&(dat2$Chromosome==17),]

top_snps <- data.frame(dat[(dat.sex$P1df<0.001),]) 
top_snps2 <- data.frame(dat2[(dat2$P1df<0.001),]) 
top_snps3 <- data.frame(dat3[(dat3$P1df<0.001),]) 
top_snps4 <- data.frame(dat4[(dat4$P1df<0.001),]) 

dat.snps.5 = top_snps[which(top_snps$Chromosome==5),]
dat.snps.10 = top_snps[which(top_snps$Chromosome==10),]
dat.snps = rbind(dat.snps.5, dat.snps.10)
dat.snps = dat.snps$X

dat2.snps.5 = top_snps2[which(top_snps2$Chromosome==5),]
dat2.snps.8 = top_snps2[which(top_snps2$Chromosome==8),]
dat2.snps.17 = top_snps2[which(top_snps2$Chromosome==17),]
dat2.snps = rbind(dat2.snps.5, dat2.snps.8, dat2.snps.17)
dat2.snps = dat2.snps$X

dat3.snps.5 = top_snps3[which(top_snps3$Chromosome==5),]
dat3.snps.15 = top_snps3[which(top_snps3$Chromosome==15),]
dat3.snps.21 = top_snps3[which(top_snps3$Chromosome==21),]
dat3.snps = rbind(dat3.snps.5, dat3.snps.15, dat3.snps.21)
dat3.snps = dat3.snps$X

dat4.snps.1 = top_snps4[which(top_snps4$Chromosome==1),]
dat4.snps.2 = top_snps4[which(top_snps4$Chromosome==2),]
dat4.snps.22 = top_snps4[which(top_snps4$Chromosome==22),]
dat4.snps.77 = top_snps4[which(top_snps4$Chromosome==77),]
dat4.snps = rbind(dat4.snps.1, dat4.snps.2, dat4.snps.22, dat4.snps.77)
dat4.snps = dat4.snps$X

source("qqman_edit.R")

dir.create("Manhattan_plots")
png('Manhattan_plots/together_abel_ms.png', res=300, family="ArialMT",units='mm', width=400, height=275)
par(mfrow=c(2,2),mar=c(5, 5, 4, 2) + 0.1)
manhattan.edit(dat, chr = "Chromosome", bp = "Position", p = "P1df", snp = "X", 
          col=c("#CED38C", "#3e4e19"),
          suggestiveline = F, genomewideline = -log10(0.05/85995), annotateTop = TRUE,
          highlight = dat.snps, 
          cex=1, cex.axis=1, cex.lab=1.4, ylim = c(0, (-log10(min(dat$P1df)) + 1)), main = "ES.coli dominance")
manhattan.edit(dat2, chr = "Chromosome", bp = "Position", p = "P1df", snp = "X", 
          col=c("#CED38C", "#3e4e19"),
          suggestiveline = F, genomewideline = -log10(0.05/85995), annotateTop = TRUE,
          highlight = dat2.snps,
          cex=1, cex.axis=1, cex.lab=1.4, ylim = c(0, (-log10(min(dat2$P1df)) + 1)), main= "Inverse Simpson")
manhattan.edit(dat3, chr = "Chromosome", bp = "Position", p = "P1df", snp = "X", 
          col=c("#CED38C", "#3e4e19"),
          suggestiveline = F, genomewideline = -log10(0.05/85995), annotateTop = TRUE,
          highlight = dat3.snps,
          cex=1, cex.axis=1, cex.lab=1.4, ylim = c(0, (-log10(min(dat3$P1df)) + 1)),main= "BrayCurtis PCoA")
manhattan.edit(dat4, chr = "Chromosome", bp = "Position", p = "P1df", snp = "X", 
         col=c("#CED38C", "#3e4e19"),
          suggestiveline = F, genomewideline = -log10(0.05/85995), annotateTop = TRUE,
          highlight = dat4.snps,
          cex=1, cex.axis=1, cex.lab=1.4, ylim = c(0, (-log10(min(dat4$P1df)) + 1)),main= "gUniFrac PCoA")
dev.off()


library("GenABEL")
estlambda(dat$P1df)[1]
estlambda(dat2$P1df)[1]
estlambda(dat3$P1df)[1]
estlambda(dat4$P1df)[1]
```

```
ecoli = read.table('GWAS_final/RepeatABEL/GWAS_ES_coli_rankNorm_abel.csv', header = T, sep = ',') 
eferg = read.table('GWAS_final/RepeatABEL/GWAS_ES_fergusonii_rankNorm_abel.csv', header = T, sep = ',') 
tyzz = read.table('GWAS_final/RepeatABEL/GWAS_TZ_unclassified_rankNorm_abel.csv', header = T, sep = ',')
```

```
dat5 = ecoli
dat5 = dat5[!is.na(dat5$P1df), ]
dat5 = dat5[dat5$Chromosome!=3,]
dat5 = dat5[dat5$Chromosome!=13,]

dat6 = eferg
dat6 = dat6[!is.na(dat6$P1df), ]
dat6 = dat6[dat6$Chromosome!=3,]
dat6 = dat6[dat6$Chromosome!=13,]

dat7 = tyzz
dat7 = dat7[!is.na(dat7$P1df), ]
dat7 = dat7[dat7$Chromosome!=3,]
dat7 = dat7[dat7$Chromosome!=13,]


# access top SNP
min(dat7$P1df) 
dat7$X[which(dat7$P1df == min(dat7$P1df))]

top_snps <- data.frame(dat7[(dat7$P1df<0.001),]) #2, 6, 16, 21, 
dat5[(dat5$P1df<0.01)&(dat5$Chromosome==5),]

top_snps5 <- data.frame(dat5[(dat5$P1df<0.001),])
top_snps6 <- data.frame(dat6[(dat6$P1df<0.001),])
top_snps7 <- data.frame(dat7[(dat7$P1df<0.001),])

dat5.snps.1 = top_snps5[which(top_snps5$Chromosome==1),]
dat5.snps.5 = top_snps5[which(top_snps5$Chromosome==5),]
dat5.snps.15 = top_snps5[which(top_snps5$Chromosome==15),]
dat5.snps = rbind(dat5.snps.5, dat5.snps.15)
dat5.snps = dat5.snps$X

dat6.snps.2 = top_snps6[which(top_snps6$Chromosome==2),]
dat6.snps.6 = top_snps6[which(top_snps6$Chromosome==6),]
dat6.snps.16 = top_snps6[which(top_snps6$Chromosome==16),]
dat6.snps = rbind(dat6.snps.2, dat6.snps.6, dat6.snps.16)
dat6.snps = dat6.snps$X

dat7.snps.2 = top_snps7[which(top_snps7$Chromosome==2),]
dat7.snps.4 = top_snps7[which(top_snps7$Chromosome==4),]
dat7.snps = rbind(dat7.snps.2, dat7.snps.4)
dat7.snps = dat7.snps$X


png('Manhattan_plots/taxa_together_abel_ms.png', res=300, family="ArialMT",units='mm', width=400, height=275)
par(mfrow=c(2,2),mar=c(5, 5, 4, 2) + 0.1)
qqman::manhattan(dat5, chr = "Chromosome", bp = "Position", p = "P1df", snp = "X", 
          col=c("#CED38C", "#BCA888"),
          suggestiveline = F, genomewideline = -log10(0.05/85995), annotateTop = TRUE,
          highlight = dat5.snps, 
          cex=1, cex.axis=1, cex.lab=1.4, ylim = c(0, (-log10(min(dat$P1df)) + 1)), main = "Escherichia-Shigella coli")
qqman::manhattan(dat6, chr = "Chromosome", bp = "Position", p = "P1df", snp = "X", 
          col=c("#CED38C", "#BCA888"),
          suggestiveline = F, genomewideline = -log10(0.05/85995), annotateTop = TRUE,
          highlight = dat6.snps,
          cex=1, cex.axis=1, cex.lab=1.4, ylim = c(0, (-log10(min(dat2$P1df)) + 1)), main= "Escherichia-Shigella fergusonii")
qqman::manhattan(dat7, chr = "Chromosome", bp = "Position", p = "P1df", snp = "X", 
          col=c("#CED38C", "#BCA888"),
          suggestiveline = F, genomewideline = -log10(0.05/85995), annotateTop = TRUE,
          highlight = dat7.snps,
          cex=1, cex.axis=1, cex.lab=1.4, ylim = c(0, (-log10(min(dat3$P1df)) + 1)),main= "Tyzzerella unclassified")
dev.off()

library("GenABEL")
estlambda(dat7$P1df)[1]
estlambda(dat8$P1df)[1]
estlambda(dat9$P1df)[1]
```

```
# QQ: 
# insert correct file name and inflation factor
png('Manhattan_plots/QQ_together_abel_ms.png', family = "sans", res=300, units='mm', width = 550, height = 225)
par(mfrow=c(2,4), mar=c(5, 5, 4, 2) + 0.5)
qq(dat$P1df, main = "ES.coli dominance Genomic inflation = 0.76", 
   pch = 16, col = "#a888bc", cex=1, las = 1, cex.axis=1.8, cex.lab=1.8, cex.main=1.6,frame.plot = FALSE, adj=1)
qq(dat2$P1df, main = "Inverse Simpson Genomic inflation = 0.74", 
   pch = 16, col = "#88BCA8", cex=1, las = 1, cex.axis=1.8, cex.lab=1.8, cex.main=1.6,frame.plot = FALSE, adj=1)
qq(dat3$P1df, main = "Bray-Curtis PCoA Genomic inflation = 0.78", 
   pch = 16, col = "#7D9D33", cex=1, las = 1, cex.axis=1.8, cex.lab=1.8, cex.main=1.6,frame.plot = FALSE, adj=1)
qq(dat4$P1df, main = "gUniFrac PCoA Genomic inflation = 0.68", 
   pch = 16, col = "#BCA888", cex=1, las = 1, cex.axis=1.8, cex.lab=1.8, cex.main=1.6,frame.plot = FALSE, adj=1)
qq(dat5$P1df, main = "Escherichia-Shigella coli Genomic inflation = 0.75", 
   pch = 16, col = "#CED38C", cex=1, las = 1, cex.axis=1.8, cex.lab=1.8, cex.main=1.6,frame.plot = FALSE, adj=1)
qq(dat6$P1df, main = "Escherichia-Shigella fergusonii Genomic inflation = 1", 
   pch = 16, col = "#DCC949", cex=1, las = 1, cex.axis=1.8, cex.lab=1.8, cex.main=1.6,frame.plot = FALSE, adj=1)
qq(dat7$P1df, main = "Tyzzerella sp. Genomic inflation = 0.87", 
   pch = 16, col = "#4F7190", cex=1, las = 1, cex.axis=1.8, cex.lab=1.8, cex.main=1.6,frame.plot = FALSE, adj=1)
dev.off()
```

```
##Create SNP files for Magma 
##One for each phenotype

snp.magma <- dat[c("V3","V14")]
colnames(snp.magma) <- c("SNP","P")
dir.create("GWAS_final/Magma")
write.table(snp.magma,"GWAS_final/Magma/SNP_files/GWAS/SNP_file_dom_abel.tsv",sep=' ', quote=FALSE, row.names = F)
```

### 7. Gene enrichment pathway analysis - Magma

Edit the gmt files with ENSEMBL gene IDs to have gene symbols
instead

```
library(dplyr)
library(tidyverse)
library('biomaRt')
ensembl <- useMart("ensembl")
datasets <- listDatasets(ensembl)


######Chicken######
martgg <- useDataset("ggallus_gene_ensembl", useMart("ensembl"))

gg <- read.csv("GWAS_final/Magma/gmt_databses/Gallus_gallus_GO_KEGG_ensembl_gene_id.gmt.gz", sep = "\t", header = F)
gene.list <- gg[,3:495]
#Sys.setenv("http_proxy" = "http://my.proxy.org:9999")
G_list <- getBM(filters= "ensembl_gene_id", attributes= c("ensembl_gene_id","hgnc_symbol"),values=gene.list,mart= martgg)
write.table(G_list$ensembl_gene_id, "GWAS_final/Magma/gg_ensembleIDs.csv", sep = '\t', row.names = F)


##read in new table where ensembl IDs were uploaded onto the online database to get more gene symbol hits
gg_geneS <- read.table("GWAS_final/Magma/gg_gene_symbols.txt", header = T, sep = '\t')
gg2 = gg
cols <- 3:495
gg2[cols] <- lapply(gg2[cols], function(x) 
  gg_geneS$gene_symbol[match(x, gg_geneS$gg_ensemblID)])
gg2[is.na(gg2)] = ''

##Filter gmt file to only have genes identified in kakapo genome annotation file (downloaded from NeSI)
annot <- read.table("GWAS_final/Magma/gmt_databses/filtered_mic_annotation.txt", header = F, sep = '\t')
gg2[cols] <- lapply(gg2[cols], function(x) 
  annot$V1[match(x, annot$V1)])

##Need to delete the rows where there are now no genes left > change all the empty cells to NA
gg2 <- gg2 %>% 
  mutate(across(everything(), ~ifelse(.=="", NA, as.character(.))))
gg2 <- gg2[!(is.na(gg2[,3:495])),]
#Remove duplicated geneset pathways
gg3 <- gg2[!duplicated(gg2$V1),,drop=FALSE]
#remove NAs for final gmt file
gg2[is.na(gg2)] = ''

write.table(gg3,"GWAS_final/Magma/Gallus_gallus_genes_filt.gmt", row.names = F,sep = ' ', quote = F, col.names = F)


######Human######
marths <- useDataset("hsapiens_gene_ensembl", useMart("ensembl"))
hs <- read.csv("GWAS_final/Magma/gmt_databses/Homo_sapiens_GO_KEGG_ensembl_gene_id.gmt", sep = '\t', header = F)
hs.list <- hs[,3:1356]
Ghs_list <- getBM(filters= "ensembl_gene_id", attributes= c("ensembl_gene_id","hgnc_symbol"),values=hs.list,mart= marths)

col3 <- 3:1356
hs[col3] <- lapply(hs[col3], function(x)
  Ghs_list$hgnc_symbol[match(x, Ghs_list$ensembl_gene_id)])
hs[is.na(hs)] = ''

hs[col3] <- lapply(hs[col3], function(x) 
  annot$V1[match(x, annot$V1)])

hs <- hs %>% 
  mutate(across(everything(), ~ifelse(.=="", NA, as.character(.))))
hs <- hs[!(is.na(hs[,3:1356])),]
#Remove duplicated geneset pathways
hs <- hs[!duplicated(hs$V1),,drop=FALSE]
#remove NAs for final gmt file
hs[is.na(hs)] = ''

write.table(hs,"GWAS_final/Magma/Homo_sapiens_genes.gmt", row.names = F,sep = ' ', quote = F, col.names = F)
```

```
GO_db <- read.csv("GWAS_final/Magma/gmt_databases/Human_GOALL_with_GO_iea_March_01_2021_symbol.gmt", header = F, sep = '\t')

GO_db[,1:2] <- lapply(GO_db[,1:2], function(x) gsub(" ", "_", x))

GO_db2 <- GO_db[!apply(GO_db[,3:231] == "", 1, all), ] 

#Remove duplicated geneset pathways

GO_db3 = GO_db2 %>% distinct(GO_db2$V1, .keep_all = TRUE)
GO_db3 = GO_db3 %>% distinct(GO_db3$V2, .keep_all = TRUE)

write.table(GO_db3,"GWAS_final/Magma/Human_GOALL_filt.gmt", row.names = F,sep = ' ', quote = F, col.names = F)
```

Upload gmt and SNP files to NeSI and run magma.sl and magma\_db.sl
scripts.

```
#Run (example): 
awk '{ print $(NF-1), $NF}' Tyzzerella_abel.gsa.out > reformatted_output_files/Tyzzerella_abel.out to get output for following R analyses
```

Investigate significant gene pathways

```
### repeatABEL with Chicken pathways###

##PCOA##
pcoa_RA <- read.table("GWAS_final/Magma/output_files/pcoa_RA.out",sep = " ")
pcoa_RA = pcoa_RA[-c(1:4),]
names(pcoa_RA) <- pcoa_RA[1,]
pcoa_RA = pcoa_RA[-1,]

dat2 <- data.frame(pcoa_RA)
dat2[1] <- lapply(dat2[1], p.adjust, method="bonferroni")
dat2[(dat2$P<0.05),]


##E.COLI DOMINANCE##
dom_RA <- read.table("GWAS_final/Magma/output_files/dom_RA.out",sep = " ")
dom_RA = dom_RA[-c(1:4),]
names(dom_RA) <- dom_RA[1,]
dom_RA = dom_RA[-1,]

dat2 <- data.frame(dom_RA)
dat2[1] <- lapply(dat2[1], p.adjust, method="bonferroni")
dat2[(dat2$P<0.05),]


##INVERSE SIMPSON##
invsimpR_RA <- read.table("GWAS_final/Magma/output_files/repeatABEL/invsimpR_RA.out",sep = " ")
invsimpR_RA = invsimpR_RA[-c(1:4),]
names(invsimpR_RA) <- invsimpR_RA[1,]
invsimpR_RA = invsimpR_RA[-1,]

dat2 <- data.frame(invsimpR_RA)
dat2[1] <- lapply(dat2[1], p.adjust, method="bonferroni")
dat2[(dat2$P<0.05),]


##GUNIFRAC##
gunifrac_RA <- read.table("GWAS_final/Magma/output_files/repeatABEL/gunifrac_RA.out",sep = " ")
gunifrac_RA = gunifrac_RA[-c(1:4),]
names(gunifrac_RA) <- gunifrac_RA[1,]
gunifrac_RA = gunifrac_RA[-1,]

dat2 <- data.frame(gunifrac_RA)
dat2[1] <- lapply(dat2[1], p.adjust, method="bonferroni")
dat2[(dat2$P<0.05),]


##E.COLI##
Ecoli_RA <- read.table("GWAS_final/Magma/output_files/repeatABEL/Ecoli_abel.out",sep = " ")
Ecoli_RA = Ecoli_RA[-c(1:4),]
names(Ecoli_RA) <- Ecoli_RA[1,]
Ecoli_RA = Ecoli_RA[-1,]

dat2 <- data.frame(Ecoli_RA)
dat2[1] <- lapply(dat2[1], p.adjust, method="bonferroni")
dat2[(dat2$P<0.05),]


##E.FERGUSONII##
Eferg_RA <- read.table("GWAS_final/Magma/output_files/repeatABEL/Eferg_abel.out",sep = " ")
Eferg_RA = Eferg_RA[-c(1:4),]
names(Eferg_RA) <- Eferg_RA[1,]
Eferg_RA = Eferg_RA[-1,]

dat2 <- data.frame(Eferg_RA)
dat2[1] <- lapply(dat2[1], p.adjust, method="bonferroni")
dat2[(dat2$P<0.05),]


##TYZZERELLA##
Tyzz_RA <- read.table("GWAS_final/Magma/output_files/repeatABEL/Tyzzerella_abel.out",sep = " ")
Tyzz_RA = Tyzz_RA[-c(1:4),]
names(Tyzz_RA) <- Tyzz_RA[1,]
Tyzz_RA = Tyzz_RA[-1,]

dat2 <- data.frame(Tyzz_RA)
dat2[1] <- lapply(dat2[1], p.adjust, method="bonferroni")
dat2[(dat2$P<0.05),]
```

### 8. BayesR Heritibility

Copy plink\_f\_nr\_mic files to new folder in BayesR folder. Make named
copies for each phenotype. Run awk ‘FNR==NR{a[NR]=$3;next}{$6=a[FNR]}1’
<(tail -133 ../RepeatABEL/Tyzzerella\_unclassified\_rankNorm.tsv)
./plink/plink\_f\_nr\_tyzz\_mic.fam | cat >
./plink/plink\_f\_nr\_tyzz\_mic.fam2 Check fam2 file is correct, then change
to regular .fam mv plink/plink\_f\_nr\_tyzz\_mic.fam2
plink/plink\_f\_nr\_tyzz\_mic.fam

Run bayesR.sl scripts. For phenotypes that are autocorrelated,
increase number of chains: -numit 200000 -burnin 50000 -thin 100 >
get 1500 data points back

Example analysis for one phenotype:

```
library(lme4)
library(ggplot2)
library(scales)
library(coda)
library(dplyr)
library(tidyr)
library(qqman)
library(RColorBrewer)
library(hrbrthemes)
library(extrafont)

prefix<-"Y:/GWAS_final/BayesR_plots/bayesR_Eferg/Eferg_abel" ### This is the prefix of the BayesR output files
famFile<-"Y:/NeSI_files_final/BayesR/plink/plink_f_nr_pcoa_mic.fam" 
bimFile<-"Y:/NeSI_files_final/BayesR/plink/plink_f_nr_pcoa_mic.bim"


# get point estimates

gvFile <-paste(prefix,".gv",sep="")
paramFile <- paste(prefix,".param",sep="")
modelFile<-paste(prefix,".model",sep="")
hypFile <-paste(prefix,".hyp",sep="")
predicted_gvFile<-paste(prefix,".gv",sep="") ### GEBVs of training population
frqFile<-paste(prefix,".frq",sep="")

ModelSummary <- read.table(modelFile,sep="",head=F,row.names=1) 
ModelSummary$V2 <- as.character(ModelSummary$V2)

# run for Vk1 and Vk4 >> look for these individually
ModelSummary['Vk1',] <- gsub("\\-", "E+", ModelSummary['Vk1',]) #run this (0) and 2 for dom
ModelSummary['Vk1',] <- gsub("\\+", "E+", ModelSummary['Vk1',]) #run 1 and 2 for pcoa #run 1 for Eferg
ModelSummary['Vk2',] <- gsub("\\+", "E+", ModelSummary['Vk2',]) #run 2 3 and 4 for gunifrac  #just 2 for Ecoli
ModelSummary['Vk3',] <- gsub("\\+", "E+", ModelSummary['Vk3',]) #only run this for invsimpR
ModelSummary['Vk4',] <- gsub("\\+", "E+", ModelSummary['Vk4',]) #run all 4 for Tyzz

ModelSummary$V2 <- as.numeric(ModelSummary$V2)

(N_SNPs <- ModelSummary["Nsnp",])
(Va <- ModelSummary["Va",])
(Ve <- ModelSummary["Ve",])
(Herit <- ModelSummary["Va",]/sum(ModelSummary[3:4,]))

(N_SNPs_0 <- ModelSummary["Nk1",])
(N_SNPs_0.0001 <- ModelSummary["Nk2",])
(N_SNPs_0.001 <- ModelSummary["Nk3",])
(N_SNPs_0.01 <- ModelSummary["Nk4",])
(PVE_0.0001 <- ModelSummary["Vk2",]/sum(ModelSummary[3:4,]))
(PVE_0.001 <- ModelSummary["Vk3",]/sum(ModelSummary[3:4,]))
(PVE_0.01 <- ModelSummary["Vk4",]/sum(ModelSummary[3:4,]))
(PGE_0.0001 <- ModelSummary["Vk2",]/ModelSummary["Va",])
(PGE_0.001 <- ModelSummary["Vk3",]/ModelSummary["Va",])
(PGE_0.01 <- ModelSummary["Vk4",]/ModelSummary["Va",])
```

```
# get confidence intervals

ModelPostParams <- read.table(hypFile,sep="",head=T)

#run both for dom pcoa gunifrac Ecoli Eferg
#don't run this for invsimpR or Tyzz
for (i in c("Va", "Vk2", "Vk3", "Vk4")) {
  print(i)
  ModelPostParams[i] <- gsub("-", "E-", unlist(ModelPostParams[i]))
  ModelPostParams[i] <- gsub("EE", "E", unlist(ModelPostParams[i]))
  ModelPostParams[i] <- as.numeric(unlist(ModelPostParams[i]))
}

for (i in colnames(ModelPostParams)){
  ModelPostParams[i] = as.numeric(as.character(unlist(ModelPostParams[i])))
  print(i) # here: Va, Vk2, Vk3, Vk4
}


ModelPostParams$Herit <- ModelPostParams$Va/(ModelPostParams$Va + ModelPostParams$Ve)
ModelPostParams$PGE_0.0001 <- ModelPostParams$Vk2/(ModelPostParams$Vk2 + ModelPostParams$Vk3 + ModelPostParams$Vk4)
ModelPostParams$PGE_0.001 <- ModelPostParams$Vk3/(ModelPostParams$Vk2 + ModelPostParams$Vk3 + ModelPostParams$Vk4)
ModelPostParams$PGE_0.01 <- ModelPostParams$Vk4/(ModelPostParams$Vk2 + ModelPostParams$Vk3 + ModelPostParams$Vk4)


(Va_95 <- quantile(ModelPostParams$Va, c(0.025, 0.975)))
(Ve_95 <- quantile(ModelPostParams$Ve, c(0.025, 0.975)))
(N_SNPs_95 <- quantile(ModelPostParams$Nsnp, c(0.025, 0.975)))
(Herit_95 <- quantile((ModelPostParams$Va/(ModelPostParams$Va + ModelPostParams$Ve)), c(0.025, 0.975)))
(N_SNPs_0_95 <- quantile(ModelPostParams$Nk1, c(0.025,0.975)))
(N_SNPs_0.0001_95 <- quantile(ModelPostParams$Nk2, c(0.025,0.975)))
(N_SNPs_0.001_95 <- quantile(ModelPostParams$Nk3, c(0.025,0.975)))
(N_SNPs_0.01_95 <- quantile(ModelPostParams$Nk4, c(0.025,0.975)))###########################
(PGE_0.0001_95 <- quantile(ModelPostParams$PGE_0.0001, c(0.025,0.975)))
(PGE_0.001_95 <- quantile(ModelPostParams$PGE_0.001, c(0.025,0.975)))
(PGE_0.01_95 <- quantile(ModelPostParams$PGE_0.01, c(0.025,0.975)))

median(ModelPostParams$Va/(ModelPostParams$Va + ModelPostParams$Ve))
mean(ModelPostParams$Va/(ModelPostParams$Va + ModelPostParams$Ve))
#Tyzerella = #4F7190
#ES ferg = #F7EE55
#ES coli = #CAE0AB
#dom = #a888bc
#inv simp = #88BCA8
#bray pcoa = #7D9D33
#gunifrac = #BCA888

library(ggplot2)
library(ggtext)

eferg.hert.hist = ggplot(ModelPostParams, aes(x = Herit)) + 
    geom_density(fill = "#F7EE55") + 
    xlab(bquote("h"^2)) +
    ylab("Density") +
    theme_ipsum() + 
    theme(axis.line.x=element_line(color="black",size=1.0,linetype=1), axis.line.y=element_line(color="black",size=1.0,linetype=1),
                        panel.grid.major = element_blank(), panel.grid.minor = element_blank(), panel.background = element_blank()) +
    ggtitle("*Escherichia-Shigella fergusonii*") +
    theme(axis.title.y=element_text(size=28), axis.title.x=element_text(size=28), 
        axis.text.x=element_text(size=26), axis.text.y=element_text(size=26), plot.title = element_markdown(size=30))

eferg.hert.hist
```

```
# look at autocorrelation 
ggplot(ModelPostParams, aes(x=Replicate, y=Nsnp))+geom_line()
ggplot(ModelPostParams, aes(x=Replicate, y=Va))+geom_line()
ggplot(ModelPostParams, aes(x=Replicate, y=Ve))+geom_line()
ggplot(ModelPostParams, aes(x=Replicate, y=Nk1))+geom_line()
ggplot(ModelPostParams, aes(x=Replicate, y=Herit))+geom_line()

# BayesR, plot MCMC chains:
# insert correct file name
eferg.mcmc = ggplot(ModelPostParams, aes(x=Replicate, y=Herit))+geom_line(colour="#F7EE55")+
  theme_ipsum() + theme(axis.line.x=element_line(color="black",size=1.0,linetype=1), axis.line.y=element_line(color="black",size=1.0,linetype=1),
                        panel.grid.major = element_blank(), panel.grid.minor = element_blank(), panel.background = element_blank()) +
  ylab("Heritability\n") + 
  xlab("\nMCMC sample") + ggtitle("*Escherichia-Shigella fergusonii*") +
  theme(axis.title.y=element_text(size=17), axis.title.x=element_text(size=17), 
        axis.text.x=element_text(size=15), axis.text.y=element_text(size=14), plot.title = element_markdown(size = 17)) 
eferg.mcmc

# ideally <0.1
autocorr(as.mcmc(ModelPostParams$Nsnp))
autocorr(as.mcmc(ModelPostParams$Va))
autocorr(as.mcmc(ModelPostParams$Ve))
autocorr(as.mcmc(ModelPostParams$Herit))
```

```
# load SNp effects

ModelSNPs <- read.table(paramFile,sep="",head=T)

for (i in colnames(ModelSNPs)){
  ModelSNPs[i] = as.numeric(as.character(unlist(ModelSNPs[i])))
  print(i) 
}
dels = which(is.na(unlist(ModelSNPs['beta'])))

dels = array(which(is.na(unlist(ModelSNPs['beta']))))
ModelSNPs = ModelSNPs[complete.cases(ModelSNPs), ]
ModelSNPs['beta'] <- as.numeric(unlist(ModelSNPs['beta']))

ModelSNPs$rank <- rank(abs(ModelSNPs$beta))
```

```
# chromosome partitioning

SNP_Info <- read.table(bimFile,sep="",header=F)
SNP_Info <- SNP_Info[-dels,] #for eferg only
colnames(SNP_Info) <- c("Chrom","SNP", "cM", "BP","REF", "ALT")
SNPs <- cbind(SNP_Info[,c(1,2,4)], ModelSNPs)
head(SNPs)

# allele Freqs
SNP_Freq <- read.table(frqFile,sep="",header=F)
SNP_Freq <- SNP_Freq[-dels,] #for eferg only
SNP_Freq = data.frame(SNP_Freq)
head(SNP_Freq)
colnames(SNP_Freq) <- c("Freq")
SNPs <- cbind(SNPs,SNP_Freq)
head(SNPs)

# estimate SNP effect as 2p(1 - p)*beta^2
SNPs['V'] <- 2 * SNPs$Freq * (1 - SNPs$Freq) * abs(SNPs$beta)^2

# showing top 20 SNPs 
SNPs[order(SNPs$V, decreasing = TRUE),][1:20,]

## get variance
GenomVar <- sum(SNPs$V)
ChromVar <- aggregate(V ~ Chrom,data=SNPs,sum)
head(ChromVar)
ChromVar$PVa <- ChromVar$V/GenomVar
sum(ChromVar$PVa)


ChromVar$Size <- tapply(SNPs$BP, SNPs$Chrom, max)
```

```
# Plot amount of Va explained by each chromosome

# label the biggest and those that explain most Va via ChromLabel
idChrom <- ChromVar[(ChromVar$PVa  >= 0.03 | ChromVar$Size >= 60000000),"Chrom"] 
ChromVar$ChromLabel <- ChromVar$Chrom
ChromVar$ChromLabel[!(ChromVar$ChromLabel %in% idChrom)] <- NA

ylim_max <- max(ChromVar$PVa)+0.01

# BayesR, plot chromosome partitioning:
eferg.cp = ggplot(ChromVar, aes(x=Size/1000000, y=PVa))+geom_point(size=6,shape=21,colour="black",fill="#F7EE55",alpha=0.8) +
  theme_ipsum() + theme(axis.line.x=element_line(color="black",size=1.0,linetype=1), axis.line.y=element_line(color="black",size=1.0,linetype=1),
                        panel.grid.major = element_blank(), panel.grid.minor = element_blank(), panel.background = element_blank()) +
  xlim (0, max(ChromVar$Size)/1000000) + ylim (0,ylim_max) +
  geom_text(aes(label=ChromLabel),size=6,hjust=-0.9) +
  ylab ("Variance Explained\n") + 
  xlab("\nChromosome Size (Mb)") + ggtitle("*Escherichia-Shigella fergusonii*") +
  theme(axis.title.y=element_text(size=17), axis.title.x=element_text(size=17), 
        axis.text.x=element_text(size=15), axis.text.y=element_text(size=14), axis.title = element_text(size = 17), 
        plot.title = element_markdown(hjust = 0.5)) 
eferg.cp
```

#9. Plots

```
library(cowplot)

cp.alpha = plot_grid(dom.cp, invsimp.cp, pcoa.cp, gunifrac.cp, labels = c("A","B","C","D"), label_size = 30, ncol=4)
cp.alpha
ggsave("CP_alpha_plots.png", units='mm', width=750, height=175, bg = "white", dpi=300)

cp.taxa = plot_grid(ecoli.cp,eferg.cp,tyzz.cp, labels = c("A","B","C"), label_size = 30, ncol = 3)
cp.taxa
ggsave("CP_taxa_plots.png", units='mm', width=500, height=175, bg = "white", dpi=300)


mcmc.together = plot_grid(dom.mcmc, invsimp.mcmc, pcoa.mcmc, gunifrac.mcmc, ecoli.mcmc, eferg.mcmc, tyzz.mcmc, labels = c("A","B","C","D","E","F","G"), label_size = 30, ncol=4)
mcmc.together
ggsave("MCMC_together_plots.png", units='mm', width=750, height=350, bg = "white", dpi=300)


hist.post.param = plot_grid(dom.hert.hist, simp.hert.hist, pcoa.hert.hist, gunifrac.hert.hist, ecoli.hert.hist, eferg.hert.hist, tyzz.hert.hist, 
                            labels = c("A","B","C","D","E","F","G"), label_size = 30, ncol=4)
hist.post.param
ggsave("Histpostparam_together_plots.png", units='mm', width=750, height=400, bg = "white", dpi=300)
```

```
png('Manhattan_plots_alpha_abel.png', res=300, family="sans",units='mm', width=700, height=150, bg = "white")
par(mfrow=c(1,4),mar=c(5, 8, 4, 2) + 0.1)
manhattan.edit(dat, chr = "Chromosome", bp = "Position", p = "P1df", snp = "X", 
          col=c("#CED38C", "#3e4e19"),
          suggestiveline = F, genomewideline = -log10(0.05/85995), annotateTop = TRUE,
          highlight = dat.snps, 
          cex=1, cex.axis=1.4, cex.lab=2, ylim = c(0, (-log10(min(dat$P1df)) + 1)), adj=1)
manhattan.edit(dat2, chr = "Chromosome", bp = "Position", p = "P1df", snp = "X", 
          col=c("#CED38C", "#3e4e19"),
          suggestiveline = F, genomewideline = -log10(0.05/85995), annotateTop = TRUE,
          highlight = dat2.snps,
          cex=1, cex.axis=1.4, cex.lab=2,cex.ylim = c(0, (-log10(min(dat2$P1df)) + 1)), adj=1)
manhattan.edit(dat3, chr = "Chromosome", bp = "Position", p = "P1df", snp = "X", 
          col=c("#CED38C", "#3e4e19"),
          suggestiveline = F, genomewideline = -log10(0.05/85995), annotateTop = TRUE,
          highlight = dat3.snps,
          cex=1, cex.axis=1.4, cex.lab=2,ylim = c(0, (-log10(min(dat3$P1df)) + 1)), adj=1)
manhattan.edit(dat4, chr = "Chromosome", bp = "Position", p = "P1df", snp = "X", 
         col=c("#CED38C", "#3e4e19"),
          suggestiveline = F, genomewideline = -log10(0.05/85995), annotateTop = TRUE,
          highlight = dat4.snps,
          cex=1, cex.axis=1.4, cex.lab=2., ylim = c(0, (-log10(min(dat4$P1df)) + 1)), adj=1)
dev.off()
```

```
png('Manhattan_plots_taxa_abel.png', res=300, family="sans",units='mm', width=550, height=150)
par(mfrow=c(1,3),mar=c(5, 8, 4, 2) + 0.1)
manhattan.edit(dat5, chr = "Chromosome", bp = "Position", p = "P1df", snp = "X", 
          col=c("#CED38C", "#3e4e19"),
          suggestiveline = F, genomewideline = -log10(0.05/85995), annotateTop = TRUE,
          highlight = dat5.snps, 
          cex=1, cex.axis=1.4, cex.lab=2, ylim = c(0, (-log10(min(dat$P1df)) + 1)), adj=1)
manhattan.edit(dat6, chr = "Chromosome", bp = "Position", p = "P1df", snp = "X", 
          col=c("#CED38C", "#3e4e19"),
          suggestiveline = F, genomewideline = -log10(0.05/85995), annotateTop = TRUE,
          highlight = dat6.snps,
          cex=1, cex.axis=1.4, cex.lab=2, ylim = c(0, (-log10(min(dat2$P1df)) + 1)), adj=1)
manhattan.edit(dat7, chr = "Chromosome", bp = "Position", p = "P1df", snp = "X", 
          col=c("#CED38C", "#3e4e19"),
          suggestiveline = F, genomewideline = -log10(0.05/85995), annotateTop = TRUE,
          highlight = dat7.snps,
          cex=1, cex.axis=1.4, cex.lab=2,ylim = c(0, (-log10(min(dat3$P1df)) + 1)), adj=1)
dev.off()
```
